# Overcoming Pitfalls when Estimating Variation in Parental Care: A Simulation Study

**DOI:** 10.1101/2023.10.16.562548

**Authors:** R. Mundry, F. Dal Pesco, A. Avilès de Diego, J. Fischer

## Abstract

**Background:** In many animal species, parents allocate resources to enhance offspring survival and thereby reproductive success (‘parental investment’). Parental care behaviors are a crucial part of parental investment, and one critical question in behavioral ecology is to which degree variation in parental care affects offspring development and survival. Crucially, parental care behaviors vary with offspring age, which creates non-trivial problems for statistical analyses, specifically when sampling is incomplete or biased.

**Methods:** We conducted a simulation study to illustrate the problem when analyzing average parental care behaviors while not correcting for offspring age (‘conventional approach’), and propose a modeling approach to correct for offspring age and sampling biases. We simulated a set of different scenarios, varying with regard to sample size, sampling scheme, and variation in parental behavior. To model parent behavior as a function of offspring age, we fitted a Generalized Linear Mixed Model to each simulated data set. We included offspring age as a fixed effect and extracted Best Linear Unbiased Predictors (BLUPs) for the random intercept of parent identity. These BLUPs represent the parental behavior corrected for offspring age. Finally, we assessed how our proposed modeling approach compares to the conventional approach.

**Results:** The proposed modeling approach was superior to the conventional approach. More specifically, the conventional approach clearly overestimated the variation between parents, and even diagnosed appreciable variation when there was none in the simulated data. This bias was exacerbated in incomplete or unbalanced sampling schemes.

**Conclusion.:** Our simulation study clearly demonstrates the necessity of correcting for offspring age in analyses of parental behavior. We strongly suggest including random slopes of offspring age within parent-offspring dyads, because doing so can reveal parent-specific trajectories of the modeled behavior against offspring age. Our approach allows for the inclusion of further categorical and continuous predictor variables that may affect parental behavior, such as parity or ecological conditions. In summary, the proposed modeling approach has several advantages over conventional methods.

## Introduction

In many animal species, parents allocate resources to their offspring to enhance offspring survival and reproductive success at the cost of their future reproduction. This resource allocation is known as ‘parental investment’ (Trivers, 1972). Parental investment theory conceives the investment as a continuous process, starting at conception, with a gradual shift in the return of investment as the offspring matures (Trivers, 1972), resulting in an increasing conflict between parent and offspring over the allocation of resources (Trivers, 1974). Parental investment theory seeks to explain the variation in the extent and nature of parental investment between species and individuals. Factors affecting variation in parental investment between individuals include the certainty of parent-offspring relatedness, the potential for future reproduction, and the availability and distribution of resources (Davies et al., 2012).

Investments that are provided directly to the offspring are known as parental care (Smiseth et al., 2012). Well-known forms of parental care include the incubation of eggs in birds (Porras-Reyes et al., 2021; Da et al., 2018), nest- and egg guarding in fish (Lindström, St. Mary, Pampoulie, 2006), and insects provisioning food for larvae (Capodeanu-Nägler et al., 2018). In primates, care for infants is particularly extensive. In addition to female nursing – as observed in all mammals – infants are carried and protected for prolonged times (Altmann, 1980; van Noordwijk, 2012). Evidently, parental care varies with offspring age (e.g., Hinde & Spencer-Booth, 1968; Nash, 1978). Any estimation of variation in parental care across species or individuals must therefore take offspring age into account.

The age-dependent variation in parental care poses the question of how to precisely estimate the extent of variation in parental contribution. For instance, if we were interested in how the time spent feeding young varies with parent sex, the variable ‘time spent provisioning’ will vary with offspring age. This link between the variable observed (time spent provisioning) and offspring age can create substantial problems, particularly if the age at which offspring are observed varies between parent-offspring dyads. In such cases, when offspring age is not taken into account, one would get biased estimates of parental behavior. More specifically, variation between parents will be confounded with variation related to the age at which offspring were observed (Fig. 1). Although the issues arising from the failure to take offspring age into account are evident, there are several examples where the authors did not consider variation in offspring age in their analyses (e.g., Fairbanks & McGuire, 1987; Schino, D’Amato, Troisi, 1995; Bardi & Huffman, 2002; De Lathouwers & Van Elsacker, 2004; but see Dura et al., 2018).

**Fig. 1.**
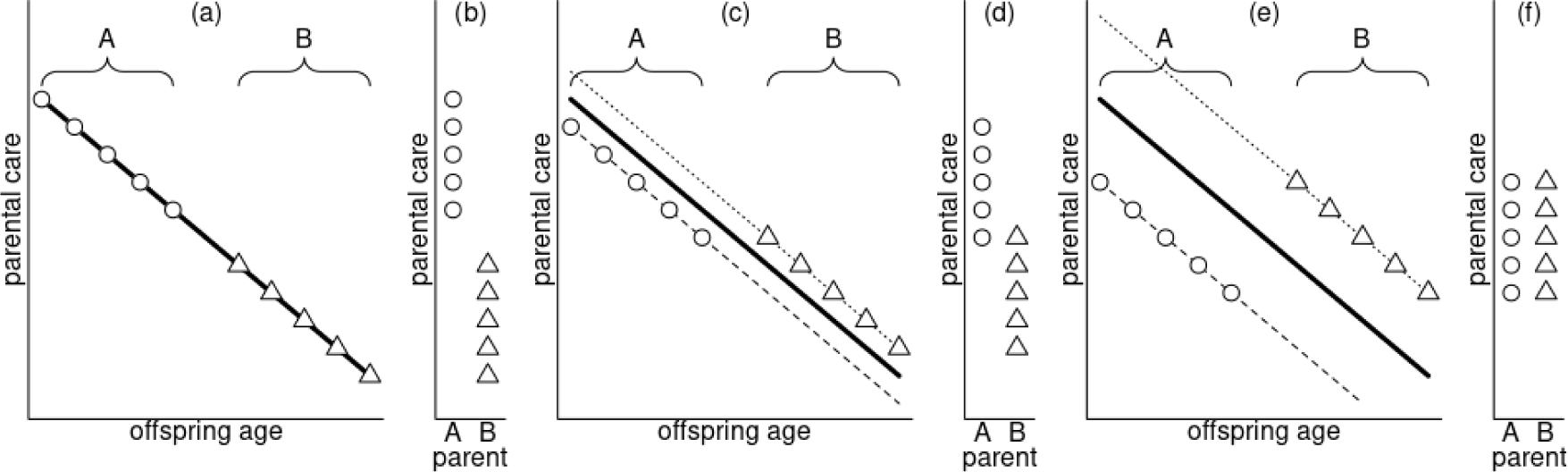
Illustration of the effects of unbalanced sampling on the estimation of variation between parents (depicted by different symbols) with regard to their propensity to show parental care. Note that the two parents differ with regard to when they were observed during offspring development. The thick line represents the average parental care. In (a), two parents do not differ in their parental care, but parent A, depicted by round symbols, will be assessed to have shown a higher parental care than parent B, represented by triangles, if one ignores offspring age (b). In (c), the two parents differ in parental care, whereby parent A performs less than average parental care, and parent B higher than average parental care. However, if one would just average parental care irrespective of offspring age, one would come to the opposite conclusion (d). In (e), the difference in parental care follows the same pattern as in (c) but with a greater magnitude. This time, averaging parental care irrespective of offspring age would lead to the conclusion that both parents perform the behavior to the same extent (f).

Here we propose a novel modeling approach that solves this problem by modeling parental care as a function of offspring age and the identity of the parent-offspring dyad. In brief, we use a (Generalized) Linear Mixed Model with the parental behavior of interest as the response, offspring age as a fixed effect, parent-offspring dyad as a random intercept, and offspring age as a random slope. We then extract Best Linear Unbiased Predictors (BLUPs; Baayen 2008) for the random intercept of parent-offspring dyad. These BLUPs are then used as estimates of parental investment that are corrected for offspring age. The BLUPs provide information on the extent to which individual parental investment is below or above the average parental investment for a given age of the offspring.

We present a simulation study to illustrate the problem when using average parental care while not correcting for offspring age and investigated to which extent the use of the average investment (‘conventional’ method hereafter) might bias the assessment of variation in parental care. To this end, we simulated different sampling schemes and different levels of stochastic variation in the individual observations (see methods for details). Finally, we examined whether and to which degree our proposed modeling approach alleviates the above-mentioned problems.

## Methods

### Simulation settings

We conducted a simulation study to evaluate the performance of the ‘conventional’ and our proposed ‘modeling’ method. We assumed uni-parental care and simulated a parental behavior in which a parent shows this behavior towards their offspring 95% of the time after birth/hatching and 5% of the time at the end of the considered period of offspring development, here set to have a duration of 1 (Fig. 2, thick dashed line). We simulated two scenarios concerning the variation between parents: in the first scenario (null scenario), all parents showed precisely the same amount of parental care. In the second scenario, parents differed in their care (variable parental care scenario, Fig. 2, thin dashed lines). The null scenario allows us to assess to what extent the conventional and the modeling approach give rise to “false positives”; that is, whether the results would indicate that parents differ in their care when they do not. The variable parental care scenario, in turn, allows us to asses to which extent the two approaches retrieve the simulated parent-specific propensities to display the behavior.

**Fig. 2.**
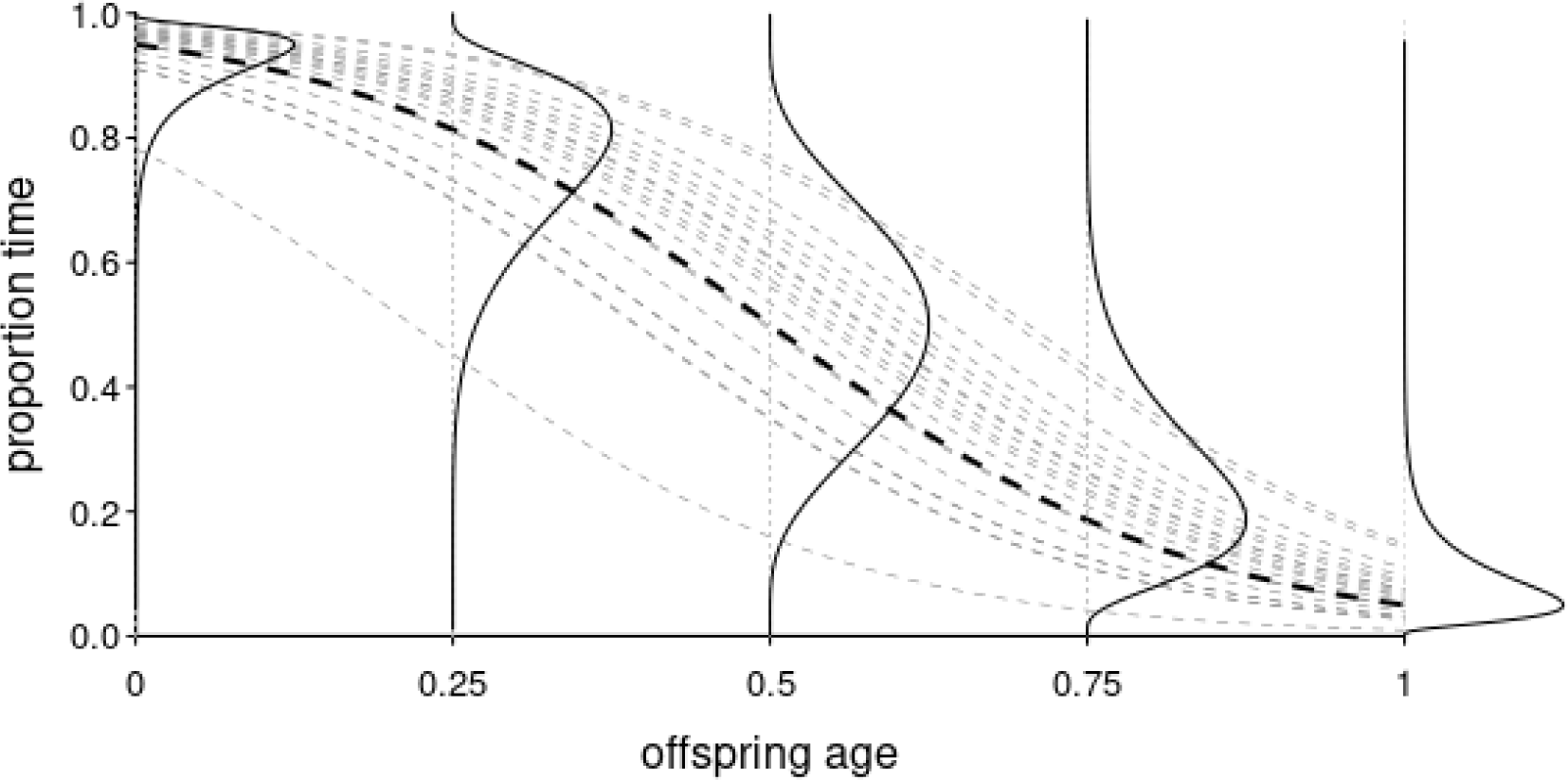
Simulated average parental behavior and variation between parents with regard to their propensity to perform the behavior. The thick dashed line depicts the average parent as simulated. Solid lines depict the magnitude of simulated variation among parents, of which the thin dashed lines illustrate an example of 20 randomly chosen parents. The magnitude of random variation between parents depicted arose from assigning parents a random intercept effect sampled from a normal distribution with a mean of zero and a standard deviation of 0.75.

Furthermore, we simulated three different sampling schemes. In the first, each parent-offspring dyad was observed over the entire period. The offspring ages at which dyads were observed were identical for all parent-offspring dyads (Fig. 3a). In the second scenario, the ages at which parent-offspring dyads were observed were randomly sampled during the observation period (Fig. 3b). Lastly, we simulated a scenario in which each parent-offspring dyad was observed for half of the total length of the considered period of offspring development (Fig. 3c). In each of the three scenarios, the number of observations per parent-offspring dyad was identical for each dyad. We further simulated two sample sizes with regard to the number of parent-offspring dyads (N = 20 or 80 dyads) and two sample sizes for the number of observations per parent-offspring dyad (N = 21 or 81 observations). Finally, we simulated different magnitudes of stochastic variation of the actual observations (precision parameter, see below).

**Fig. 3.**
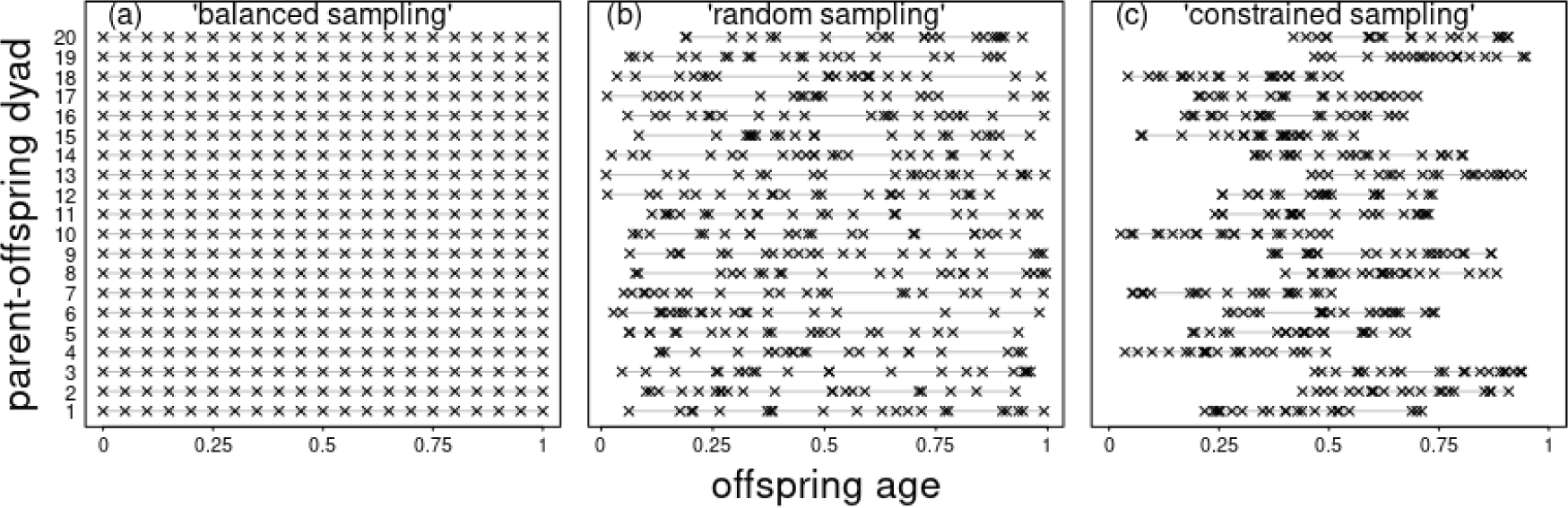
Illustration of the three different observation schemes we simulated. Each line represents one parent-offspring dyad. Laying crosses depict days at which observations take place. (a) Observation days take place at the exact same offspring ages across dyads; (b) observation days are randomly distributed within the considered period of offspring development; (c) observation days are randomly distributed within the considered period of offspring development, but for each offspring, over a time span of at most half that period.

For the random sampling scenario (Fig. 3b), we sampled the age from a uniform distribution between the minimum and maximum age per offspring. To model a scenario where offspring are observed for only a part of the period under consideration (‘constrained’ sampling; Fig. 3c), we randomly sampled the onset of the observation period per offspring from the first half of the observation period and the remaining observations from an interval between the onset age and the onset age plus 0.5 (assuming a uniform distribution in both cases).

Once the ages at which offspring were observed had been simulated, we subtracted 0.5 from their age to ease the simulation (see method section *Estimation of variation in parental investment* and discussion). We then simulated the parents’ behavior, *b*, in link-space (i.e., we first simulated log(*b*/(1-*b*)), i.e., logit(*b*)). This approach allows the simulation of a behavior that follows a trajectory in the form depicted in Fig. 2 (thick dashed line). In addition, this approach allows to easily simulate parents with varying propensities to display the behavior. To simulate varying parental investment, we randomly sampled one value for each parent from a normal distribution with a mean of zero and a standard deviation of 0.75 (Fig. 2). The sampled values per parent were then added to the fixed effects intercept, leading to variation of parents in their propensity to display the offspring directed behavior. Hence, in link space, the behavior was simulated as

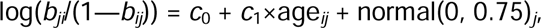

where *j* is the index of the *j*^th^ parent-offspring dyad, *i* is the *i*^th^ observation of the *j*^th^ parent-offspring dyad, *b_ij_* is the proportion of time the parent of the *j*^th^ parent-offspring dyad shows the behavior in the *i*^th^ observation (in link space), *c*_0_ is the intercept, *c*_1_ is the slope of the behavior against offspring age, and normal(0, 0.75)*_j_* represents the value randomly sampled from a normal distribution with a mean of 0 and a standard deviation of 0.75 which was assigned to the *j*^th^ parent-offspring dyad. In the next step, we transformed logit(*b*) to a space that represents the actual proportions, namely

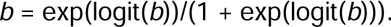

After that, *b* was bound between 0 and 1, and the variation in the trajectories of different parent-offspring dyads could, for instance, look as depicted in Fig. 2 (thin dashed lines).

Once the parent-offspring age-specific proportions had been determined, we sampled values from a beta distribution (Bolker 2008). In addition to its mean, the beta distribution has a precision parameter, *phi*, governing the magnitude of variation of the observations around their mean. We chose values of 5, 10, 20, and 40 for *phi,* which leads to expected distributions of the proportions as depicted in Fig. 4. Note that the larger the value of *phi*, the smaller the magnitude of stochastic variation of the actual observations. In practice, the value of *phi*, as estimated for empirical data, will likely depend on the duration of the focal periods to a considerable extent: assuming the behavior under study is a state which usually lasts for a while (such as nursing or carrying), then the magnitude of stochastic variation in the observed proportions will likely be smaller when observations periods are longer.

**Fig. 4.**
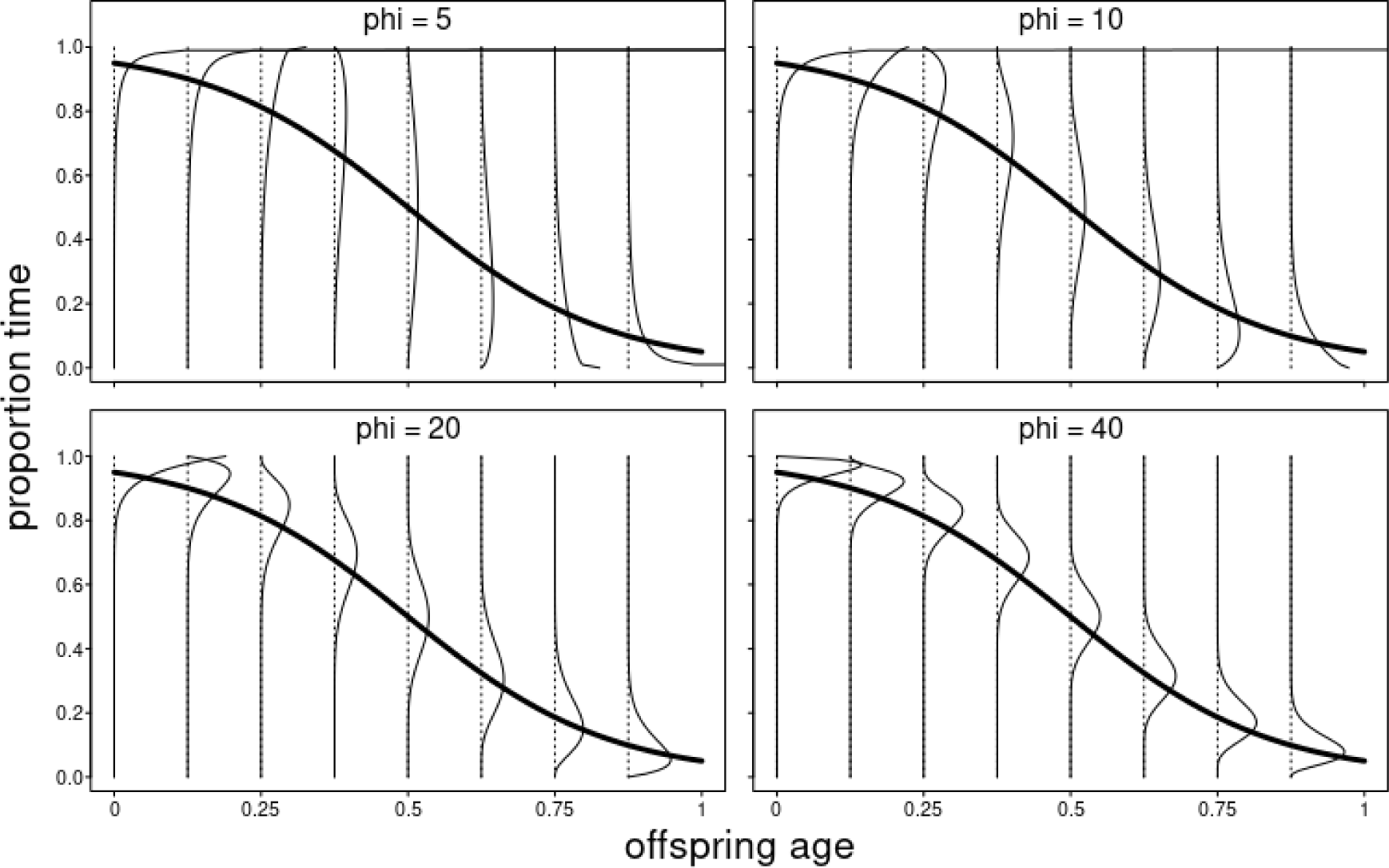
Illustration of the effect of the precision parameter *phi* on the shape of a beta distribution. Depicted are the average proportions (y-axes) and how they depend on offspring age (x-axes) in our simulations (thick solid line). Thin solid lines depict the density of beta distributions for a given age (vertical dotted lines). Different plots show different values of *phi*. Note that the larger the value of *phi*, the smaller the expected scatter of the actual observations around the simulated mean.

### Estimation of variation in parental investment

We analyzed each of the simulated data sets using two different methods, the conventional and the proposed modeling method. First, we determined for each parent the average proportion of time it showed the behavior across all observations (hereafter conventional parent-specific investment). However, these averages per parent express the parent-specific investment in response space, whereas the respective outcome of the modeling approach (see below) expresses them in link space. To directly compare the estimates of parent-specific investments and their variability between the conventional and the modeling approach, we transformed average investments per parent to link space. That is, we converted each average (‘b’) according to b’=log(b/(1-b)). The values of b’ then served as estimates of parent-specific investment in subsequent evaluations. For simulations under the null scenario, we also determined the standard deviation of b’ across parents separately for each simulated data set.

Second, we modeled parent behavior as a function of offspring age. To this end, we used a Generalized Linear Mixed Model (GLMM; Baayen 2008), fitted with a beta error distribution and logit link function (McCullagh & Nelder 1989), separately for each simulated data set. We modeled offspring age as a fixed effect and included the parent-offspring dyad as a random intercepts effect and a random slope of offspring age (Schielzeth & Forstmeier 2009; Barr et al. 2013). We did not estimate a parameter for the correlation among random intercepts and slopes, mainly because we simulated no random slope of age. Moreover, this decision helped to reduce computation time and increase the probability of model convergence. Before fitting the model, we subtracted 0.5 from age. This step has the effect that the parent-specific estimates of their investment refer to an offspring of an age corresponding to half of the considered period of development (which has relevance since the model comprised a random slope of age; see discussion). From the fitted model, we extracted Best Linear Unbiased Predictors (BLUPs; Baayen 2008) for the random intercepts of parent-offspring dyads as estimates of the parent-specific propensities to display the behavior (hereafter ‘modeled’ parental investment). We also extracted the standard deviation estimated for the random intercept effect of parent-offspring dyad. The standard deviation provides information about the estimated magnitude of variation in parental effort.

### Evaluation

We first evaluated the null scenario simulations in which parents, by definition, did not vary in their parental investment. To this end, we compared the estimated standard deviations for the paternal investment obtained from the two approaches (all sampling schemes). Since we simulated no variation in parental investment in this scenario, the estimated variation should ideally be zero. Secondly, we determined the extent to which the estimated parent-specific investments were correlated with the average age of the offspring (only random and constrained sampling). To this end, we regressed the estimated parent-specific investments against offspring age and determined the *R^2^* of the regression. We pooled across all simulations with the same parameter setting for number of parent-offspring dyads, number of observations per parent-offspring dyad, sampling scheme, and precision parameter *phi*. We ran this analysis only for the random and constrained sampling schemes because the average age of the offspring did not vary between dyads in the balanced sampling scheme (see Fig. 3).

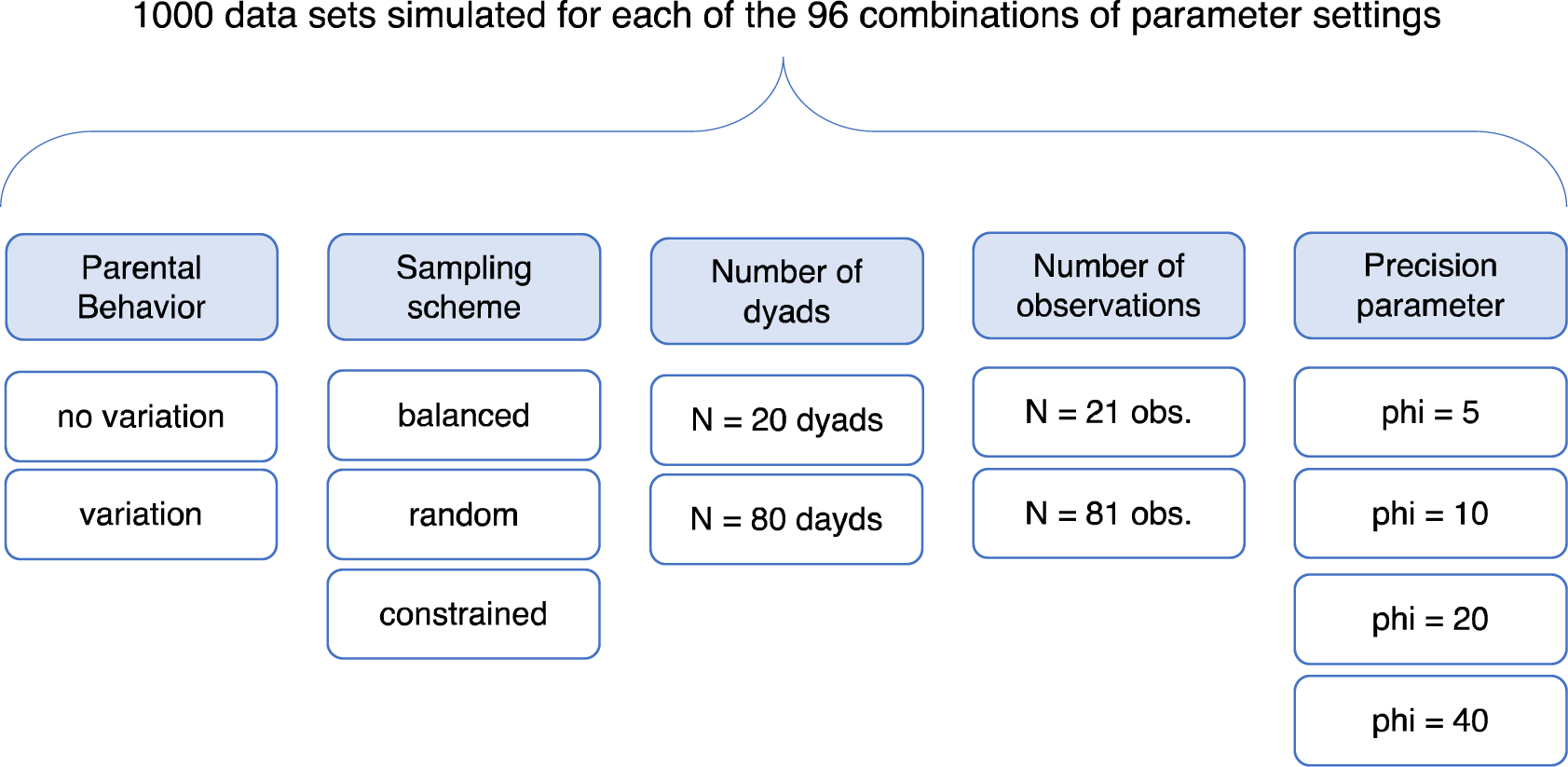

For the simulations under the variable parental investment scenario, we compared the estimates of parent-specific investment with the simulated ones. We also determined how much the estimates were explained by the average age of the offspring (only random and constrained sampling). In the case of the balanced sampling scheme, we regressed the estimates of the parent-specific investment against the simulated ones and determined the *R^2^*. In the case of random and constrained sampling, we regressed the estimates of the parent-specific investment against the simulated ones and the average age of the offspring. We determined partial *R^2^* for each of the two predictors. Again, we pooled the estimates of all simulated data sets with the same respective parameter setting.

### Implementation

The simulation and analysis of the simulated data were implemented in R (version 4.3.1; R Core Team 2023). For random sampling from uniform and beta distributions, we used the functions runif and rbeta, respectively. We fitted the models using the function glmmTMB of the equally named package (version 1.1.8; Brooks et al. 2017). We simulated 1000 data sets for each of the 96 combinations of parameter settings with regard to simulation scenarios (null scenario or variable parental investment scenario), sampling scheme (balanced, random, or constrained), number of parent-offspring dyads (20 or 80), number of

## Results

### No variation between parents

When we simulated no variation between parents, the conventional method strikingly overestimated the variation among parent-specific behaviors (Fig. 6; for detailed results, see Supporting Information Fig. SI 1 to SI 4). This overestimation was lowest for balanced sampling and highest for constrained sampling. Compared to the effects of the sampling scheme, all other simulated parameters had relatively minor effects: the magnitude of overestimation was slightly larger in case of 21 as compared to 81 observations per dyad, particularly in case of random sampling. Furthermore, the standard deviation slightly decreased with decreasing magnitude of stochastic variation in the observed behavior (precision parameter *phi*). The number parent-offspring dyads had a relatively minor effect.

**Figure 6.**
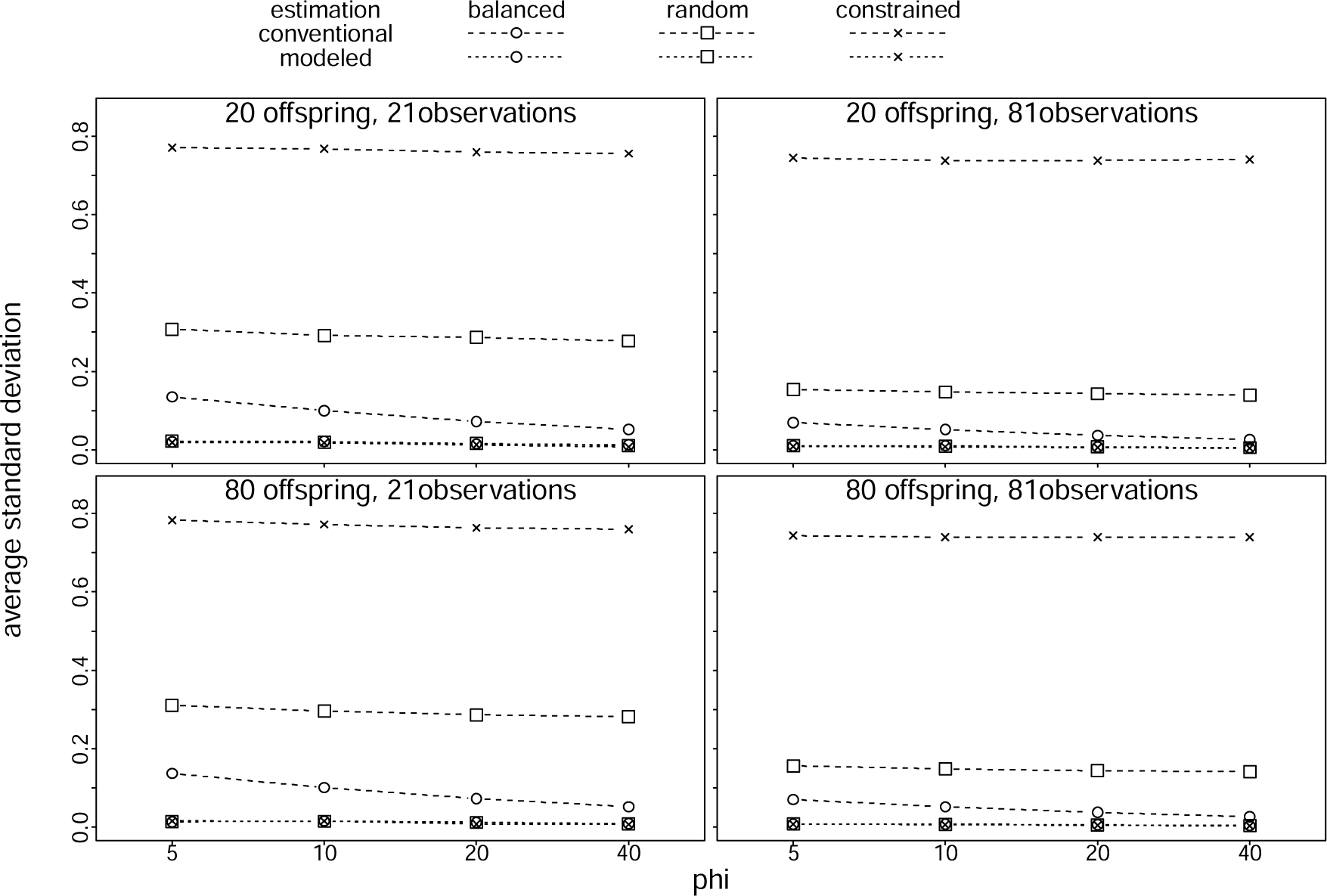
Overview of the results of the simulation when we simulated no differences between parents with regard to their behavior. Depicted are the average conventional (dashed lines) and modeled (dotted lines) estimated standard deviations of parental behavior (y-axes), separately for balanced sampling (open dots), random sampling (crosses), and constrained sampling (laying crosses), against the simulated precision parameters (x-axes). Each plot depicts the results for a given number of parent-infant dyads and number of observations per dyad. Detailed results are depicted in Fig. SI 1 to SI 4.

In the case of random and constrained sampling, the overestimation of variation in parent-specific behavior was mainly driven by the varying ages at which offspring were observed. We found that the parent-specific estimates of their behavior strongly correlated negatively with offspring age (Fig. 7), whereby the correlation increased with increasing precision parameter phi and was stronger for constrained than for random sampling (Fig. 7, Fig. SI 5 to SI 12). The effects of sample size on the magnitude of the correlation were relatively minor.

**Fig. 7.**
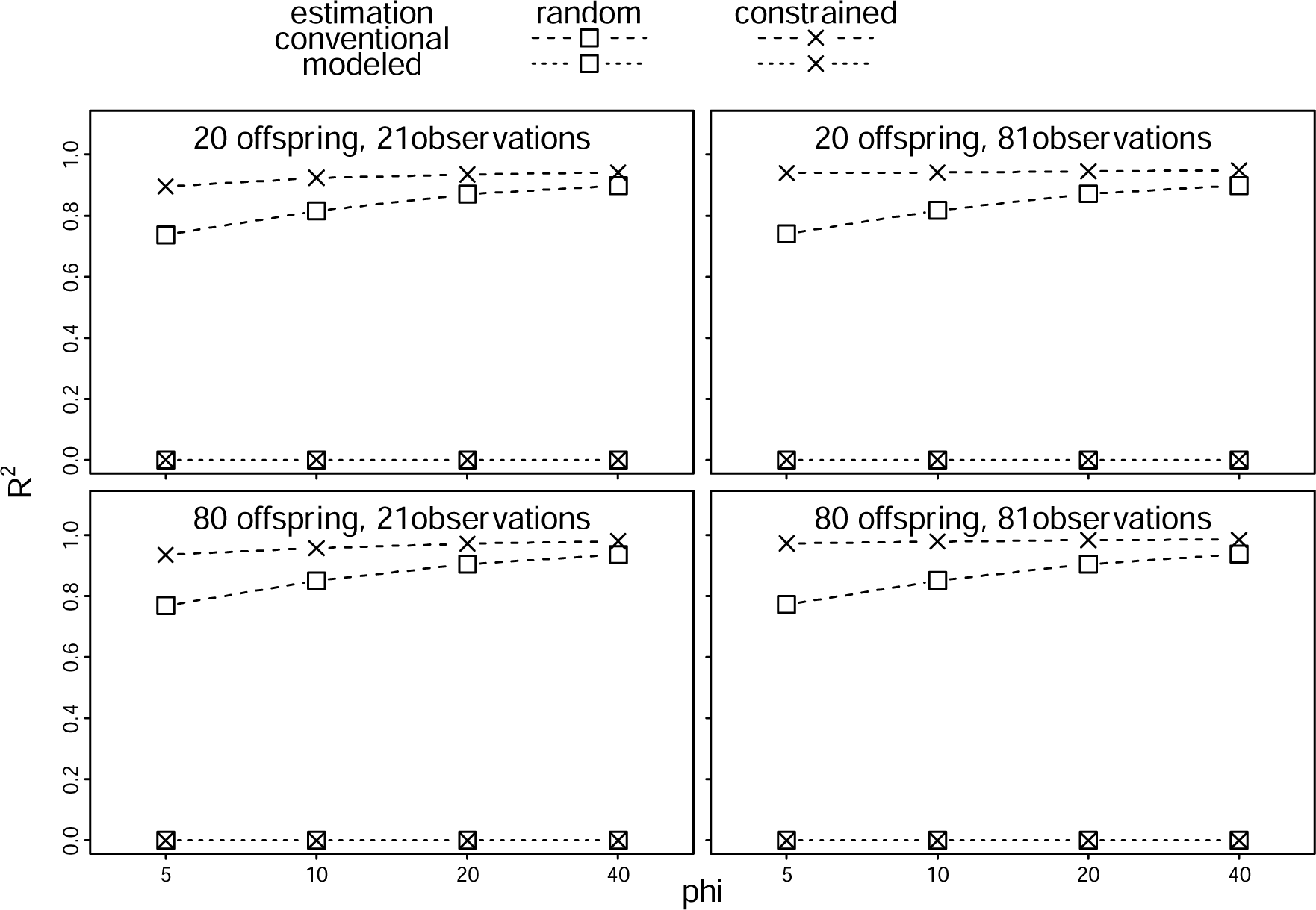
Explained variance (R^2^) of regressions of estimated parental behavior against average offspring age for different sample sizes, sampling schemes, precision parameters, and estimation methods. The simulated data did not comprise any variation among parents with regard to their parental behavior. Detailed results are shown in Fig. SI 5 to SI 12.

The modeling approach, in contrast, consistently estimated smaller standard deviations of parental behavior which, on average, were essentially zero (Fig. 6, Fig. SI 1 to SI 4). Nevertheless, also when modeling the magnitude of variation in parental behavior, we found that the among-parent variation (which was simulated to be zero) was occasionally overestimated. The sampling scheme had a relatively minor effect on the magnitude of overestimation. Instead, the estimated standard deviations slightly decreased with an increasing number of observations per parent-offspring dyad and with increasing precision parameter *phi* (Fig. SI 1 to SI 4). When we modeled variation in parent-specific behavior, it did not correlate with offspring age (Fig. 7, Fig. SI 5 to SI 12).

### Variation between parents

Across all simulation parameter settings (i.e., each combination of sampling scheme, number of parent-offspring dyads, number observations per dyad, and precision parameter *phi*) and estimation method (conventional or modeling), the estimated parent-specific behaviors correlated with the simulated ones (Fig. 8; for detailed results see Fig. SI 13 to Fig. SI 24). In all combinations of parameter settings, we found a large variance explained by regressions of estimated against simulated parental behavior (i.e., > 0.75.). Nevertheless, the variance explained increased with decreasing magnitude of stochastic variation in the simulated behavior (increasing precision parameter phi), was higher with a larger number of parent-offspring dyads and also with a higher number of observations per dyad. Furthermore, although in almost all parameter settings the differences between different sampling schemes and the two estimation methods were relatively low, the variance explained by the regression was clearly lower when applying the conventional method in the case of constrained sampling combined with having only 20 parent-offspring dyads. Thus, in this specific parameter setting the conventional method greatly underperformed compared to the modeling method.

**Fig 8.**
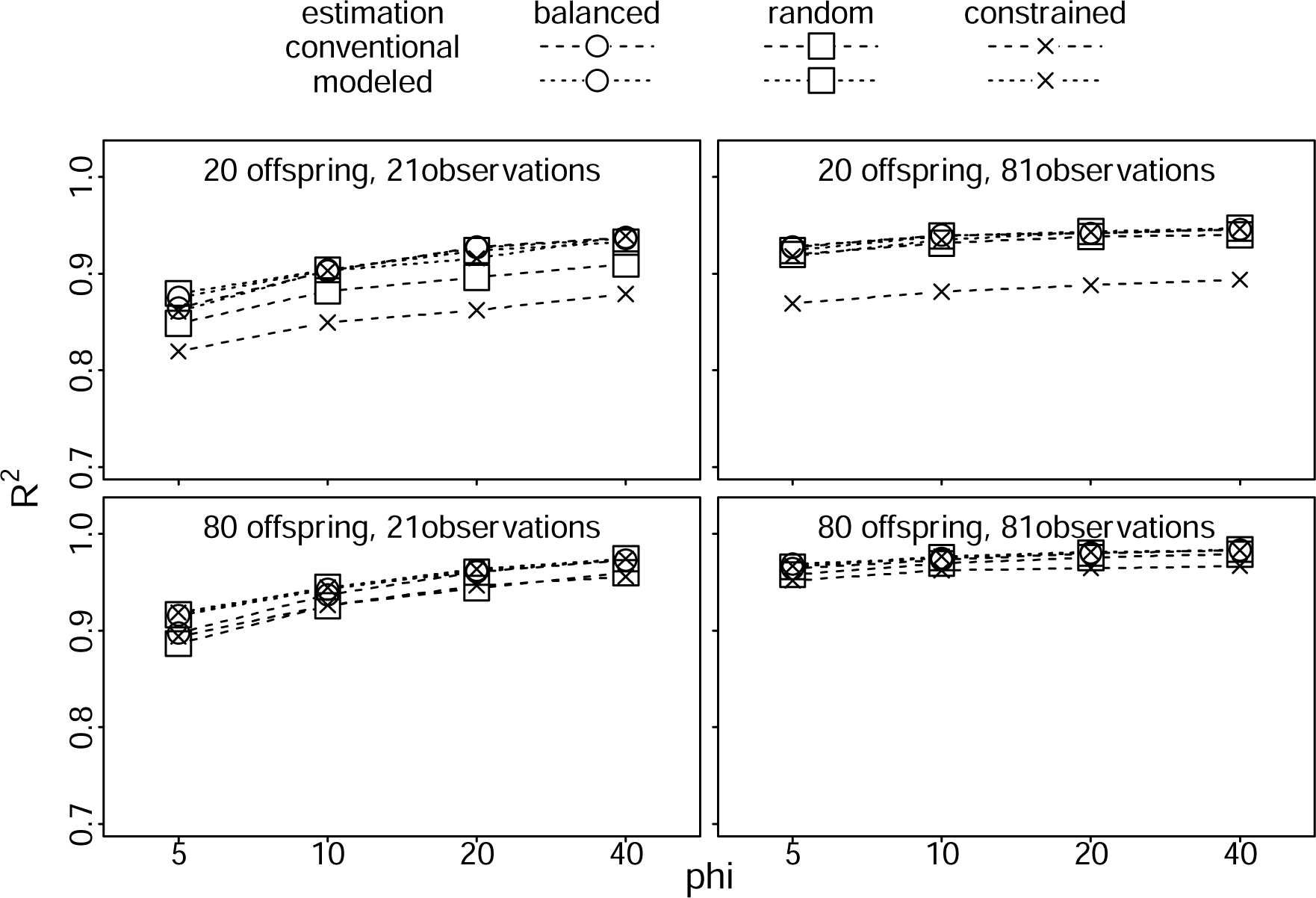
Explained variance (*R*^2^ or partial *R*^2^) of regressions of estimated parental behavior against simulated parental behavior for different sample sizes, sampling schemes, precision parameters, and estimation methods (for more detailed results see Fig. SI 13 to Fig. SI 24). The simulated parents varied in their parental behavior (i.e., varying-parent scenario).

Despite the clear correlation between the estimated and the simulated parent-specific behavior, we also found that the conventionally estimated parent-specific behavior strongly correlated with the average age of the offspring in case of random or constrained sampling (Fig. 9, 10; for detailed results, see Fig. SI 17 to Fig. SI 24). Partial *R*^2^ of conventionally estimated parent-specific behaviors regressed against the average age of the offspring were invariably at least ca. 0.5 in the case of random sampling and at least ca. 0.8 in the case of constrained sampling (Fig. 10). Practically, this means that the parent-specific behavior was overestimated for younger and underestimated for older offspring (Fig. 10, SI 17, SI 19, SI 21, and SI 23). In contrast, when modeling parent-specific behavior, we did not find such a correlation between the average offspring age and the estimate of parent-specific behavior. The respective partial *R*^2^ were invariably about zero (Fig. 9, 10, SI 18, SI 20, SI 22, and SI 24), and a bias of the modeled parent-specific estimates by offspring age was usually not recognizable. The only exception was the simulation with considerable stochastic variation in the simulated behavior (precision parameter phi = 5) and constrained sampling. Here, the parent-specific behavior was slightly overestimated for older and slightly underestimated for younger offspring (Fig. SI 22). However, this bias was much smaller than the one we found when using the conventional method (Fig. SI 21).

**Fig. 9.**
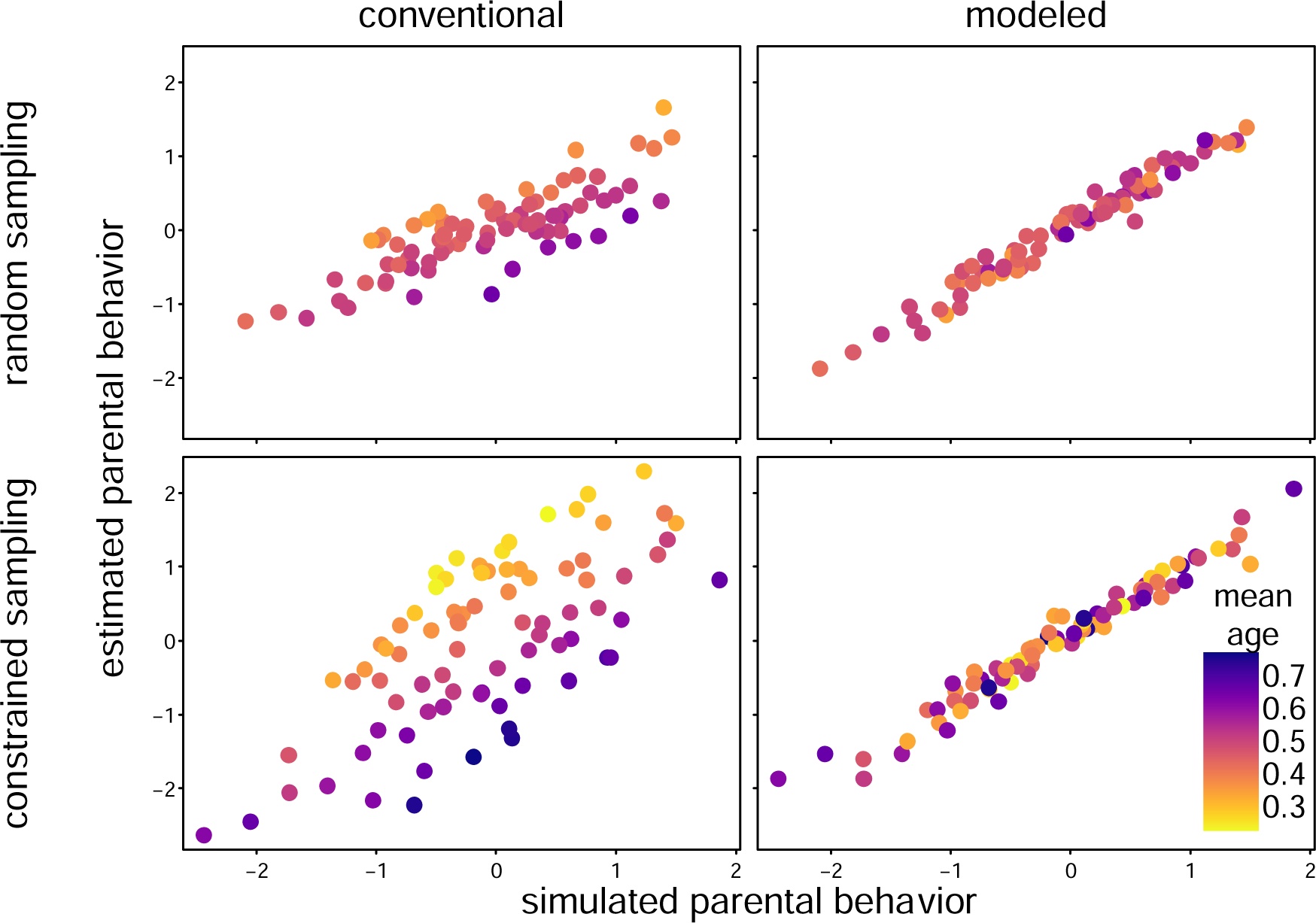
Estimated parental behavior (y-axes) and how it varied with simulated parental behavior (x-axes) and average offspring age (colors). Depicted are results for two randomly chosen simulated data sets (one with random (top) and one with constrained (bottom) sampling) with 80 parent-offspring dyads, 21 observations per parent-offspring dyad, and a precision parameter of 10. Note that the conventionally estimated parent-specific estimates clearly correlated with average offspring age whereas the modeled estimates did not. For detailed results see Fig. SI 17 to SI 24.

**Fig 10.**
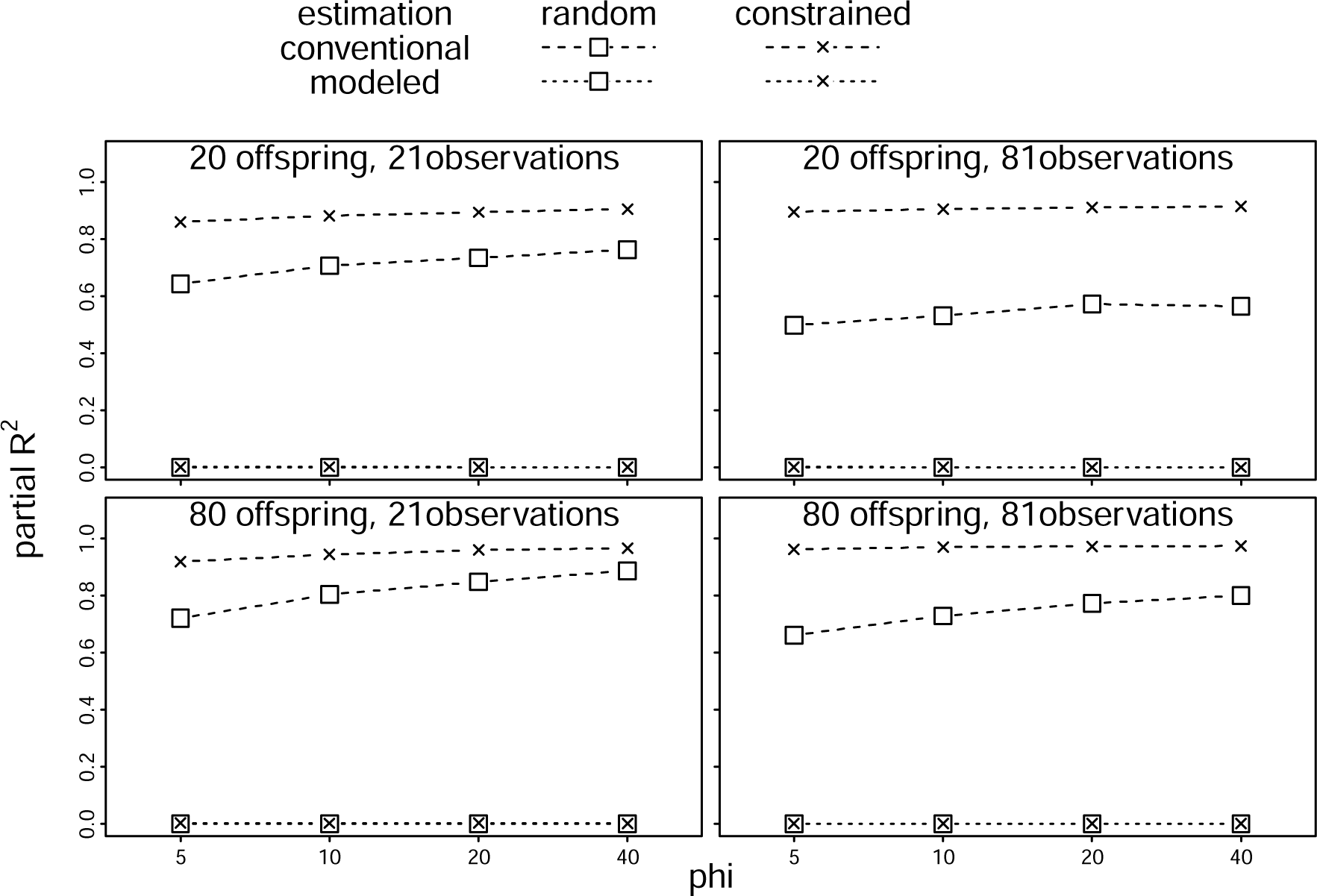
Explained variance (partial R^2^) of regressions of estimated parental behavior against offspring age for different sample sizes, sampling schemes, precision parameters, and estimation methods (for more detailed results Figures see Fig. SI 17 to Fig. SI 24). In the simulation, parents varied with regard to their parental behavior.

## Discussion

Overall, the conventional approach was much poorer than the modeling approach in estimating variation among parental behavior. When we simulated no variation among parents, the conventional method consistently overestimated the magnitude of variation among parents. In contrast, the modeling approach did so only occasionally, and when it did, to a lesser extent than the conventional approach. The overestimation of among-parent variation by the conventional approach was largely driven by the fact that the estimates of parent-specific behavior negatively correlated with offspring age in the case of random and constrained sampling. But it was remarkable that this overestimation even happened in the case of completely balanced sampling in which offspring did not vary at all concerning at which ages they were observed. Also, when we simulated variation among parents with regard to their behavior, the modeling approach was superior to the conventional approach. This result was again mainly driven by random and constrained sampling leading to a negative correlation between average offspring age and the conventionally estimated parent-specific behavior. Hence, the message from our simulations is unequivocal: just averaging parental behavior should not be used. We show that our modeling approach is far superior. Importantly, we here used a parent behavior being a proportion as an example, but we do not see any reason, why our results should be different with a behavior being, for instance, a count, an unbound quantity, or binary. In the following, we briefly discuss some of the issues that may arise when using the modeling approach we proposed here and flag some potentially important points to keep in mind when using it.

### Inclusion of random slopes of offspring age

The model we fitted to the simulated data included a random slope of offspring age within the parent-offspring dyad. We did so for two reasons: first, even though we did not simulate such a random slope to be present in the simulated parental behavior, it seems likely that any real data comprise such a component. There is no reason to assume that parents vary in their average parental behavior but not at all in the trajectories their behavior follows when the offspring gets older. Second, it has been repeatedly shown that the failure to include random slopes present in the observed response variable can lead to ‘overconfident’ models and drastically inflated type I error rates (Schielzeth & Forstmeier 2009; Barr et al. 2013; Aarts et al. 2015). Even though the model we proposed here does not capitalize on significance testing, it should be best practice to include such random slopes whenever they are theoretically identifiable.

Crucially, including a random slope of offspring age within parent-offspring dyad has an important consequence on the assessment of parent-specific behavior. Specifically, a random slope of a covariate like offspring age within the random effects factor parent-offspring dyad is conceptually equivalent to their interaction. Thus, the parent-specific trajectories of their behavior against offspring age might cross during offspring development (Fig. 11). For instance, a parent that shows a relatively high care behavior (compared to other parents with offspring of the same age) when its offspring is relatively young, might show relatively little care (compared to other parents with infants of the same age) when its offspring is older. It had thus been suggested to mean-center the covariates for which one includes random slopes (here, offspring age; Araya-Ajoy et al. 2015). As a result, the BLUPs are conditional on an offspring age which equals the average offspring age across the entire data set (vertical line in Fig. 11). Practically this means that the estimates of parent-specific behavior represent the differences between parents at a certain offspring age but not across the entire period of offspring development considered (Fig 11; see the supporting information for a potential ad-hoc *s*olution for extracting the estimated parent specific behavior in the presence of a random slope of offspring age).

**Fig. 11.**
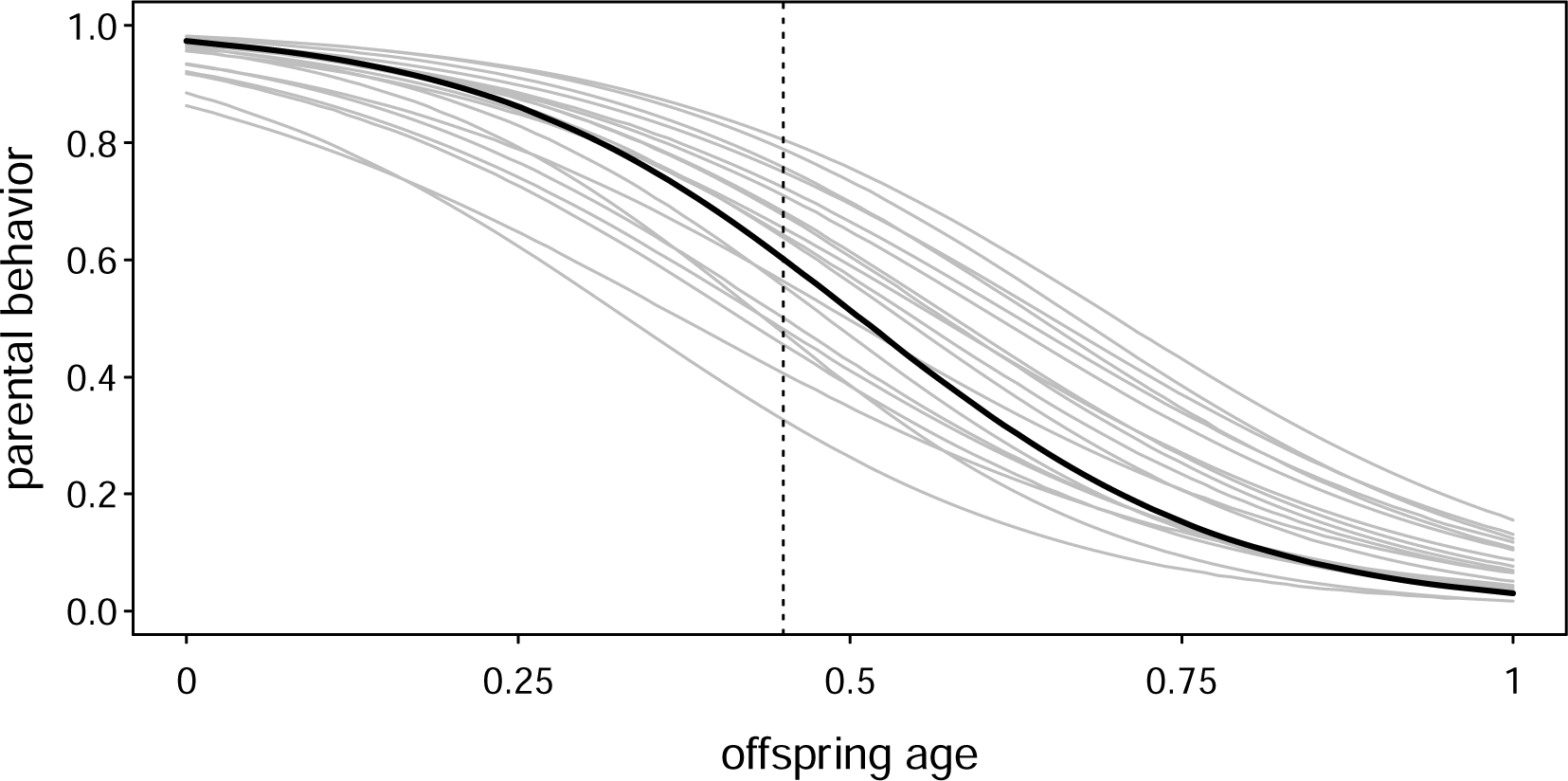
Effect of including a random slope of offspring age within parent ID on the assessment of parent-specific behavior. Including the random slope has the effect that the model can reveal parent-specific trajectories of the modeled behavior against offspring age (different lines). Importantly, under such conditions, a parent can potentially no longer be unambiguously characterized as showing a relatively high (or low) care behavior. For instance, the parent depicted by the thick line displays the behavior relatively frequently (compared to other parents with offspring of the same age) when its offspring is young and relatively rarely when its offspring is older. When mean centering offspring age, then the BLUPs of parent—specific behavior represents the relative care behavior for offspring of an age equaling the average age across the entire data set (e.g., at the vertical line), but this is not necessarily equivalent to their relative behavior over the entire period considered.

A side aspect of including a random slope of offspring age within parent is that one could also estimate a parameter for the correlation between the parent-specific random intercepts and slopes (e.g., Barr et al. 2013). The models we fitted to the simulated data did not include such a parameter as we did not simulate one present in the response and because we wanted to ease model convergence and reduce computation time. It is probably advisable to include such a correlation parameter in any actual study for two reasons. First, it seems plausible that such a correlation is present in the data. For instance, one could imagine that parents that invest relatively much into their offspring also decrease their care behavior faster as their offspring gets older compared to parents who invest relatively little into their offspring. Second, although the effects of neglected correlation parameters are less investigated than the effects of neglected random slopes and, so far, seem less severe (e.g., Schielzeth & Forstmeier 2009; Barr et al. 2013), there are still clear indications that neglecting such correlation parameters can lead to overconfident models and type I errors. Hence, we feel it should be a regular practice to include them.

### Estimating the effects of variables such as age or parity on parental behavior

Frequently, estimating variation in parental behavior is not the goal of the analysis but rather an intermediate step. The ultimate aim may instead be to examine the relationship between parental behavior and other variables such as parent age, parity, brood or litter size, social rank, weather, or food availability. Some of these predictor variables vary only between parent-offspring dyads (e.g., parity), while others may change on a daily basis (e.g., temperature). Integrating such questions into the modeling approach proposed here is straightforward, as it uses parental behavior as the response in a model that can well comprise other predictors in addition to offspring age. In case one might want to estimate the effects of parity and temperature on the proportion of time a parent stays in body contact with its offspring, the model would be something like (in lme4/glmmTMB syntax):

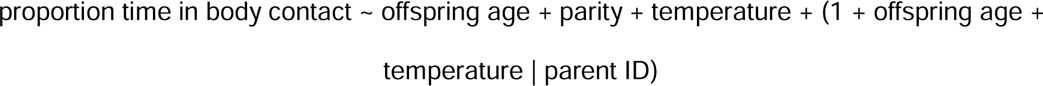

Note that the model would only have an appropriate random-effects structure when each parent has only one offspring in the data (see below). In such a model, one would not estimate the effect of, e.g., parity on an estimated measure of parental behavior but instead directly estimate the effect of parity on the behavior considered. If, for instance, the fixed effects estimate of parity is positive, this means that parents display the behavior the more the more offspring they already have. If such a model reveals a random intercept of parent ID or a random slope of offspring age not being essentially zero, this would give hints to parents varying in offspring care beyond what is explained by parity (what might be considered parental ‘style’) and/or the presence of effects other than parity that vary between parents, which affect how much they invest into their offspring.

### Estimating the effects of parental behavior on variables such as offspring growth or survival

In the previous section, we addressed how one could model the effects of other variables on parental behavior. Still, one might also be interested in how parental behavior affects the fate of the offspring (e.g., its growth, survival, or lifetime reproductive success). A simple but not very satisfactory solution would be first to model parental behavior as proposed here, extract the parent-offspring-specific estimates of their behavior (e.g., BLUPs for the random intercept of parent-offspring dyad), and then use them as predictors in subsequent analyses, such as offspring survival models. Such an approach is not very satisfactory because it takes the point estimates for the BLUPs as a given and neglects the uncertainty associated with them (Houslay & Wilson 2017). A preferable solution would be to propagate their uncertainty into the follow-up analysis. Ideally one would integrate both steps into a single model; that is, to model parent behavior as proposed here and at the same time model the effect of parent-specific behavior on the offspring’s fate (e.g., survival). Unfortunately, to our knowledge, a simple ‘canned’ solution for such an analysis is not available in packages written for the statistical language R. Yet, stan (Stan Development Team 2023), a language for Bayesian analysis to which R interfaces, should make such an analysis possible.

### Multiple variables of parental behavior

Frequently, researchers investigating parental behavior have data for several variables, which they consider potentially informing about ‘parental style’. The question is how to proceed with the analysis in such a case. First, researchers should ensure that the variables entered in the analysis occur sufficiently frequently. Second, not all variables necessarily reflect the same latent variable. For instance, the proportion of time parent and offspring are in body contact and the number of times the parent rejects an offspring’s attempts to initiate body contact might not be driven by the same underlying process and may thus follow different trajectories. It is possible to model the trajectories of several variables of parental behavior simultaneously in the framework of an R package for Bayesian implementations for mixed models, and in the stan environment, this should even be possible for variables of parental behavior which would need different error distributions (e.g., proportions and counts).

### Non-linear trajectories

When studying the magnitude of parental behavior as a function of offspring age, one must be aware that a simple linear model might not be an appropriate solution. This is because the linear model is linear in link space, imposing clear limitations about which functional relationships it can deal with and reveal. For instance, a standard linear model with a count response that is fitted with a log link function (usually the default of Poisson and negative binomial models) does not have an upper asymptote and always has a lower asymptote of zero. A lower asymptote of zero, however, might not be able to capture how the frequency of a behavior develops as an offspring ages. For instance, many primates are female philopatric, which means that female offspring stay in their natal group and hence have the possibility to interact with their mothers for a potentially large part of their adulthood. Now assume a behavior like a friendly approach, which mothers might commonly show when their offspring is young and which becomes rarer with increasing offspring age but never goes down towards zero. A standard linear count model with a log link function will not correctly capture such a trajectory (Fig. 12). Equivalently, standard logit link linear models, which are typically used for response variables that are proportions or binary, always have a lower asymptote of 0 and an upper asymptote of 1, which might not allow capturing the trajectory of the behavior appropriately as the offspring age increases. In such cases, one will need a non-linear model.

**Fig 12.**
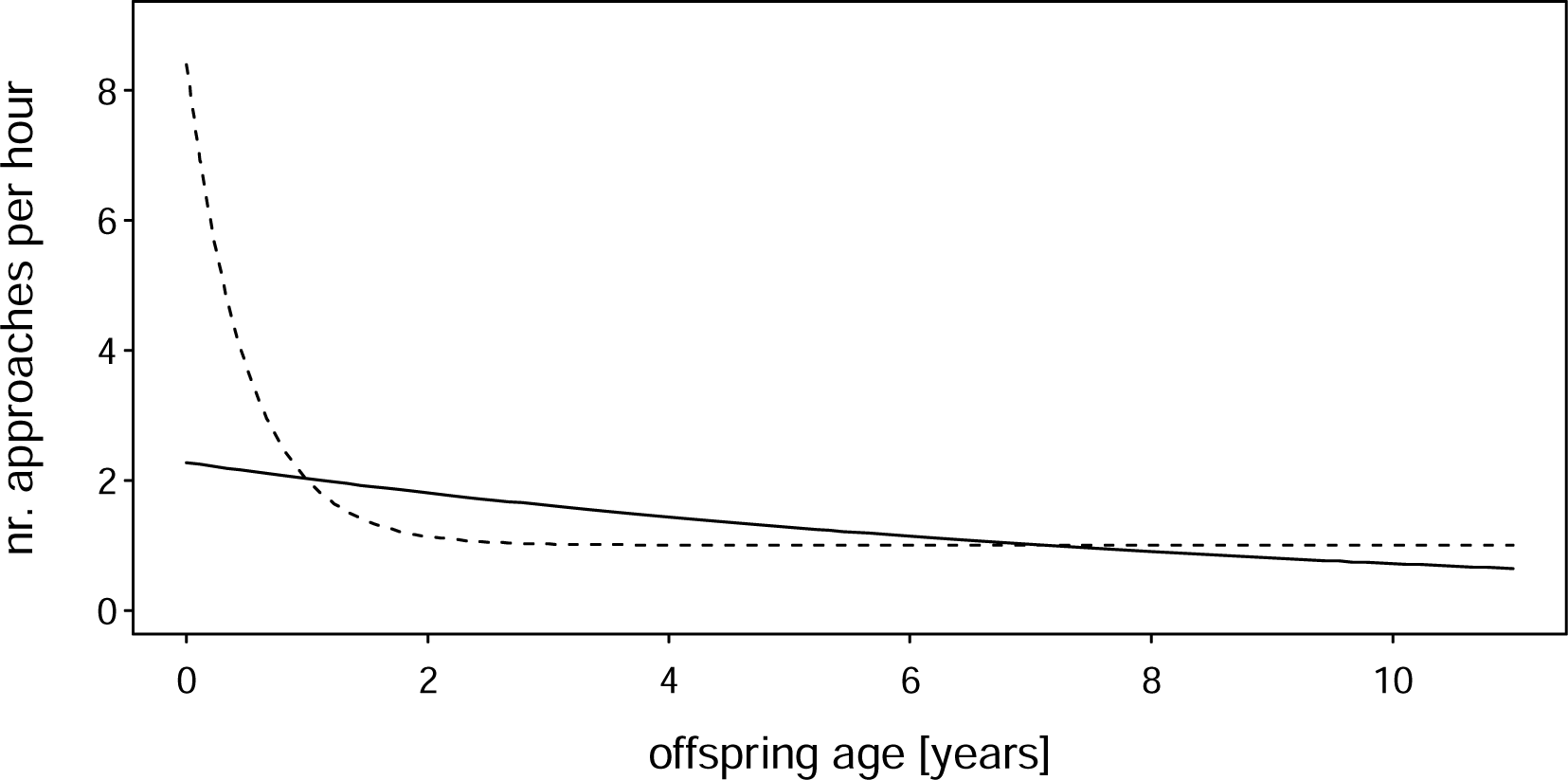
Actual and estimated effect of offspring age when using a standard linear model in a situation in which it is not appropriate. Dots depict the (simulated) parent behavior and the dashed line its offspring age dependent trajectory. When the offspring is young, the behavior occurs at relatively high rates and as the offspring reaches adulthood the rate fairly rapidly approaches an asymptote of one occurrence per hour. A standard count model fitted to these data reveals the relationship depicted by the solid line, which is not capturing the actual effect.

Finally, it is possible that certain parental behaviors are particularly common when the offspring has an intermediate age. For instance, a parent might provide more food to an offspring when it has an age about half-way between birth or hatching and independence. Such trajectories in which the behavior peaks at an intermediate offspring age may be modeled by including offspring age squared in addition to offspring age. However, care must be taken that such a model appropriately captures the actual offspring age-dependent trajectory of behavior. If it does not, one will again need a non-linear model. Setting up and fitting non-linear mixed models is a world of its own and beyond the scope of this paper. Nevertheless, we expand on the issue a little more in the Supporting Information. Nevertheless, we believe that in many cases, a linear model provides a sufficiently accurate approximation.

#### Parents with more than one offspring present in the data

A further aspect worth being considered is how to proceed with data comprising several offspring per parent. Such data could, for instance, arise from long-term studies in which parental behavior is observed for several consecutive offspring per parent. The model to be fitted then has to include a random effects factor for the identity of the parent and also one for the identity of the offspring (both associated with appropriate random slopes). The random effect of parent ID informs about the potential within-parent consistency that persists across multiple offspring, while the random effect of offspring ID accounts for offspring-to-offspring variation in parental behavior of the same parent towards different offspring. Importantly, from the perspective of the offspring, the parental ‘style’ it receives is best characterized by the addition of the BLUPs obtained for its parent and the offspring itself.

#### Concluding remarks

The conventional way of characterizing parental behavior is usually seriously biased as it does not consider the fact that parental behavior towards an offspring usually correlates with offspring age. As an alternative, we have presented a modeling approach to estimate parental behavior towards an offspring such that is not confounded by offspring age. Our approach is simple to apply in many situations, but it also has certain limitations one needs to be aware of when using it. Irrespective of these limitations, our modeling approach reveals estimates of parent specific behaviors that are not affected by average offspring age and greatly outperforms the conventional method when estimating parental care behavior.

## Supporting information

commented_code_for_parental_care

