## Supplementary material for "Overcoming Pitfalls when Estimating Variation in Parental Care: A Simulation Study": commented_code_for_parental_care: commented_code_for_parental_care.r

```
##code for the simulations conducted by [authors] for
##"Overcoming Pitfalls when Estimating Variation in Parental Investment: A Simulation Study"

##the code comprises four main sections:
  ##the first section comprises the simulation itself;
  ##the second section summarizes the simulation results into several lists (this mainly aims at making the rendering of the Supporting information faster);
  ##the third section shows how to create the Figures with results shown in the main text;
  ##the last section shows how to create the Figures with results shown in the Supporting information.

##note that throughout we simulated data sets with one offspring per parent and one parent per offspring
  ##that's why we use number offspring, number infants, and number parents interchangeably
##note also that originally we used the term 'empirical' for what we now label 'conventional' estimates
  ##that's why this term (also abbreviated 'emp') shows up in the code

##SECTION 1: code for simulation
##in this script we run a total of 6 simulations (1A, 1B, 1C, 2A, 2B, 2C)
##these are ( (see Fig. 3 of main text for an illustration of balanced. random, and constrained sampling):
  ##simulation 1, no variation between parents wrt mothering style:
    ##A balanced sampling
    ##B random samplingby by by 
    ##C constrained sampling
  ##variation between mothers wrt mothering style:
    ##A balanced sampling
    ##B random sampling
    ##C constrained sampling
##after running the simulations we summarize the results
##and after that we show how to create the plots shown in the main text and supporting information

##load librarys needed:
library(glmmTMB)
library(viridisLite)
library(kyotil)
##define function for transforming between the two parameterizations of the beta distribution (mu-phi and shape1-shape2):
beta.par1topar2<-function(shape1=NULL, shape2=NULL, mu=NULL, phi=NULL){
	if(is.null(shape1) & is.null(shape2) & !is.null(mu) & !is.null(phi)){
    shape1=mu*phi
		shape2=(1-mu)*phi
		return(list(shape1=shape1, shape2=shape2))
	}else if(!is.null(shape1) & !is.null(shape2) & is.null(mu) & is.null(phi)){
		mu=shape1/(shape1+shape2)
		phi=shape1+shape2
		return(list(mu=mu, phi=phi))
	}else{
		stop("don't know what to do: either you give me shape1 and shape2 or mu an phi")
	}
}
##define function transforming the response such that values being exactly 0 or 1 are ruled out (see Smithson, M. and Verkuilen, J. (2006). Psychological Methods, 11: 54–71.):
beta.tr<-function(x){
	if(any(is.na(x))){warning("x comprises NAs")}
	return((x*(length(x) - 1) + 0.5)/length(x))
}

##definition of constant parameters to be simulated:
  age.range=c(0, 2)##range of offspring age
  ##parameterize for simulation with mean age (i.e., 1) set to 0:
  coefs=c("(Intercept)"=qlogis(0.5), age=(qlogis(0.05)-qlogis(0.95))/2)
  names(coefs)[2]="sim.age"
  mo.icpt=0##a parameter not evaluated
  n.sims=1000
  parent.sd=0.75##standard deviation of normal distribution for which-the parent specific deviations from average investment are sampled (see Fig. 2 of main text)
  save.path=""##path where to store workspaces with simulation results
  
##define function running the simulations for a given number of parents and observations per parent and a given value of the presition parameter phi (see Fig. 4 of the main text):
sim.fun<-function(phi, n.parents, n.samples, n.sims){
  print(paste(c("NP", n.parents, "_NS", n.samples, "_PHI", phi), collapse=""))
  ##create environment from which to export objects needed to run the simulation in parallel mode:
  EP=rlang::env(age.range=age.range, coefs=coefs, mo.icpt=mo.icpt, parent.sd=parent.sd, beta.tr=beta.tr, beta.par1topar2=beta.par1topar2, n.parents=n.parents, n.samples=n.samples, phi=phi)
  ##and export them:
  clusterExport(cl=cl, varlist=c("age.range", "coefs", "mo.icpt", "parent.sd", "beta.tr", "beta.par1topar2", "n.parents", "n.samples", "phi"), envir=EP)
  ##simulation 1:
    ##1A (no variation between parents, balanced sampling):
      print("1A")
      ##create basic data set (is the same for all simulated data sets for simulation 1A):
      base.data=as.data.frame(expand.grid(parent.ID=as.factor(paste("m", 1:n.parents, sep=".")), age=seq(age.range[1], age.range[2], length.out=n.samples)))
      ##rescale age such that the value of zero corresponds to an age of one year:
      base.data$sim.age=base.data$age-1
      ##store number observations in data set:
      n=nrow(base.data)
      ##create model matrix (makes the simulation of the response easier):
      m.mat=model.matrix(object=~sim.age, data=base.data)
      ##create the mean response in ink space:
      xx=plogis(as.vector(m.mat[, names(coefs)]%*%coefs))
      ##... and determine the two shape parameters of the beta distribution to be simulated:
      beta.par=beta.par1topar2(mu=xx, phi=phi)
      ##(note that in simulation 1A there is no variation between parents, so the mean response is a given for a given offsrping age;
        ##that's why in this simulation the parameters of the beta disrtribution need to be determined only once)
      ##create environment from which to export more objects needed to run the simulation in parallel mode:
      E1=rlang::env(base.data=base.data, n=n, m.mat=m.mat, beta.par=beta.par, beta.par1topar2=beta.par1topar2)
      ##and export them:
      clusterExport(cl=cl, varlist=c("base.data", "n", "m.mat", "beta.par"), envir=E1)
      ##run the simulations for 1A:
      all.res.1A=parLapply(cl=cl, X=1:n.sims, function(s){
        ###create resonse:
        base.data$rv=rbeta(n=n, shape1=beta.par$shape1, shape2=beta.par$shape2)
        ##transform it such values being exactly 0 or 1 are ruled out:
        base.data$rv=beta.tr(base.data$rv)
        ##fit model:
        m=keepWarnings(glmmTMB(rv~age+(1+age||parent.ID), data=base.data, family=beta_family))
        ##extract warnings and result:
        w=m$warnings
        m=m$value
        ##extract estimated (i.e., modelled) standard deviation of variation in parental investment:
        re.sd=attr(summary(m)$varcor$cond$parent.ID, "stddev")
        ##extract estimated BLUPs:
        BLUPs=ranef(m)$cond$parent.ID
        ##determine conventional estimates of variation in parental investment (just the the average per parent):
        emp=tapply(base.data$rv, base.data$parent.ID, mean)
        ##and return the results:
        return(list(re.sd=re.sd, BLUPs=BLUPs, emp=emp, w=w))
      })
      ##extract conventional and modeled standard deviation of variation in parental investment from the list returned:
      emp.sd.1A=unlist(lapply(lapply(all.res.1A, "[[", "emp"), sd))
      mod.sd.1A=unlist(lapply(lapply(all.res.1A, "[[", "re.sd"), "[[", 1))

    ##1B (no variation between parents, balanced sampling):
      print("1B")
      ##run the simulation:
      all.res.1B=parLapply(cl=cl, X=1:n.sims, function(s){
        ##create data set wrt fixed and random effects (this time each individual data set needs be simulated from scratch as the ages at which parents are observed
          ##vary stochastically betewwn them):
        base.data=data.frame(parent.ID=as.factor(rep(paste("m", 1:n.parents, sep="."), each=n.samples)), age=round(runif(n=n.samples*n.parents, min=age.range[1], max=age.range[2]), 3))
        ##rescale age such that the value of zero corresponds to an age of one year:
        base.data$sim.age=base.data$age-1
        ##create model matrix (makes the simulation of the response easier):
        m.mat=model.matrix(object=~sim.age, data=base.data)
        ##create the mean response in ink space:
        xx=plogis(as.vector(m.mat[, names(coefs)]%*%coefs))
        ##determine shape 1-shape2 paranmeterization of the response to be simulated:
        beta.par=beta.par1topar2(mu=xx, phi=phi)
        ##... and simulate the response:
        base.data$rv=rbeta(n=n, shape1=beta.par$shape1, shape2=beta.par$shape2)
        ##transform it such values being exactly 0 or 1 are ruled out:
        base.data$rv=beta.tr(base.data$rv)
        ##fit model:
        m=keepWarnings(glmmTMB(rv~age+(1+age||parent.ID), data=base.data, family=beta_family))
        ##extract warnings and result:
        w=m$warnings
        m=m$value
        ##extract estimated (i.e., modelled) standard deviation of variation in parental investment:
        re.sd=attr(summary(m)$varcor$cond$parent.ID, "stddev")
        ##extract estimated BLUPs:
        BLUPs=ranef(m)$cond$parent.ID
        ##determine conventional estimates of variation in parental investment (just the the average per parent):
        emp=tapply(base.data$rv, base.data$parent.ID, mean)
        ##and return the results (this time we include also the average age per offspring)::
        return(list(re.sd=re.sd, BLUPs=BLUPs, emp=emp, inf.mean.age=tapply(base.data$age, base.data$parent.ID, mean), w=w))
      })
      ##extract conventional and modeled standard deviation of variation in parental investment from the list returned:
      emp.sd.1B=unlist(lapply(lapply(all.res.1B, "[[", "emp"), sd))
      mod.sd.1B=unlist(lapply(lapply(all.res.1B, "[[", "re.sd"), "[[", 1))
      ##extract conventional and modeled BLUPS of parent specific investment from the list returned:
      mod.where.1B=unlist(lapply(lapply(all.res.1B, "[[", "BLUPs"), "[[", "(Intercept)"))
      emp.where.1B=unlist(lapply(all.res.1B, "[[", "emp"))
      ##extract mean age per offsrping from the list returned:
      mean.age.1B=unlist(lapply(all.res.1B, "[[", "inf.mean.age"))
      
    ##note that from here on we will only add comments for new pieces of code (which had not yet been explained above; the simulations are largely the same)
    ##1C (no variation between parents, constrained sampling):
      print("1C")
      ##run the simulation:
      all.res.1C=parLapply(cl=cl, X=1:n.sims, function(s){
        ##create base data (this time for each parent we begin with sampling the minimum age at which its offspring is observed from somewhere in its first year of life;
          ##the resuting data set comprises only one rew per parent):
        base.data=data.frame(parent.ID=as.factor(paste("m", 1:n.parents, sep=".")), age=runif(n=n.parents, min=age.range[1], max=age.range[2]/2))
        ##replicate each row in base.data as many samples are to be simulated per parent:
        base.data=base.data[rep(1:nrow(base.data), each=n.samples), ]
        ##add to the minimum offspring age per parent randomly sampled values from 0 to half the age range:
        base.data$age=round(base.data$age+runif(n=n.parents*n.samples, min=age.range[1], max=age.range[2]/2), 3)
          ##(as a result, each parent is observed for a time span covering at most half the total possible age range)
        base.data$sim.age=base.data$age-1
        m.mat=model.matrix(object=~sim.age, data=base.data)
        xx=plogis(as.vector(m.mat[, names(coefs)]%*%coefs))
        beta.par=beta.par1topar2(mu=xx, phi=phi)
        base.data$rv=rbeta(n=n, shape1=beta.par$shape1, shape2=beta.par$shape2)
        base.data$rv=beta.tr(base.data$rv)
        m=keepWarnings(glmmTMB(rv~age+(1+age||parent.ID), data=base.data, family=beta_family))
        w=m$warnings
        m=m$value
        re.sd=attr(summary(m)$varcor$cond$parent.ID, "stddev")
        BLUPs=ranef(m)$cond$parent.ID
        emp=tapply(base.data$rv, base.data$parent.ID, mean)
        inf.mean.age=tapply(base.data$age, base.data$parent.ID, mean)
        return(list(re.sd=re.sd, BLUPs=BLUPs, emp=emp, inf.mean.age=inf.mean.age, w=w))
      })
      emp.sd.1C=unlist(lapply(lapply(all.res.1C, "[[", "emp"), sd))
      mod.sd.1C=unlist(lapply(lapply(all.res.1C, "[[", "re.sd"), "[[", 1))
      mod.where.1C=unlist(lapply(lapply(all.res.1C, "[[", "BLUPs"), "[[", "(Intercept)"))
      emp.where.1C=unlist(lapply(all.res.1C, "[[", "emp"))
      mean.age.1C=unlist(lapply(all.res.1C, "[[", "inf.mean.age"))

  ##the following simulations are in principle the same as the respective previos ones, but this time parents vary in the average propensity to show the behavior
    ##2A (balanced sampling, varying parents):
      print("2A")
      base.data=as.data.frame(expand.grid(parent.ID=as.factor(paste("m", 1:n.parents, sep=".")), age=seq(age.range[1], age.range[2], length.out=n.samples)))
      base.data$sim.age=base.data$age-1
      n=nrow(base.data)
      m.mat=model.matrix(object=~sim.age, data=base.data)
      E1=rlang::env(base.data=base.data, n=n, m.mat=m.mat)
      clusterExport(cl=cl, varlist=c("base.data", "n", "m.mat"), envir=E1)
      all.res.2A=parLapply(cl=cl, X=1:n.sims, function(s){
        ##simulate parent specific propensities to diplay the behavior:
        sim.blups=rnorm(n=n.parents, mean=0, sd=parent.sd)##n.parents samples from a normal distribution with mean=0 and sd=parent.sd
        names(sim.blups)=levels(base.data$parent.ID)
        parent.props=sim.blups[as.numeric(base.data$parent.ID)]
        xx=plogis(as.vector(m.mat[, names(coefs)]%*%coefs) + parent.props)##parent.probs are added to the linear predictor
        beta.par=beta.par1topar2(mu=xx, phi=phi)
        base.data$rv=rbeta(n=n, shape1=beta.par$shape1, shape2=beta.par$shape2)
        base.data$rv=beta.tr(base.data$rv)
        m=keepWarnings(glmmTMB(rv~age+(1+age||parent.ID), data=base.data, family=beta_family))
        w=m$warnings
        m=m$value
        re.sd=attr(summary(m)$varcor$cond$parent.ID, "stddev")
        BLUPs=ranef(m)$cond$parent.ID
        emp=tapply(base.data$rv, base.data$parent.ID, mean)
        return(list(re.sd=re.sd, BLUPs=BLUPs, emp=emp, sim.blups=sim.blups, w=w))
      })
      emp.where.2A=unlist(lapply(all.res.2A, "[[", "emp"))
      mod.where.2A=unlist(lapply(lapply(all.res.2A, "[[", "BLUPs"), "[[", "(Intercept)"))
      sim.where.2A=unlist(lapply(all.res.2A, "[[", "sim.blups"))

    ##2B (random sampling, varying parents):
      print("2B")
      all.res.2B=parLapply(cl=cl, X=1:n.sims, function(s){
        base.data=data.frame(parent.ID=as.factor(rep(paste("m", 1:n.parents, sep="."), each=n.samples)), age=round(runif(n=n.samples*n.parents, min=age.range[1], max=age.range[2]), 3))
        base.data$sim.age=base.data$age-1
        m.mat=model.matrix(object=~sim.age, data=base.data)
        sim.blups=rnorm(n=n.parents, mean=0, sd=parent.sd)
        names(sim.blups)=levels(base.data$parent.ID)
        parent.props=sim.blups[as.numeric(base.data$parent.ID)]
        xx=plogis(as.vector(m.mat[, names(coefs)]%*%coefs) + parent.props)
        beta.par=beta.par1topar2(mu=xx, phi=phi)
        base.data$rv=rbeta(n=nrow(base.data), shape1=beta.par$shape1, shape2=beta.par$shape2)
        base.data$rv=beta.tr(base.data$rv)
        m=keepWarnings(glmmTMB(rv~age+(1+age||parent.ID), data=base.data, family=beta_family))
        w=m$warnings
        m=m$value
        re.sd=attr(summary(m)$varcor$cond$parent.ID, "stddev")
        BLUPs=ranef(m)$cond$parent.ID
        emp=tapply(base.data$rv, base.data$parent.ID, mean)
        #browser()
        inf.mean.age=tapply(base.data$age, base.data$parent.ID, mean)
        return(list(re.sd=re.sd, BLUPs=BLUPs, emp=emp, sim.blups=sim.blups, inf.mean.age=inf.mean.age, w=w))
      })
      emp.where.2B=unlist(lapply(all.res.2B, "[[", "emp"))
      mod.where.2B=unlist(lapply(lapply(all.res.2B, "[[", "BLUPs"), "[[", "(Intercept)"))
      sim.where.2B=unlist(lapply(all.res.2B, "[[", "sim.blups"))
      mean.age.2B=unlist(lapply(all.res.2B, "[[", "inf.mean.age"))
      col.2B=unlist(lapply(1-(mean.age.2B-min(mean.age.2B))/diff(range(mean.age.2B)), function(x){
        return(plasma(n=1, begin=x, end=x, direction=1, alpha=0.2))
      }))
    
    ##2C (random sampling, varying parents):
      print("2C")
      all.res.2C=parLapply(cl=cl, X=1:n.sims, function(s){
        base.data=data.frame(parent.ID=as.factor(paste("m", 1:n.parents, sep=".")), age=runif(n=n.parents, min=age.range[1], max=age.range[2]/2))
        base.data=base.data[rep(1:nrow(base.data), each=n.samples), ]
        base.data$age=round(base.data$age+runif(n=n.parents*n.samples, min=age.range[1], max=age.range[2]/2), 3)
        base.data$sim.age=base.data$age-1
        m.mat=model.matrix(object=~sim.age, data=base.data)
        sim.blups=rnorm(n=n.parents, mean=0, sd=parent.sd)
        names(sim.blups)=levels(base.data$parent.ID)
        parent.props=sim.blups[as.numeric(base.data$parent.ID)]
        xx=plogis(as.vector(m.mat[, names(coefs)]%*%coefs) + parent.props)
        beta.par=beta.par1topar2(mu=xx, phi=phi)
        base.data$rv=rbeta(n=n, shape1=beta.par$shape1, shape2=beta.par$shape2)
        base.data$rv=beta.tr(base.data$rv)
        m=keepWarnings(glmmTMB(rv~age+(1+age||parent.ID), data=base.data, family=beta_family))
        w=m$warnings
        m=m$value
        re.sd=attr(summary(m)$varcor$cond$parent.ID, "stddev")
        BLUPs=ranef(m)$cond$parent.ID
        emp=tapply(base.data$rv, base.data$parent.ID, mean)
        inf.mean.age=tapply(base.data$age, base.data$parent.ID, mean)
        return(list(re.sd=re.sd, BLUPs=BLUPs, emp=emp, inf.mean.age=inf.mean.age, sim.blups=sim.blups, w=w))
      })
      emp.where.2C=unlist(lapply(all.res.2C, "[[", "emp"))
      mod.where.2C=unlist(lapply(lapply(all.res.2C, "[[", "BLUPs"), "[[", "(Intercept)"))
      sim.where.2C=unlist(lapply(all.res.2C, "[[", "sim.blups"))
      mean.age.2C=unlist(lapply(all.res.2C, "[[", "inf.mean.age"))
      col.2C=unlist(lapply(1-(mean.age.2C-min(mean.age.2C))/diff(range(mean.age.2C)), function(x){
        return(plasma(n=1, begin=x, end=x, direction=-1, alpha=0.2))
      }))
  save(file=paste(c(save.path, "/NP", n.parents, "_NS", n.samples, "_PHI", phi, ".RData"), collapse=""), 
    list=c("all.res.1A", "all.res.1B", "all.res.1C", "all.res.2A", "all.res.2B", "all.res.2C", "col.2B", "col.2C", "emp.sd.1A", "emp.sd.1B", "emp.sd.1C",
      "emp.where.1B", "emp.where.1C", "emp.where.2A", "emp.where.2B", "emp.where.2C", "mean.age.1B", "mean.age.1C", "mean.age.2B", "mean.age.2C", 
      "mod.sd.1A", "mod.sd.1B", "mod.sd.1C", "mod.where.1B", "mod.where.1C", "mod.where.2A", "mod.where.2B", "mod.where.2C", 
      "sim.where.2A", "sim.where.2B", "sim.where.2C", "phi", "n.parents", "n.samples"))
}  

library(parallel)
cl <- makeCluster(spec=120)
parLapply(cl=cl, X=1:length(cl), function(x){
  library(glmmTMB)
  library(kyotil)
  library(viridisLite)
})
n.sims=1000
for(n.parents in rev(c(20, 80))){
  for(n.samples in rev(c(21, 81))){
    for(phi in c(5, 10, 20, 40)){
      rm(list=setdiff(ls(), c("sim.fun", "phi", "cl", "beta.par1topar2", "beta.tr", "age.range", "coefs", "mo.icpt", "n.sims", "parent.sd", "save.path", "n.parents", "n.samples")))
      sim.fun(phi, n.parents=n.parents, n.samples=n.samples, n.sims=n.sims)
    }
  }
}
stopCluster(cl=cl)
#######################################################################################################################################
#######################################################################################################################################
#######################################################################################################################################
#######################################################################################################################################
#######################################################################################################################################
##SECTION 2: summarize the simulation results into several lists

setwd("")##the working directory has to be set to where the workspaces are stored that resulted from the simulation
##the follwiwng summarizes results such that plotting will be relatively fast:
  ##for the first part of evaluation (null-scenario, i.e., no variation between parents):
    xthresh=0.01##a parameter not evaluated anymore (but kept because otherwise some of the subequent code might not work)
    ##list all the workspaces that resulted from the simulation in a vector:
    all.ws=list.files(full.names=T, recursive=F)
    ##rearrange that vector into a list with one vector per each combination of number parents and number samples per parent:
    all.ws=list(
      all.ws[grepl(x=all.ws, pattern="NP20_NS21", fixed=T)],
      all.ws[grepl(x=all.ws, pattern="NP20_NS81", fixed=T)],
      all.ws[grepl(x=all.ws, pattern="NP80_NS21", fixed=T)],
      all.ws[grepl(x=all.ws, pattern="NP80_NS81", fixed=T)]
    )
    names(all.ws)=c("NP20_NS21", "NP20_NS81", "NP80_NS21", "NP80_NS81")##name the vector according to number parents and number samples per parent
    
    all.summary=lapply(all.ws, function(x){##go through each element of the list all.ws (i.e., for each cobination of number parents and number samples per parent)
      res=lapply(x, function(y){##go through each of the names of the workspaces in a given vector in all.ws (i.e., for each value of phi)
        load(y)##load the resp. workspace
        ##extract conventional 'estimates' of parent specific (just the average per parent) for null-scenario simulations:
        emp.est.1A=lapply(all.res.1A, "[[", "emp")
        emp.est.1B=lapply(all.res.1B, "[[", "emp")
        emp.est.1C=lapply(all.res.1C, "[[", "emp")
        ##transform them to link space (to make then comparable to BLUPs):
        emp.est.1A=lapply(emp.est.1A, qlogis)
        emp.est.1B=lapply(emp.est.1B, qlogis)
        emp.est.1C=lapply(emp.est.1C, qlogis)
        ##center them to a mean of zero:
        emp.est.1A=lapply(emp.est.1A, function(x){
          as.vector(scale(x, scale=F))
        })
        emp.est.1B=lapply(emp.est.1B, function(x){
          as.vector(scale(x, scale=F))
        })
        emp.est.1C=lapply(emp.est.1C, function(x){
          as.vector(scale(x, scale=F))
        })
        ##extract standard deviations of conventional estimates of parental investment:
        emp.sd.1A=lapply(emp.est.1A, sd)
        emp.sd.1B=lapply(emp.est.1B, sd)
        emp.sd.1C=lapply(emp.est.1C, sd)
        ##note that all the above extractions resulted in lists with one vector for each simulated data set
        ##extract modelled estimates of parent specific investment:
        #mod.est.2A=unlist(lapply(lapply(all.res.2A, "[[", "BLUPs"), "[[", "(Intercept)"))
        #mod.est.2B=unlist(lapply(lapply(all.res.2A, "[[", "BLUPs"), "[[", "(Intercept)"))
        #mod.est.2C=unlist(lapply(lapply(all.res.2A, "[[", "BLUPs"), "[[", "(Intercept)"))
        ##contruct the list to be returned:
        return(list(
          ylim=range(c(unlist(emp.sd.1A), mod.sd.1A, unlist(emp.sd.1B), mod.sd.1B, unlist(emp.sd.1C), mod.sd.1C)),##y-axis limits neededd for plots
          all.zeros=c("1A"=mean(mod.sd.1A<=xthresh), "1B"=mean(mod.sd.1B<=xthresh), "1C"=mean(mod.sd.1C<=xthresh)),##not evaluated anymore
          ##averages of modelled standard deviation of parental investment per sampling scenario (somehow needed for plotting):
          all.mod.means=c("1A"=mean(mod.sd.1A), "1B"=mean(mod.sd.1B), "1C"=mean(mod.sd.1C)),
          ##averages of conventinoal estimated standard deviation of parental investment per sampling scenario (somehow needed for plotting):
          all.emp.means=c("1A"=mean(unlist(emp.sd.1A)), "1B"=mean(unlist(emp.sd.1B)), "1C"=mean(unlist(emp.sd.1C))),
          n.obs=n.samples,##numper observations per parent (are existing in the resp. workspace loaded)
          n.inf=n.parents,##number parents (are existing in the resp. workspace loaded)
          ##the following three are the conventional estimates of the standard deviation of parent specific average investments (are just unlisted here
            ##to reveal vectors with one sd per each simulated data set):
          emp.sd.1A=unlist(emp.sd.1A),
          emp.sd.1B=unlist(emp.sd.1B),
          emp.sd.1C=unlist(emp.sd.1C),
          ##the following three are the modelled estimates of the standard deviation of parent specific average investments (were in the resp. workspace loaded)
          mod.sd.1A=unlist(mod.sd.1A),
          mod.sd.1B=unlist(mod.sd.1B),
          mod.sd.1C=unlist(mod.sd.1C),
          ##the following three are conventional estimates of parental investment extracted above (are just returned as named parts of the list):
          emp.est.1A=emp.est.1A,
          emp.est.1B=emp.est.1B,
          emp.est.1C=emp.est.1C,
          ##mean age per offspring in simulations with random or constrained sampling (are in the workspace loaded):
          mean.age.1B=mean.age.1B,
          mean.age.1C=mean.age.1C,
          ##modelled parent specific estimated of their investment (are in the workspace loaded):
          mod.est.1B=mod.where.1B,
          mod.est.1C=mod.where.1C
        ))
      })
      ##just a sanity check:
        n.obs=unique(unlist(lapply(res, "[[", "n.obs")))##determine unique number of observations per parent
        n.inf=unique(unlist(lapply(res, "[[", "n.inf")))##determine unique number of parents
        if(length(n.obs)>1 | length(n.inf)>1){stop("sample size problem")}
      ##extract the simulated value of phi:
      xx=strsplit(x, split="_", fixed=T)
      xx=matrix(unlist(xx), nrow=length(xx), byrow=T)
      xx=xx[, ncol(xx)]
      xx=gsub(xx, pattern="PHI", replacement="", fixed=T)
      xx=gsub(xx, pattern=".RData", replacement="", fixed=T)
      names(res)=paste("phi", xx, sep="_")##...and and name the list created by the inner lapply with the values of phi
      ##return the resulting list:
      return(res)
    })
    names(all.summary)=c("20_inf_21_obs", "20_inf_81_obs", "80_inf_21_obs", "80_inf_81_obs")
    save(file="./new_result_summary/all_summary.RData", list="all.summary")

    ##the following summarizes the simulations in which we simulated variation in parental investment
    ##(the lapply extracts the simulated modelled, and conventional estimates of parent specific investments):
    all.summary.diff=lapply(all.ws, function(x){
      res=lapply(x, function(y){
        load(y)
        ##extract conventional 'estimates' of parent specific inestment (just the average per parent) for null-scenario simulations:
        emp.est.2A=lapply(all.res.2A, "[[", "emp")
        emp.est.2B=lapply(all.res.2B, "[[", "emp")
        emp.est.2C=lapply(all.res.2C, "[[", "emp")
        ##transform them to link space (to make then comparable to BLUPs):
        emp.est.2A=lapply(emp.est.2A, qlogis)
        emp.est.2B=lapply(emp.est.2B, qlogis)
        emp.est.2C=lapply(emp.est.2C, qlogis)
        ##center them to a mean of zero:
        emp.est.2A=lapply(emp.est.2A, function(x){
          as.vector(scale(x, scale=F))
        })
        emp.est.2B=lapply(emp.est.2B, function(x){
          as.vector(scale(x, scale=F))
        })
        emp.est.2C=lapply(emp.est.2C, function(x){
          as.vector(scale(x, scale=F))
        })
        ##extract modelled estimates of parent specific investment:
        mod.est.2A=unlist(lapply(lapply(all.res.2A, "[[", "BLUPs"), "[[", "(Intercept)"))
        mod.est.2B=unlist(lapply(lapply(all.res.2A, "[[", "BLUPs"), "[[", "(Intercept)"))
        mod.est.2C=unlist(lapply(lapply(all.res.2A, "[[", "BLUPs"), "[[", "(Intercept)"))
        ##extract simulated parent specific investments:
        sim.blups.2A=unlist(lapply(all.res.2A, "[[", "sim.blups"))
        sim.blups.2B=unlist(lapply(all.res.2B, "[[", "sim.blups"))
        sim.blups.2C=unlist(lapply(all.res.2C, "[[", "sim.blups"))
        ##return the resulting lists and vectors in a named list
        return(list(
          n.obs=n.samples,
          n.inf=n.parents,
          sim.blups.2A=sim.blups.2A,
          sim.blups.2B=sim.blups.2B,
          sim.blups.2C=sim.blups.2C,
          emp.est.2A=emp.est.2A,
          emp.est.2B=emp.est.2B,
          emp.est.2C=emp.est.2C,
          mean.age.2B=mean.age.2B,
          mean.age.2C=mean.age.2C,
          mod.est.2A=mod.where.2A,
          mod.est.2B=mod.where.2B,
          mod.est.2C=mod.where.2C
        ))
      })
      ##same sanity checks as above:
        n.obs=unique(unlist(lapply(res, "[[", "n.obs")))
        n.inf=unique(unlist(lapply(res, "[[", "n.inf")))
        if(length(n.obs)>1 | length(n.inf)>1){stop("sample size probelm")}
      ##extract the simulated value of phi:
      xx=strsplit(x, split="_", fixed=T)
      xx=matrix(unlist(xx), nrow=length(xx), byrow=T)
      xx=xx[, ncol(xx)]
      xx=gsub(xx, pattern="PHI", replacement="", fixed=T)
      xx=gsub(xx, pattern=".RData", replacement="", fixed=T)
      names(res)=paste("phi", xx, sep="_")##...and and name the list created by the inner lapply with the values of phi
      return(res)
    })
    names(all.summary.diff)=c("20_inf_21_obs", "20_inf_81_obs", "80_inf_21_obs", "80_inf_81_obs")
    save(file="./new_result_summary/all_summary_diff.RData", list="all.summary.diff")

    ##create summary object as needed for plotting estimated against simulated BLUPs for balanced simulations (basically just rearranges the existing list all.summary.diff):
    phi=c(5, 10, 20, 40)
    all.sim.res.bal.with.diff=lapply(phi, function(p){
      lapply(all.summary.diff, function(x){
        x[[paste(c("phi", p), collapse="_")]]
      })
    })
    names(all.sim.res.bal.with.diff)=paste("phi", phi, sep="_")
    save(file="./new_result_summary/all_sim_res_bal_with_diff.Rdata", list="all.sim.res.bal.with.diff")
    
    ##the following summarizes the simulations in which we simulated variation in parental investment
    ##(the lapply extracts the simulated modelled, and conventional estimates of parent specific investments,
      ##and also the mean age per each offspring):
    all.summary.2BC=lapply(all.ws, function(x){
      res=lapply(x, function(y){
        load(y)
        #browser()
        return(list(mean.age.2B=as.vector(unlist(mean.age.2B)),
                    mean.age.2C=as.vector(unlist(mean.age.2C)),
                    sim.where.2B=as.vector(unlist(sim.where.2B)),
                    sim.where.2C=as.vector(unlist(sim.where.2C)),
                    mod.where.2B=as.vector(unlist(mod.where.2B)),
                    mod.where.2C=as.vector(unlist(mod.where.2C)),
                    emp.where.2B=as.vector(qlogis(unlist(emp.where.2B))),
                    emp.where.2C=as.vector(qlogis(unlist(emp.where.2C)))
        ))
      })
      ##extract the simulated value of phi:
      xx=strsplit(x, split="_", fixed=T)
      xx=matrix(unlist(xx), nrow=length(xx), byrow=T)
      xx=xx[, ncol(xx)]
      xx=gsub(xx, pattern="PHI", replacement="", fixed=T)
      xx=gsub(xx, pattern=".RData", replacement="", fixed=T)
      names(res)=paste("phi", xx, sep="_")##...and and name the list created by the inner lapply with the values of phi
      return(res)
    })
    names(all.summary.2BC)=c("20_inf_21_obs", "20_inf_81_obs", "80_inf_21_obs", "80_inf_81_obs")
    save(file="./new_result_summary/all_summary_2BC.Rdata", list="all.summary.2BC")

    
#######################################################################################################################################
#######################################################################################################################################
#######################################################################################################################################
#######################################################################################################################################
#######################################################################################################################################
##SECTION 3: code for creating the Figures with results shown in the main text
##note that the saving of the Figures will only work when there is a folder named FIGs in the path sat as save.path above
  violin.fun<-function(x, y, scale.fac=1, resol=0.1, ylim=NULL, xlab, ylab, col="grey"){
    breaks=seq(0, max(y), by=resol)
    breaks=c(breaks, max(breaks+resol))
    bin.y=cut(abs(y), breaks=breaks, include.lowest=T, label=F)
    bin.y=(resol/2+(bin.y-1)*resol)*sign(y)
    xx=aggregate(x=bin.y, by=list(x=x, y=bin.y), length)
    colnames(xx)[ncol(xx)]="freq"
    xx$freq=xx$freq/max(xx$freq)
    if(is.null(ylim)){ylim=range(y)}
    #par(mar=c(3, 3, 0.2, 0.2), mgp=c(1.7, 0.3, 0), tcl=-0.15, las=1)
    plot(1, 1, xlim=c(0.5, length(unique(xx$x))+0.5), ylim=ylim,
      xlab=xlab, ylab=ylab, xaxt="n", type="n", yaxt="n")
    rect(xleft=xx$x-xx$freq*scale.fac/2, xx$x+xx$freq*scale.fac/2, ybottom=xx$y-resol/2, ytop=xx$y+resol/2, border=NA, col=col)
  }
  plot.bin.fun<-function(x, y, x.bin.w, y.bin.w){
    breaks=seq(0, max(abs(x))+x.bin.w, by=x.bin.w)
    binned.x=cut(abs(x), breaks, labels=F, include.lowest=T)
    binned.x=(x.bin.w/2 + (binned.x-1)*x.bin.w)*sign(x)
    breaks=seq(0, max(abs(y))+y.bin.w, by=y.bin.w)
    binned.y=cut(abs(y), breaks=breaks, labels=F, include.lowest=T)
    binned.y=(y.bin.w/2 + (binned.y-1)*y.bin.w)*sign(y)
    to.plot=aggregate(1:length(x), by=data.frame(x=binned.x, y=binned.y), length)
    names(to.plot)[ncol(to.plot)]="freq"
    return(to.plot)
  }

    ##determine y-axis limits:
    all.ylim=lapply(all.summary, function(x){
      range(unlist(lapply(x, "[[", "ylim")))
    })##but as these appeared to not work well, we don't use it´
    all.phi=paste("phi", c(5, 10, 20, 40), sep="_")
    this.ylim=lapply(all.summary, function(x){
      lapply(x, function(y){
        range(c(y$all.emp.means, y$all.mod.means))
      })
    })
    this.ylim=range(unlist(this.ylim))
    ##set point characters etc.:
    all.pch=c(1, 22, 4)
    cexes=c(1.5, 2, 1)*1
    cex.lab=1.7
    cex.axis.lab=1.9
    cex.text=1.25
    lmat=rbind(c(9, 7, 7), cbind(5, matrix(1:4, ncol=2, byrow=T)), c(8, 6, 6))

    dev.off()
    X11(width=10)##this will only work on Linux computers on Mac and Windows, replace by quartz and windows respectively
    layout(lmat, widths=c(1, rep(6, 2)), heights=c(2, rep(6, 2), 1))##set up split plotting window
    par(mar=c(0.5, 0.5, 0.2, 0.2), mgp=c(1.7, 0.3, 0), tcl=-0.2, las=0, cex.axis=1.5)##define graphical parameters
    for(n.inf in c(20, 80)){##for each number of parents
      for(n.obs in c(21, 81)){##for each number of observations per parent
        y=all.summary[[paste(c(n.inf, "inf", n.obs, "obs"), collapse="_")]]##extract relevant part of the list wrt number of parents and number of observations per parent
        xx=lapply(y[all.phi], "[[", "all.mod.means")##extract modelled results per value of phi
        xx=matrix(unlist(xx), nrow=length(xx), byrow=T)##restruture object obtained
        ##begin with plot:
        plot(x=1, 1, type="n", xlim=c(0.5, 4.5), xaxs="i", xaxt="n", ylim=c(0, max(this.ylim))*c(1, 1.1), yaxt="n", xlab="", ylab="")
        ##add points (and lines):
        for(i in 1:3){
          points(x=1:4, y=xx[, i], type="b", pch=all.pch[i], cex=cexes[i], lty=3)
          points(x=1:4, y=xx[, i], pch=all.pch[i], cex=cexes[i], lwd=1)
        }
        xx=lapply(y[all.phi], "[[", "all.emp.means")##extract conventionally estimated results per value of phi (and add to plot as above):
        xx=matrix(unlist(xx), nrow=length(xx), byrow=T)
        for(i in 1:3){
          points(x=1:4, y=xx[, i], type="b", pch=all.pch[i], cex=cexes[i], lty=2)
          points(x=1:4, y=xx[, i], pch=all.pch[i], cex=cexes[i], lwd=1)
        }
        ##add axies where needed:
        if(n.inf==80){
          axis(side=1, at=1:4, labels=c(5, 10, 20, 40))
        }else{
          axis(side=1, at=1:4, labels=NA)
        }
        if(n.obs==21){
          axis(side=2, at=seq(0, 0.8, by=0.1))
        }else{
          axis(side=2, at=seq(0, 0.8, by=0.1), labels=NA)
        }
        ##add margin text on top of plot:
        mtext(text=paste(c(n.inf, " offspring, ", n.obs, "observations"), collapse=""), side=3, line=-1.5, cex=cex.text)
      }
    }
    ##add legend:
    par(mar=c(0.5, 0.2, 0.2, 0.2), mgp=c(1.7, 0.3, 0), tcl=-0.2, las=0)
    plot(1, 1, type="n", axes=F, ann=F)
    text(labels="average standard deviation", x=0.8, y=1, cex=cex.axis.lab, srt=90)
    par(mar=c(0.2, 0.5, 0.2, 0.2), mgp=c(1.7, 0.3, 0), tcl=-0.2, las=0)
    plot(1, 1, type="n", axes=F, ann=F)
    text(labels="phi", x=1, y=1, cex=cex.axis.lab)

    par(mar=c(0.1, 0.1, 0.2, 0.2), mgp=c(1.7, 0.3, 0), tcl=-0.2, las=0)
    plot(1, 1, type="n", axes=F, ann=F, xlim=c(0, 1.2), ylim=c(-0.2, 1.2))
    x=c(0.4, 0.6, 0.8)
    y=c(0.5, 0.2)
    hll=0.05
    text(labels=c("balanced", "random", "constrained"), x=x, y=rep(0.8, 3), cex=cex.lab)
    text(labels=c("estimation"), x=0.2, y=0.8, cex=cex.lab)
    text(labels=c("conventional", "modeled"), x=rep(0.2, 2), y=y, cex=cex.lab)
    segments(x0=rep(x, each=2)-hll, x1=rep(x, each=2)+hll, y0=rep(y, times=2), y1=rep(y, times=2), lty=rep(2:3, times=2))
    points(x=rep(x, each=2), y=rep(y, times=3), pch=19, col="white", cex=2)
    points(x=rep(x, each=2), y=rep(y, times=3), pch=rep(all.pch, each=2), cex=rep(cexes, each=2), lwd=1)
    savePlot(file="./FIGS/FIG_6_sd_null_scen.png", type="png")
    dev.copy2pdf(file="./FIGS/FIG_6_sd_null_scen.pdf")

    ##################################################
    ##Fig. 7, main text (R2 as a function of sample sixes and phi):
    ##extract R2:
    all.R2=lapply(all.summary, function(x){
      lapply(x, function(y){
        meB=lm(unlist(y$emp.est.1B)~y$mean.age.1B)
        meC=lm(unlist(y$emp.est.1C)~y$mean.age.1C)
        mmB=lm(unlist(y$mod.est.1B)~y$mean.age.1B)
        mmC=lm(unlist(y$mod.est.1C)~y$mean.age.1C)
        return(c(meB=summary(meB)$r.squared, meC=summary(meC)$r.squared, mmB=summary(mmB)$r.squared, mmC=summary(mmC)$r.squared))
      })
    })
    ##determine y-axis limits:
    all.ylim=lapply(all.R2, function(x){
      lapply(x, function(y){
        range(y)
      })
    })
    all.ylim=range(unlist(all.ylim))
    all.phi=paste("phi", c(5, 10, 20, 40), sep="_")
    ##set out layout matrix (for the splitting of the plotting window):
    lmat=rbind(cbind(5, matrix(1:4, ncol=2, byrow=T)), c(8, 6, 6))
    lmat=rbind(c(9, 7, 7), lmat)
    ##general plotting parameters:
    all.pch=rep(c(22, 4), times=2)
    all.cex=rep(c(2, 1.5), times=2)*1
    cex.axes1=1.5
    cex.text=1
    cex.legend=1.5

    dev.off()
    X11(height=5)##this will only work on Linux computers on Mac and Windows, replace by quartz and windows respectively
    layout(lmat, widths=c(1, rep(7, 2)), heights=c(2.5, rep(7, 2), 1))##set up plotting window
    par(mar=c(0.5, 0.5, 0.2, 0.2), mgp=c(1.7, 0.3, 0), tcl=-0.2, las=0)##set graphical parameters
    for(n.inf in c(20, 80)){##for each number parents
      for(n.obs in c(21, 81)){##for each sample size
        y=all.R2[[paste(c(n.inf, "inf", n.obs, "obs"), collapse="_")]]##extract R2
        ##reformat R2:
        xnames=names(y)
        y=matrix(unlist(y), nrow=length(y), byrow=T)
        rownames(y)=xnames
        y=y[all.phi, ]
        ##begin with plot:
        plot(x=1, 1, type="n", xlim=c(0.5, 4.5), xaxs="i", xaxt="n", ylim=c(0, 1)*c(1, 1.1), yaxt="n", xlab="", ylab="")
        ##add points and lines:
        pch=rep(c(3, 4), times=2)
        lty=rep(2:3, each=2)
        for(i in 1:4){
          points(x=1:4, y=y[, i], type="b", pch=all.pch[i], lty=lty[i], cex=all.cex[i])
          points(x=1:4, y=y[, i], lwd=1, pch=all.pch[i], lty=lty[i], cex=all.cex[i])
        }
        ##add axes:
        if(n.inf==80){
          axis(side=1, at=1:4, labels=c(5, 10, 20, 40))
        }else{
          axis(side=1, at=1:4, labels=NA)
        }
        if(n.obs==21){
          axis(side=2, at=seq(0, 1, by=0.2))
        }else{
          axis(side=2, at=seq(0, 1, by=0.2), labels=NA)
        }
        ##add top pargin text indicating sample sizes:
        mtext(text=paste(c(n.inf, " offspring, ", n.obs, "observations"), collapse=""), side=3, line=-1.7, cex=cex.text)
      }
    }
    ##add legend etc.:
    par(mar=c(0.5, 0.2, 0.2, 0.2), mgp=c(1.7, 0.3, 0), tcl=-0.2, las=0)
    plot(1, 1, type="n", axes=F, ann=F)
    text(labels=bquote(R^2), x=0.8, y=1, cex=cex.axes1, srt=90)
    par(mar=c(0.2, 0.5, 0.2, 0.2), mgp=c(1.7, 0.3, 0), tcl=-0.2, las=0)
    plot(1, 1, type="n", axes=F, ann=F)
    text(labels="phi", x=1, y=1, cex=cex.axes1)
    #legend:
    par(mar=c(0.1, 0.5, 0.1, 0.2), mgp=c(1.7, 0.3, 0), tcl=-0.2, las=0)
    plot(1, 1, type="n", axes=F, ann=F, xlim=c(0, 1), ylim=c(-0.2, 1.2))
    x=c(0.4, 0.6)
    y=c(0.5, 0.2)
    hll=0.05
    text(labels=c("random", "constrained"), x=x, y=rep(0.8, 2), cex=cex.legend)
    text(labels=c("estimation"), x=0.2, y=0.8, cex=cex.legend)
    text(labels=c("conventional", "modeled"), x=rep(0.2, 2), y=y, cex=cex.legend)
    segments(x0=rep(x, each=2)-hll, x1=rep(x, each=2)+hll, y0=rep(y, times=2), y1=rep(y, times=2), lty=rep(2:3, times=2))
    points(x=rep(x, each=2), y=rep(y, times=2), pch=19, col="white", cex=2*1.5)
    points(x=rep(x, each=2), y=rep(y, times=2), pch=rep(all.pch[1:2], each=2), cex=rep(all.cex[1:2], each=2), lwd=1)
    dev.copy2pdf(file="./FIGS/FIG_7_age_R2_null.pdf")
    savePlot(file="./FIGS/FIG_7_age_R2_null.png", type="png")

    ############################################################################################
    ##Fig. 8 of main text:
    ##extract partial R2 of BLUPs (modelled and conventionally estimated) against simulated parent specific investment:
    allR2.2=lapply(all.summary.diff, function(x){
      lapply(x, function(y){
        ##fit regressions of estimated against simulated parent specific investment and extract R2 (balanced sampling):
        est.2A=summary(lm(unlist(y$emp.est.2A)~y$sim.blups.2A))$r.squared
        mod.2A=summary(lm(y$mod.est.2A~y$sim.blups.2A))$r.squared
        names(est.2A)=names(mod.2A)="BLUPs"
        ##fit regression of conventionally estimated parent specidif investment against simulated parent specific investment and average offsrping age and extract R2 (random sampling):
        m=lm(unlist(y$emp.est.2B)~y$sim.blups.2B+y$mean.age.2B)
        m=as.data.frame(drop1(m))$RSS
        m=(m[-1]-m[1])/m[-1]
        names(m)=c("BLUPs", "AGE")
        est.2B=m
        ##fit regression of modelled parent specidif investment against simulated parent specific investment and average offsrping age and extract R2 (random sampling):
        m=lm(y$mod.est.2B~y$sim.blups.2B+y$mean.age.2B)
        m=as.data.frame(drop1(m))$RSS
        m=(m[-1]-m[1])/m[-1]
        names(m)=c("BLUPs", "AGE")
        mod.2B=m
        ##fit regression of conventionally estimated parent specidif investment against simulated parent specific investment and average offsrping age and extract R2 (constrained sampling):
        m=lm(unlist(y$emp.est.2C)~y$sim.blups.2C+y$mean.age.2C)
        m=as.data.frame(drop1(m))$RSS
        m=(m[-1]-m[1])/m[-1]
        names(m)=c("BLUPs", "AGE")
        est.2C=m
        ##fit regression of modelled parent specidif investment against simulated parent specific investment and average offsrping age and extract R2 (constrained sampling):
        m=lm(y$mod.est.2C~y$sim.blups.2C+y$mean.age.2C)
        m=as.data.frame(drop1(m))$RSS
        m=(m[-1]-m[1])/m[-1]
        names(m)=c("BLUPs", "AGE")
        mod.2C=m
        return(list(est.2A=est.2A, est.2B=est.2B, est.2C=est.2C, mod.2A=mod.2A, mod.2B=mod.2B, mod.2C=mod.2C))
      })
    })
    ##just extract the (partial) R2 against simulated parent specific investment:
    all.BLUPs.R2=lapply(allR2.2, function(x){
      lapply(x, function(y){
        lapply(y, function(z){
          #browser()
          z["BLUPs"]
        })
      })
    })
    ##determine y-axis limits:
    all.ylim=range(unlist(all.BLUPs.R2))##eventually discarded
    all.ylim=c(0.7, 1)
    ##set up lalyout matrix for splitting up of plot window:
    lmat=rbind(cbind(5, matrix(1:4, ncol=2, byrow=T)), c(8, 6, 6))
    lmat=rbind(c(9, 7, 7), lmat)
    ##some general parameters etc.:
    all.phi=paste("phi", c(5, 10, 20, 40), sep="_")
    all.pch=rep(c(1, 22, 4), times=2)
    cexes=rep(c(1.5, 2, 1), times=2)*1.5
    cex.text=1
    cex.legend=1.5
    cex.lab=1.8

    dev.off()
    X11(height=5)##this will only work on Linux computers on Mac and Windows, replace by quartz and windows respectively
    ##set up plotting window:
    layout(lmat, widths=c(1, rep(6, 2)), heights=c(3, rep(6, 2), 1))
    par(mar=c(0.5, 0.5, 0.2, 0.2), mgp=c(1.7, 0.3, 0), tcl=-0.2, las=0, cex.axis=1.5)##general graphical parameters
    for(n.inf in c(20, 80)){##for each number parents
      for(n.obs in c(21, 81)){##for each sample size
        y=all.BLUPs.R2[[paste(c(n.inf, "inf", n.obs, "obs"), collapse="_")]]##extract relevant R2...
        ##... and reformat them:
        xnames=names(y)
        y=matrix(unlist(y), nrow=length(y), byrow=T)
        rownames(y)=xnames
        y=y[all.phi, ]
        ##begin with plot:
        plot(x=1, 1, type="n", xlim=c(0.5, 4.5), xaxs="i", xaxt="n", ylim=all.ylim*c(1, 1.025), yaxt="n", xlab="", ylab="")
        lty=rep(2:3, each=3)
        ##add points (and lines):
        for(i in 1:6){
          points(x=1:4, y=y[, i], type="b", lty=lty[i], pch=all.pch[i], cex=cexes[i])
          points(x=1:4, y=y[, i], lty=lty[i], pch=all.pch[i], cex=cexes[i], lwd=1) 
        }
        ##add axes:
        if(n.inf==80){
          axis(side=1, at=1:4, labels=c(5, 10, 20, 40))
        }else{
          axis(side=1, at=1:4, labels=NA)
        }
        if(n.obs==21){
          axis(side=2, at=seq(0.7, 1, by=0.1))
        }else{
          axis(side=2, at=seq(0.7, 1, by=0.1), labels=NA)
        }
        ##add text with sample sizes:
        mtext(text=paste(c(n.inf, " offspring, ", n.obs, "observations"), collapse=""), side=3, line=-1.8, cex=cex.text)
      }
    }
    ##add legend etc.:
    par(mar=c(0.5, 0.2, 0.2, 0.2), mgp=c(1.7, 0.3, 0), tcl=-0.2, las=0)
    plot(1, 1, type="n", axes=F, ann=F)
    text(labels=bquote(R^2), x=0.8, y=1, cex=cex.lab, srt=90)
    par(mar=c(0.2, 0.5, 0.2, 0.2), mgp=c(1.7, 0.3, 0), tcl=-0.2, las=0)
    plot(1, 1, type="n", axes=F, ann=F)
    text(labels="phi", x=1, y=1, cex=cex.lab)
    par(mar=c(0.1, 0.5, 0.1, 0.2), mgp=c(1.7, 0.3, 0), tcl=-0.2, las=0)
    plot(1, 1, type="n", axes=F, ann=F, xlim=c(0, 1), ylim=c(-0.2, 1.2))
    x=c(0.4, 0.6, 0.8)
    y=c(0.5, 0.2)
    hll=0.05
    text(labels=c("balanced", "random", "constrained"), x=x, y=rep(0.8, 3), cex=cex.legend)
    text(labels=c("estimation"), x=0.2, y=0.8, cex=cex.legend)
    text(labels=c("conventional", "modeled"), x=rep(0.2, 2), y=y, cex=cex.legend)
    segments(x0=rep(x, each=2)-hll, x1=rep(x, each=2)+hll, y0=rep(y, times=2), y1=rep(y, times=2), lty=rep(2:3, times=2))
    points(x=rep(x, each=2), y=rep(y, times=3), pch=19, col="white", cex=2.5)
    points(x=rep(x, each=2), y=rep(y, times=3), pch=rep(all.pch[1:3], each=2), cex=rep(cexes[1:3], each=2), lwd=1)
    dev.copy2pdf(file="./FIGS/FIG_8_parent_varying_sim_and_est_R2.pdf")
    savePlot(file="./FIGS/FIG_8_parent_varying_sim_and_est_R2.png", type="png")

    ########################################################################################
    ##Fig. 9 of the main text (age and estimate, one example):
    n.obs=21##number observations per parent (can be 21 or 81; for the main text we chose 21)
    n.inf=80##number parents (can be 20 or 80; for the main text we chose 80)
    phi=10##precision parameter (can be 5, 19, 20, or 40; ; for the main text we chose 10)
    ##set up layout matrix (i.e., splitting of plot window)
    lmat=matrix(1:4, ncol=2, byrow=T)
    lmat=cbind(c(rep(6, 2), NA), rbind(lmat, 5))
    lmat=rbind(c(NA, 7, 8), lmat)
    lmat=cbind(c(NA, 9, 10, NA), lmat)
    lmat[is.na(lmat)]=max(lmat, na.rm=T)+(1:sum(is.na(lmat)))
    ##extract relevant output wrt number parents, number observation per parent and precision parameter:
    to.plot=all.summary.diff[[paste(c(n.inf, "inf", n.obs, "obs"), collapse="_")]][[paste(c("phi", phi), collapse="_")]]
    ##extract simulated and modelled BLUPs (note that the simulated and properly modelled ones are already unlisted, so we just select the first n.inf ones):
    simB=to.plot$sim.blups.2B[1:n.inf]
    simC=to.plot$sim.blups.2C[1:n.inf]
    empB=to.plot$emp.est.2B[[1]]
    empC=to.plot$emp.est.2C[[1]]
    modB=to.plot$mod.est.2B[1:n.inf]
    modC=to.plot$mod.est.2C[1:n.inf]
    ##extract simulated offspring age (are already unlisted, so we just select the first n.inf ones):
    ageB=as.vector(to.plot$mean.age.2B[1:n.inf])
    ageC=as.vector(to.plot$mean.age.2C[1:n.inf])
    ##define the coloration of points in scale ranging from minimum to maximum age):
    age.range=range(c(ageB, ageC))
    xx.age=(ageB-min(age.range))/diff(range(age.range))
    colB=sapply(xx.age, function(x){
      plasma(n=1, begin=1-x, end=1-x, alpha=0.9)
    })
    xx.age=(ageC-min(age.range))/diff(range(age.range))
    colC=sapply(xx.age, function(x){
      plasma(n=1, begin=1-x, end=1-x, alpha=0.9)
    })
    text.cex=1.75##size of fonts

    dev.off()
    X11(height=5)##see above
    ##set up plotting window:
    layout(lmat, widths=rep(c(1, 6), each=2), heights=c(1, rep(6, 2), 1))
    layout.show(max(lmat))
    par(mar=c(0.5, 0.5, 0.2, 0.2), mgp=c(1.7, 0.3, 0), tcl=-0.15, las=1)
    ##plots:
    plot(simB, empB, xlab="", ylab="", xaxt="n", yaxt="n", xlim=range(c(simB, simC)), ylim=range(empB, modB, empC, modC), pch=19, col=colB, cex=1.5)
    axis(side=1, labels=NA)
    axis(side=2)
    plot(simB, modB, xlab="", ylab="", xaxt="n", yaxt="n", xlim=range(c(simB, simC)), ylim=range(empB, modB, empC, modC), pch=19, col=colB, cex=1.5)
    axis(side=1, labels=NA)
    axis(side=2, labels=NA)
    plot(simC, empC, xlab="", ylab="", xaxt="n", yaxt="n", xlim=range(c(simB, simC)), ylim=range(empB, modB, empC, modC), pch=19, col=colC, cex=1.5)
    axis(side=1)
    axis(side=2)
    plot(simC, modC, xlab="", ylab="", xaxt="n", yaxt="n", xlim=range(c(simB, simC)), ylim=range(empB, modB, empC, modC), pch=19, col=colC, cex=1.5)
    axis(side=1)
    axis(side=2, labels=NA)
    ##add legend:
    l.resol=250
    labs=pretty(age.range)
    labs=labs[labs>=min(age.range) &  labs<=max(age.range)]
    y=seq(from=par()$usr[3]+diff(range(par()$usr[3:4]))*0.05, to=par()$usr[3]+diff(range(par()$usr[3:4]))*0.5, length.out=l.resol)
    col=seq(from=min(age.range), to=max(age.range), length.out=l.resol)
    col=(col-min(age.range))/diff(range(age.range))
    col=sapply(col, function(x){
      plasma(n=1, begin=1-x, end=1-x, alpha=0.9)
    })
    segments(x0=par()$usr[2]-diff(range(par()$usr[1:2]))*0.2, x1=par()$usr[2]-diff(range(par()$usr[1:2]))*0.1, y0=y, y1=y, col=col, lend=1)
    text(labels=sprintf("%.1f", labs/2), x=rep(par()$usr[2]-diff(range(par()$usr[1:2]))*0.1, length(labs))-0.08, 
      y=par()$usr[3]+diff(range(par()$usr[3:4]))*0.05 + (labs-min(age.range))/diff(range(age.range))*diff(range(par()$usr[3:4]))*0.45,
      pos=4, cex=1.5)
    text(labels="mean\nage", x=mean(c(par()$usr[2]-diff(range(par()$usr[1:2]))*0.2, par()$usr[2]-diff(range(par()$usr[1:2]))*0.1+strwidth("0.0", cex=1.5))),
      y=max(y)+diff(range(par()$usr[3:4]))*0.1, cex=1.5)
    ##axis labels etc.:
    par(mar=c(0.2, 0.5, 0.2, 0.2), mgp=c(1.7, 0.3, 0), tcl=-0.15, las=1)
    plot(1, 1, axes=F, ann=F, type="n")
    text(x=1, y=0.9, labels="simulated parental behavior", cex=text.cex)
    par(mar=c(0.5, 0.2, 0.2, 0.2), mgp=c(1.7, 0.3, 0), tcl=-0.15, las=1)
    plot(1, 1, axes=F, ann=F, type="n")
    text(x=0.9, y=1, labels="estimated parental behavior", cex=text.cex, srt=90)
    par(mar=c(0.2, 0.5, 0.2, 0.2), mgp=c(1.7, 0.3, 0), tcl=-0.15, las=1)
    plot(1, 1, axes=F, ann=F, type="n")
    text(x=1, y=0.9, labels="conventional", cex=text.cex)
    plot(1, 1, axes=F, ann=F, type="n")
    text(x=1, y=0.9, labels="modeled", cex=text.cex)
    par(mar=c(0.5, 0.2, 0.2, 0.2), mgp=c(1.7, 0.3, 0), tcl=-0.15, las=1)
    plot(1, 1, axes=F, ann=F, type="n")
    text(x=0.9, y=1, labels="random sampling", cex=text.cex, srt=90)
    plot(1, 1, axes=F, ann=F, type="n")
    text(x=0.9, y=1, labels="constrained sampling", cex=text.cex, srt=90)
    dev.copy2pdf(file="./FIGS/FIG_9_age_and_est.pdf")
    savePlot(file="./FIGS/FIG_9_age_and_est.png", type="png")

    #####################################################################################################
    ##Fig. 10 of the main text (age and estimate, all R^2):
    ##extract R2 of age:
    all.AGE.R2=lapply(allR2.2, function(x){
      lapply(x, function(y){
        xx=lapply(y, function(z){
          z["AGE"]
        })
        xx=xx[!is.na(xx)]
      })
    })
    all.ylim=range(unlist(all.AGE.R2), na.rm=T)##determine ylim based on R1
    all.ylim=c(0, 1)##eventually we decided to just use this
    all.phi=paste("phi", c(5, 10, 20, 40), sep="_")##vector with names needed to extract R2 correctly
    ##set up layout matrix (i.e., the splitting of the plot window:
    lmat=rbind(cbind(5, matrix(1:4, ncol=2, byrow=T)), c(8, 6, 6))
    lmat=rbind(c(9, 7, 7), lmat)
    ##set some graphical settings:
    all.pch=rep(c(22, 4), times=2)
    cexes=rep(c(2, 1), times=2)
    cex.axes=1.7
    cex.text=1.1
    cex.legend=1.7
    
    dev.off()
    X11(height=5)##see above
    ##set up plotting window:
    layout(lmat, widths=c(1, rep(6, 2)), heights=c(3, rep(6, 2), 1))
    par(mar=c(0.5, 0.5, 0.2, 0.2), mgp=c(1.7, 0.3, 0), tcl=-0.2, las=0)
    for(n.inf in c(20, 80)){##for 10 or 80 parents
      for(n.obs in c(21, 81)){##for 21 or 81 samples per parent
        ##extract relevant R2:
        y=all.AGE.R2[[paste(c(n.inf, "inf", n.obs, "obs"), collapse="_")]]
        ##reformat y:
        xnames=names(y)
        y=matrix(unlist(y), nrow=length(y), byrow=T)
        rownames(y)=xnames
        y=y[all.phi, ]
        ##begin with plot:
        plot(x=1, 1, type="n", xlim=c(0.5, 4.5), xaxs="i", xaxt="n", ylim=all.ylim*c(1, 1.1), yaxt="n", xlab="", ylab="")
        pch=rep(c(3, 4), times=2)
        lty=rep(2:3, each=2)
        ##add data:
        for(i in 1:4){
          points(x=1:4, y=y[, i], type="b", pch=all.pch[i], lty=lty[i], cex=cexes[i])
        }
        ##add axes:
        if(n.inf==80){
          axis(side=1, at=1:4, labels=c(5, 10, 20, 40))
        }else{
          axis(side=1, at=1:4, labels=NA)
        }
        if(n.obs==21){
          axis(side=2, at=seq(0, 1, by=0.2))
        }else{
          axis(side=2, at=seq(0, 1, by=0.2), labels=NA)
        }
        ##add margin text with number parents and number observation per parent:
        mtext(text=paste(c(n.inf, " offspring, ", n.obs, "observations"), collapse=""), side=3, line=-1.3, cex=cex.text)
      }
    }
    ##axis labels etc.
    par(mar=c(0.5, 0.2, 0.2, 0.2), mgp=c(1.7, 0.3, 0), tcl=-0.2, las=0)
    plot(1, 1, type="n", axes=F, ann=F)
    text(labels=bquote(partial~R^2), x=0.8, y=1, cex=cex.axes, srt=90)
    par(mar=c(0.2, 0.5, 0.2, 0.2), mgp=c(1.7, 0.3, 0), tcl=-0.2, las=0)
    plot(1, 1, type="n", axes=F, ann=F)
    text(labels="phi", x=1, y=1, cex=cex.axes)
    par(mar=c(0.1, 0.5, 0.1, 0.2), mgp=c(1.7, 0.3, 0), tcl=-0.2, las=0)
    plot(1, 1, type="n", axes=F, ann=F, xlim=c(0, 1), ylim=c(-0.2, 1.2))
    x=c(0.4, 0.6)
    y=c(0.5, 0.2)
    hll=0.05
    text(labels=c("random", "constrained"), x=x, y=rep(0.8, 2), cex=cex.legend)
    text(labels=c("estimation"), x=0.2, y=0.8, cex=cex.legend)
    text(labels=c("conventional", "modeled"), x=rep(0.2, 2), y=y, cex=cex.legend)
    segments(x0=rep(x, each=2)-hll, x1=rep(x, each=2)+hll, y0=rep(y, times=2), y1=rep(y, times=2), lty=rep(2:3, times=2))
    points(x=rep(x, each=2), y=rep(y, times=2), pch=19, col="white", cex=2)
    points(x=rep(x, each=2), y=rep(y, times=2), pch=rep(all.pch[1:2], each=2), cex=rep(cexes[1:2], each=2))
    dev.copy2pdf(file="./FIGS/FIG_10_age_and_est_allR2.pdf")
    savePlot(file="./FIGS/FIG_10_age_and_est_allR2.png", type="png")

#######################################################################################################################################
#######################################################################################################################################
#######################################################################################################################################
#######################################################################################################################################
#######################################################################################################################################
##SECTION 4: code for creating the Figures with results shown in the Supporting information
  ##FIG. SI 1 to SI 4:
      all.phi=paste("phi", c(5, 10, 20, 40), sep="_")##vector with names to address output correectly
      ##set up layout matrix for splitting the plotting window:
      ylim=range(unlist(all.ylim))#[[paste(c(n.inf, "inf", n.obs, "obs"), collapse="_")]]#unlist(range(all.ylim)))
      lmat=matrix(1:12, nrow=3, byrow=F)
      lmat=rbind(13, 14:17, lmat, 18)
      lmat=cbind(c(NA, NA, 20:22, NA), c(NA, NA, rep(19, 3), NA), lmat)
      lmat[is.na(lmat)]=(1:sum(is.na(lmat)))+max(lmat, na.rm=T)
      ##define names of parts of summary lists to be extracted
      E=c("emp.sd.1A", "emp.sd.1B", "emp.sd.1C")
      M=c("mod.sd.1A", "mod.sd.1B", "mod.sd.1C")
      sample.m=c("balanced", "random", "constrained")
      n.obs=21##number of observations per parent (can be 21 or 81; choose to get Fig. SI 1 to SI 4)
      n.inf=20##number of parents (can be 20 or 80; choose to get Fig. SI 1 to SI 4)
      scenario="null-scenario"
      dev.off()
      X11(height=6, width=6)##see above
      ##set up plotting window and graphical parmeters:
      layout(lmat, widths=c(1, 1.5, rep(5, 4)), heights=c(1, 1, rep(5, 3), 1.5))
      par(mar=c(0.2, 0.5, 0.2, 0.2), mgp=c(2, 0.3, 0), tcl=-0.15, las=1)
      ##plot:
      for(phi in all.phi){##for all values of the precision parameter
        this.this.summary=all.summary[[paste(c(n.inf, "inf", n.obs, "obs"), collapse="_")]][[phi]]
        for(i in 1:3){##for each sampling scenario
          ##plot:
          violin.fun(x=rep(1:2, each=1000), y=c(this.this.summary[[E[i]]], this.this.summary[[M[i]]]), 
                  scale.fac=1, resol=0.01, ylim=ylim, xlab="", ylab="", col="black")
          ##add axes
          if(phi%in%all.phi[1]){
            axis(side=2)
          }else{
            axis(side=2, labels=F)
          }
          if(i==3){mtext(text=c("conventional", "modeled"), side=1, at=1:2, line=0.2, cex=0.7)}
        }
      }
      ##axis labels etc._
      par(mar=c(0.1, 0.5, 0.1, 0.2), mgp=c(2, 0.3, 0), tcl=-0.15, las=1)
      plot(1, 1, axes=F, ann=F, type="n")#xlim=c(0, 1), ylim=c(0, 1), 
      text(x=1, y=1, adj=0.5, cex=1.5,
        labels=bquote(bold(.(paste(c(n.inf, " offspring, ", n.obs, " observations per offspring, ", "null-scenario"), collapse="")))))
      for(phi in all.phi){
        plot(1, 1, axes=F, ann=F, type="n")#xlim=c(0, 1), ylim=c(0, 1), 
        text(x=1, y=1, adj=0.5, cex=1.5, labels=gsub(x=phi, pattern="_", replacement=" = ", fixed=T))
      }
      plot(1, 1, axes=F, ann=F, type="n")#xlim=c(0, 1), ylim=c(0, 1), 
      text(x=1, y=1, adj=0.5, cex=1.5, labels="estimation method")
      par(mar=c(0.1, 0.1, 0.1, 0.1), mgp=c(2, 0.3, 0), tcl=-0.15, las=1)
      plot(1, 1, axes=F, ann=F, type="n")#xlim=c(0, 1), ylim=c(0, 1), 
      text(0.8, 1, labels="estimated standard deviation in parental investment", cex=1.5, srt=90)
      for(m in sample.m){
        plot(1, 1, axes=F, ann=F, type="n")#xlim=c(0, 1), ylim=c(0, 1), 
        text(1, 1, labels=m, cex=1.5, srt=90)
      }
  
  ###################################################################################################
    ##Fig. SI 5 to SI 12:
      ##general preperation:
        ##function for plotting:
        plot.fun.null.2<-function(plot.data, n.inf, n.obs, phi, x.bin.w, y.bin.w, scenario, base=10){
          ##extract relevant part wrt number parents and number observations per parent from list handed over to plot.data:
          this.summary=plot.data[[paste(c(n.inf, "inf", n.obs, "obs"), collapse="_")]]
          ##extract relevant part wrt phi:
          this.this.summary=this.summary[[paste(c("phi", phi), collapse="_")]]
          ##extract objects to be plotted (modelled and empricically estimated BLUPs and mean age per offspring):
          emp.where.rand=unlist(this.this.summary[["emp.est.1B"]])
          mod.where.rand=this.this.summary[["mod.est.1B"]]
          emp.where.real=unlist(this.this.summary[["emp.est.1C"]])
          mod.where.real=this.this.summary[["mod.est.1C"]]
          mean.age.rand=as.vector(this.this.summary[["mean.age.1B"]])
          mean.age.real=as.vector(this.this.summary[["mean.age.1C"]])
          xlim=range(c(mean.age.rand, mean.age.real))##set x-axis limits
          ##prep. for plotting (not evaluated):
          max.abs.diff=max(c(abs(mod.where.rand-median(mod.where.rand)), abs(emp.where.rand-median(emp.where.rand)),
                            abs(mod.where.real-median(mod.where.real)), abs(emp.where.real-median(emp.where.real))))
          mtext.cex=0.8##set size of margin text
          ##bin two-dimensional (i.e., create grid of average offsrping age and BLUPs):
          to.plot.emp.rand=plot.bin.fun(x=mean.age.rand, y=emp.where.rand, x.bin.w=x.bin.w, y.bin.w=y.bin.w)
          to.plot.mod.rand=plot.bin.fun(x=mean.age.rand, y=mod.where.rand, x.bin.w=x.bin.w, y.bin.w=y.bin.w)
          to.plot.emp.real=plot.bin.fun(x=mean.age.real, y=emp.where.real, x.bin.w=x.bin.w, y.bin.w=y.bin.w)
          to.plot.mod.real=plot.bin.fun(x=mean.age.real, y=mod.where.real, x.bin.w=x.bin.w, y.bin.w=y.bin.w)
          ylim=range(c(to.plot.emp.rand$y, to.plot.mod.rand$y, to.plot.emp.real$y, to.plot.mod.real$y))##set y-axis limits
          y.at=pretty(ylim)##... and where to put axis ticks
          y.at=y.at[y.at>=min(ylim) & y.at<=max(ylim)]##subset them to within range of y-values
          ##determine maximum number of data per cell of the grids:
          freq.max=max(c(to.plot.emp.rand$freq, to.plot.mod.rand$freq, to.plot.emp.real$freq, to.plot.mod.real$freq))
          ##log-transform number of data per cell of grid:
          to.plot.emp.rand$s.freq=log(to.plot.emp.rand$freq+1)/log(freq.max+1)
          to.plot.mod.rand$s.freq=log(to.plot.mod.rand$freq+1)/log(freq.max+1)
          to.plot.emp.real$s.freq=log(to.plot.emp.real$freq+1)/log(freq.max+1)
          to.plot.mod.real$s.freq=log(to.plot.mod.real$freq+1)/log(freq.max+1)
          ##set values to be shown in legend:
          legend.labels=c(base^(0:floor(log(max(freq.max), base=base))))
          ##determine distribution of conventional and modelled estimates:
          to.add.emp.rand=aggregate(x=to.plot.emp.rand$freq, to.plot.emp.rand[, "y", drop=F], sum)
          colnames(to.add.emp.rand)[2]="freq"
          to.add.mod.rand=aggregate(x=to.plot.mod.rand$freq, to.plot.mod.rand[, "y", drop=F], sum)
          colnames(to.add.mod.rand)[2]="freq"
          to.add.emp.real=aggregate(x=to.plot.emp.real$freq, to.plot.emp.real[, "y", drop=F], sum)
          colnames(to.add.emp.real)[2]="freq"
          to.add.mod.real=aggregate(x=to.plot.mod.real$freq, to.plot.mod.real[, "y", drop=F], sum)
          colnames(to.add.mod.real)[2]="freq"
          ##determine maximum frequency (needed for x-axis limits):
          f.dist.max=max(c(to.add.emp.rand$freq, to.add.mod.rand$freq, to.add.emp.real$freq, to.add.mod.real$freq))
          ##create layout matrix and set plot paramaters:
          lmat=matrix(1:8, ncol=4, byrow=T)
          lmat=cbind(13:14, 9:10, lmat[, 1:2], 11:12, lmat[, 3:4])
          lmat=rbind(c(NA, NA, rep(15, 2), NA, rep(16, 2)), lmat, c(NA, NA, 17:18, NA, 19:20))
          lmat=rbind(c(NA, NA, rep(21, 5)), lmat)
          lmat[is.na(lmat)]=(1:sum(is.na(lmat)))+max(lmat, na.rm=T)
          layout(lmat, widths=c(1, rep(c(1.25, rep(5, 2)), times=2)), heights=c(1, 1, 5, 5, 1))
          par(mar=c(0.2, 0.5, 0.5, 0.2), mgp=c(1.7, 0.3, 0), tcl=-0.15, las=0, cex.lab=1.25)
          ##plotting
          ##main plots, conventional estimates, random sampling:
          plot(x=1, y=1, xlab="offspring mean age", ylab="conventional parental investment", 
              ylim=ylim+c(-y.bin.w, y.bin.w)/2, xlim=xlim+c(-x.bin.w, x.bin.w)/2, pch=19, cex=0.3, type="n", bty="n", xaxs="i", yaxs="i", yaxt="n", xaxt="n")
          axis(side=1, labels=NA)
          rect(xleft=to.plot.emp.rand$x-x.bin.w/2, xright=to.plot.emp.rand$x+x.bin.w/2, ybottom=to.plot.emp.rand$y-y.bin.w/2, ytop=to.plot.emp.rand$y+y.bin.w/2, 
              border=NA, col=grey(1-to.plot.emp.rand$s.freq))
          axis(side=2, at=y.at)
          box()
          mtext(text="(a)", side=3, line=1-2, cex=mtext.cex)
          ##add legend:
            xx=par()[["usr"]]
            y.vals=seq(xx[3]+diff(range(xx[3:4]))*l.edge.dist, xx[3]+diff(range(xx[3:4]))*l.y.extent, length.out=l.resol)
            segments(x0=xx[1]+diff(range(xx[1:2]))*l.edge.dist, x1=xx[1]+diff(range(xx[1:2]))*l.x.extent, 
                    y0=y.vals, y1=y.vals, col=grey(seq(1, 0, length.out=l.resol)), lend=1)
            labels=legend.labels
            y=log(labels+1)
            y=y/log(freq.max+1)
            y=y*diff(c(xx[3]+diff(range(c(xx[3:4])))*l.edge.dist, xx[3]+diff(range(xx[3:4]))*l.y.extent)) + (xx[3]+diff(range(xx[3:4]))*l.edge.dist)
            text(x=rep(xx[1]+diff(range(xx[1:2]))*l.x.extent + max(strwidth(s=labels, cex=l.text.cex)) + diff(par()$usr[c(1, 2)])*0.01, length(y)), 
                y=y, labels=labels, adj=c(1, 0.5), cex=l.text.cex)
            rect(xleft=xx[1]+diff(range(xx[1:2]))*l.edge.dist, xright=xx[1]+diff(range(xx[1:2]))*l.x.extent,
                ybottom=xx[3]+diff(range(xx[3:4]))*l.edge.dist, ytop=xx[3]+diff(range(xx[3:4]))*l.y.extent, lwd=0.5)
          ##distribution of conventional estimates:
          plot(1, 1, type="n", xlim=c(0, f.dist.max), xlab="frequency", ylab="", ylim=ylim+c(-y.bin.w, y.bin.w)/2, yaxs="i", yaxt="n", xaxt="n")
          axis(side=1, labels=NA)
          axis(side=2, at=y.at, labels=NA)
          rect(xleft=0, xright=to.add.emp.rand$freq, ybottom=to.add.emp.rand$y-x.bin.w/2, ytop=to.add.emp.rand$y+x.bin.w/2, col=grey(level=0.5), border=NA)
          mtext(text="(b)", side=3, line=1-2, cex=mtext.cex)
          ##modeled estimates, random sampling:
          plot(x=1, y=1, xlab="offspring mean age", ylab="conventional parental investment", 
              ylim=ylim+c(-y.bin.w, y.bin.w)/2, xlim=xlim+c(-x.bin.w, x.bin.w)/2, pch=19, cex=0.3, type="n", bty="n", xaxs="i", yaxs="i", yaxt="n", xaxt="n")
          axis(side=1, labels=NA)
          rect(xleft=to.plot.mod.rand$x-x.bin.w/2, xright=to.plot.mod.rand$x+x.bin.w/2, ybottom=to.plot.mod.rand$y-y.bin.w/2, ytop=to.plot.mod.rand$y+y.bin.w/2, 
              border=NA, col=grey(1-to.plot.mod.rand$s.freq))
          axis(side=2, at=y.at)
          box()
          mtext(text="(c)", side=3, line=1-2, cex=mtext.cex)
          ##distribution of modeled estimates:
          plot(1, 1, type="n", xlim=c(0, f.dist.max), xlab="frequency", ylab="", ylim=ylim+c(-y.bin.w, y.bin.w)/2, yaxs="i", yaxt="n", xaxt="n")
          axis(side=1, labels=NA)
          axis(side=2, at=y.at, labels=NA)
          rect(xleft=0, xright=to.add.mod.rand$freq, ybottom=to.add.mod.rand$y-x.bin.w/2, ytop=to.add.mod.rand$y+x.bin.w/2, col=grey(level=0.5), border=NA)
          mtext(text="(d)", side=3, line=1-2, cex=mtext.cex)
          
          ##conventional estimates, constrained sampling:
          plot(x=1, y=1, xlab="offspring mean age", ylab="conventional parental investment", 
              ylim=ylim+c(-y.bin.w, y.bin.w)/2, xlim=xlim+c(-x.bin.w, x.bin.w)/2, pch=19, cex=0.3, type="n", bty="n", xaxs="i", yaxs="i", yaxt="n")
          rect(xleft=to.plot.emp.real$x-x.bin.w/2, xright=to.plot.emp.real$x+x.bin.w/2, ybottom=to.plot.emp.real$y-y.bin.w/2, ytop=to.plot.emp.real$y+y.bin.w/2, 
              border=NA, col=grey(1-to.plot.emp.real$s.freq))
          axis(side=2, at=y.at)
          box()
          mtext(text="(e)", side=3, line=1-2, cex=mtext.cex)
          ##distribution of conventional estimates:
          plot(1, 1, type="n", xlim=c(0, f.dist.max), xlab="frequency", ylab="", ylim=ylim+c(-y.bin.w, y.bin.w)/2, yaxs="i", yaxt="n")
          axis(side=2, at=y.at, labels=NA)
          rect(xleft=0, xright=to.add.emp.real$freq, ybottom=to.add.emp.real$y-x.bin.w/2, ytop=to.add.emp.real$y+x.bin.w/2, col=grey(level=0.5), border=NA)
          mtext(text="(f)", side=3, line=1-2, cex=mtext.cex)
          ##modeled estimates, constrained sampling:
          plot(x=1, y=1, xlab="offspring mean age", ylab="conventional parental investment", 
              ylim=ylim+c(-y.bin.w, y.bin.w)/2, xlim=xlim+c(-x.bin.w, x.bin.w)/2, pch=19, cex=0.3, type="n", bty="n", xaxs="i", yaxs="i", yaxt="n")
          rect(xleft=to.plot.mod.real$x-x.bin.w/2, xright=to.plot.mod.real$x+x.bin.w/2, ybottom=to.plot.mod.real$y-y.bin.w/2, ytop=to.plot.mod.real$y+y.bin.w/2, 
              border=NA, col=grey(1-to.plot.mod.real$s.freq))
          axis(side=2, at=y.at)
          box()
          mtext(text="(g)", side=3, line=1-2, cex=mtext.cex)
          ##distribution of modeled estimates:
          plot(1, 1, type="n", xlim=c(0, f.dist.max), xlab="frequency", ylab="", ylim=ylim+c(-y.bin.w, y.bin.w)/2, yaxs="i", yaxt="n")
          axis(side=2, at=y.at, labels=NA)
          rect(xleft=0, xright=to.add.mod.real$freq, ybottom=to.add.mod.real$y-x.bin.w/2, ytop=to.add.mod.real$y+x.bin.w/2, col=grey(level=0.5), border=NA)
          mtext(text="(h)", side=3, line=1-2, cex=mtext.cex)
          
          ##headers, axes labels, etc.
          par(mar=c(0.2, 0.2, 0.5, 0.2), mgp=c(1.7, 0.3, 0), tcl=-0.15, las=1, cex.lab=1.25)
          for(lab in c("parental investment", "parental investment", "", "")){
            plot(1, 1, axes=F, ann=F, type="n")
            if(nchar(lab)>0){text(x=0.9, 1, labels=lab, cex=axis.label.cex, srt=90)}
          }
          for(lab in c("random sampling", "constrained sampling")){
            plot(1, 1, axes=F, ann=F, type="n")
            text(1, 1, labels=lab, cex=main.cex, srt=90)
          }
          par(mar=c(0.2, 0.5, 0.2, 0.2), mgp=c(1.7, 0.3, 0), tcl=-0.15, las=1, cex.lab=1.25)
          for(lab in c("conventional method", "modeling method")){
            plot(1, 1, axes=F, ann=F, type="n")
            text(1, 1, labels=lab, cex=main.cex)
          }
          for(i in 1:2){
            for(lab in c("offspring mean age", "frequency")){
              plot(1, 1, axes=F, ann=F, type="n")
              text(1, 1, labels=lab, cex=axis.label.cex)
            }
          }
          plot(1, 1, type="n", ann=F, axes=F)
          t1=paste(c(n.inf, " offspring, ", n.obs, " observations per offspring"), collapse="")
          t2=paste(c("phi = ", phi, ", ", scenario), collapse="")
          text(x=1, y=1.2, labels=bquote(bold(.(t1))), cex=main.cex)
          text(x=1, y=0.8, labels=bquote(bold(.(t2))), cex=main.cex)
        }
      ##graphical parameters etc.:
        l.edge.dist=0.04
        l.x.extent=0.1
        l.y.extent=0.3
        l.resol=250
        l.text.right=0.215
        l.text.cex=0.6
        axis.label.cex=1.2
        main.cex=1.5
        x.bin.w=0.01
        y.bin.w=0.01
    ##and the plot:
      n.obs=21##number observations per parent (can be 21 or 81)
      n.inf=20##number parents (can be 20 or 80)
      phi=5##precision parameter (can be 5, 10, 20, or 40)
      scenario="null-scenario"
      dev.off()
      X11(height=6, width=6)
      plot.fun.null.2(plot.data=all.summary, n.inf=n.inf, n.obs=n.obs, phi=phi, x.bin.w=x.bin.w, y.bin.w=y.bin.w, 
                  scenario=scenario)

  ##########################################################################################################################
  ##Fig. SI 13 to SI 16:
    ##general settings:
    ##bin widths in x- and y-direction:
    x.bin.w=0.05
    y.bin.w=0.05
    ##define function creating one plot:
    plot.fun.bal<-function(phi, sampling, sim, method, x.bin.w, y.bin.w, base=10){
      ##extract results to be plotted (relevant part wrt phi):
      this.all.summary.diff=all.sim.res.bal.with.diff[[paste(c("phi", phi), collapse="_")]]
      ##extract values to be plotted (simulated and estimated parent specific investment):
      simulated=lapply(this.all.summary.diff, "[[", paste(c("sim.blups.2", sim), collapse=""))
      model.ests=lapply(this.all.summary.diff, "[[", paste(c("mod.est.2", sim), collapse=""))
      emp.ests=lapply(this.all.summary.diff, "[[", paste(c("emp.est.2", sim), collapse=""))
      ##determine x- and ylims:
      all.xlim=range(unlist(simulated))
      all.ylim=range(c(unlist(model.ests), unlist(emp.ests)))
      ##determine estimates to be plotted and simulated parent specfic investments:
      to.get=ifelse(method=="conventional", paste(c("emp.est.2", sim), collapse=""), paste(c("mod.est.2", sim), collapse=""))
      to.plot=lapply(this.all.summary.diff, function(x){##i.e., for each combination of numer parents and number samples per parent
        ##bin simulated and estimated parent specific investment:
        plot.bin.fun(x=unlist(lapply(this.all.summary.diff, "[[", paste(c("sim.blups.2", sim), collapse=""))), 
            y=unlist(lapply(this.all.summary.diff, "[[", to.get)), 
            x.bin.w=x.bin.w, y.bin.w=y.bin.w)
      })
      ##determine maximum value in any of the grid cells:
      max.freq=max(unlist(lapply(to.plot, "[[", "freq")))
      ##set up layout matrix:
      lmat=rbind(
        rep(c(14, 11), each=2),
        c(12, 12, 2, 4),
        c(1, 10, 3, 5),
        c(6, 10, 7, 8),
        c(13, 13, 9, 9)
      )
      layout(lmat, widths=rep(c(1, 5), each=2), heights=c(1, 1, rep(5, 2), 1.5))
      ##set graphical parameters:
      par(mar=c(0.2, 0.2, 0.2, 0.2), mgp=c(1.7, 0.3, 0), tcl=-0.15, las=1)
      for(n.inf in c(20, 80)){##for number parents being 20 or 80
        ##add number parents to plot:
        plot(1, 1, type="n", axes=F, ann=F)
        text(1, 1, labels=paste(c(n.inf, "inf."), collapse = " "), cex=1.5, srt=90)
        for(n.obs in c(21, 81)){
          if(n.inf==20){##add number obs. per parents to plot:
            plot(1, 1, type="n", axes=F, ann=F)
            text(1, 1, labels=paste(c(n.obs, "obs."), collapse = " "), cex=1.5)
          }
          ##extract part of list to.plot to ne plotted (relevant number parents and observations per parent):
          x.plot=to.plot[[paste(c(n.inf, "inf", n.obs, "obs"), collapse="_")]]
          ##... and plot it:
          plot(1, 1, xlim=all.xlim, ylim=all.ylim, type="n", ann=F, xaxt="n", yaxt="n")
          rect(xleft=x.plot$x-x.bin.w/2, xright=x.plot$x+x.bin.w/2, ybottom=x.plot$y-y.bin.w/2, x.plot$y+y.bin.w/2,
              col=grey(level=1-log(1+x.plot$freq)/log(1+max.freq)), border=NA)
          ##add axes:
          if(n.obs==21){
            axis(side=2)
          }else{
            axis(side=2, labels=NA)
          }
          if(n.inf==80){
            axis(side=1)
          }else{
            axis(side=1, labels=NA)
          }
        }
      }
      ##add legend:
      l.edge.dist=0.04
      l.x.extent=0.1
      l.y.extent=0.5
      l.resol=250
      l.text.right=0.15
      l.text.cex=0.6
      xx=par()[["usr"]]
      y.vals=seq(xx[4]-diff(range(xx[3:4]))*l.y.extent, xx[4]-diff(range(xx[3:4]))*l.edge.dist, length.out=l.resol)
      segments(x0=xx[1]+diff(range(xx[1:2]))*l.edge.dist, x1=xx[1]+diff(range(xx[1:2]))*l.x.extent, 
              y0=y.vals, y1=y.vals, col=grey(seq(1, 0, length.out=l.resol)), lend=1)
      labels=c(base^(0:floor(log(max.freq, base=base))))
      y=log(labels+1, base=base)
      y=y/log(1+max.freq, base=base)
      y=y*diff(range(y.vals))+min(y.vals)
      text(x=rep(xx[1]+diff(range(xx[1:2]))*l.x.extent+max(strwidth(s=labels, cex=l.text.cex))+diff(par()$usr[c(1, 2)])*0.01, length(y)), y=y, labels=labels, adj=c(1, 0.5), cex=l.text.cex)
      rect(xleft=xx[1]+diff(range(xx[1:2]))*l.edge.dist, xright=xx[1]+diff(range(xx[1:2]))*l.x.extent,
          ybottom=xx[4]-diff(range(xx[3:4]))*l.y.extent, ytop=xx[4]-diff(range(xx[3:4]))*l.edge.dist, col=NA, lwd=0.5)
      plot(1, 1, type="n", axes=F, ann=F)
      text(1, y=0.8, labels="simulated parental investment", cex=1.5)
      plot(1, 1, type="n", axes=F, ann=F)
      text(x=0.8, 1, labels=paste(c(method, " parental investment"), collapse=""), cex=1.5, srt=90)
      plot(1, 1, type="n", axes=F, ann=F)
      t2add=paste(c("phi = ", phi, ", ", sampling ," sampling, ", method, " investemt"), collapse="")
      text(x=1, labels=bquote(bold(.(t2add))), cex=1.5, adj=0.5)
    }

  ##plot:
    phi=5##can be one of 5, 10, 20, or 40
    sampling="balanced"##don't change (we created these plots only for balanced sampling)
    sim="A"##don't change (we created these plots only for balanced sampling)
    method="conventional"##can be "conventional" or "modeled"
    dev.off()
    X11(height=3.5, width=6)
    plot.fun.bal(phi=phi, sampling=sampling, sim=sim, method=method, x.bin.w=x.bin.w, y.bin.w=y.bin.w)
  

  #########################################################################################################################
  ##Fig. SI 17 to SI 24:
      ##define function that bins in two dimensions and determines average age per bin:
      plot.bin.fun.age<-function(x, y, x.bin.w, y.bin.w, age){
        breaks=seq(0, max(abs(x))+x.bin.w, by=x.bin.w)
        binned.x=cut(abs(x), breaks, labels=F, include.lowest=T)
        binned.x=(x.bin.w/2 + (binned.x-1)*x.bin.w)*sign(x)
        breaks=seq(0, max(abs(y))+y.bin.w, by=y.bin.w)
        binned.y=cut(abs(y), breaks=breaks, labels=F, include.lowest=T)
        binned.y=(y.bin.w/2 + (binned.y-1)*y.bin.w)*sign(y)
        to.plot=aggregate(1:length(x), by=data.frame(x=binned.x, y=binned.y), length)
        names(to.plot)[ncol(to.plot)]="freq"
        to.plot$mean.age=aggregate(age, by=data.frame(xx=binned.x, y=binned.y), mean)$x
        return(to.plot)
      }
      ##basic settings:
      to.do=apply(expand.grid(c(20, 80), c(21, 81)), 1, function(x){
        paste(c(x[1], "inf", x[2], "obs"), collapse="_")
      })
      ##create layout matrix:
      lmat=rbind(
        rep(c(14, 11), each=2),
        c(12, 12, 2, 4),
        c(1, 10, 3, 5),
        c(6, 10, 7, 8),
        c(13, 13, 9, 9)
      )
      ##legend parameters;
      l.x.edge.dist=0.05
      l.y.edge.dist=0.05
      l.x.extent=0.3
      l.y.extent=0.35
      l.resol=20
      l.text.right=0.215
      l.text.cex=0.6
      ##other parameters:
      x.bin.w=0.05
      y.bin.w=0.05
      freq.spread=0.5
      ##define function for plotting:
      plot.fun.parental.var<-function(what, extract, phi, x.bin.w, y.bin.w, freq.spread, adj.legend=T){
        ##determine what to add to the title of the plot:
        sampling=ifelse(extract=="B", "random", "constrained")
        estimation=ifelse(what=="emp", "conventional", "modeled")
        ##determine frequency distribution of binned outcomes and average age per combination of bins
        all.to.plot=lapply(all.summary.2BC, function(x){
          ##extract relevant part wrt phi from all.summary.2BC:
          xx=x[[paste(c("phi", phi), collapse="_")]]
          return(plot.bin.fun.age(##bin simulated and estimated parental investment and average age per bin
            x=xx[[paste(c("sim.where.2", extract), collapse="")]], 
            y=xx[[paste(c(what, ".where.2", extract), collapse="")]], 
            x.bin.w=x.bin.w, y.bin.w=y.bin.w, 
            age=xx[[paste(c("mean.age.2", extract), collapse="")]]))
        })
        ##add color value to all.to.plot:
          ##determine overall age range:
          age.range=range(unlist(lapply(all.to.plot, function(x){
            range(x$mean.age)
          })))
          ##determine overall range of number data points per cell of grid:
          freq.range=range(unlist(lapply(all.to.plot, function(x){
            range(x$freq)
          })))
          ##determine color per grid cell:
          all.to.plot=lapply(all.to.plot, function(x){
            xx.freq=(log(x$freq)-log(min(freq.range)))/log(max(freq.range))
            xx.freq=(1-freq.spread)+xx.freq*freq.spread
            xx.age=(x$mean.age-min(age.range))/diff(range(age.range))
            x$col=unlist(lapply(1:nrow(x), function(y){
              plasma(n=1, begin=1-xx.age[y], end=1-xx.age[y], alpha=xx.freq[y])
            }))
            return(x)
          })
        ##determine x- and y-limit:
        all.xlim=range(unlist(lapply(all.to.plot, function(x){
          range(x$x)
        })))
        all.ylim=range(unlist(lapply(all.to.plot, function(x){
          range(x$y)
        })))
        ##plot:
        layout(lmat, widths=rep(c(1, 5), each=2), heights=c(1, 1, rep(5, 2), 1.5))
        par(mar=c(0.2, 0.2, 0.2, 0.2), mgp=c(1.7, 0.3, 0), tcl=-0.15, las=1)
        for(n.inf in c(20, 80)){##for 20 or 80 parents
          plot(1, 1, type="n", axes=F, ann=F)##add number parents
          text(1, 1, labels=paste(c(n.inf, "inf."), collapse = " "), cex=1.5, srt=90)
          for(n.obs in c(21, 81)){##for 21 or 81 observations per parent
            if(n.inf==20){##add number observations per parent
              plot(1, 1, type="n", axes=F, ann=F)
              text(1, 1, labels=paste(c(n.obs, "obs."), collapse = " "), cex=1.5)
            }
            ##extract relevant data wrt number parents and number observations per parent:
            this.to.plot=all.to.plot[[paste(c(n.inf, "inf", n.obs, "obs"), collapse="_")]]
            ##begin with plot:
            plot(1, 1, type="n", xlab="", ylab="", xaxt="n", yaxt="n", xlim=all.xlim, ylim=all.ylim)
            ##add axes:
            if(n.obs==21){
              axis(side=2)
            }else{
              axis(side=2, labels=NA)
            }
            if(n.inf==80){
              axis(side=1)
            }else{
              axis(side=1, labels=NA)
            }
            rect(xleft=this.to.plot$x-x.bin.w/2, xright=this.to.plot$x+x.bin.w/2, 
                ybottom=this.to.plot$y-y.bin.w/2, ytop=this.to.plot$y+y.bin.w/2, border=NA, col=this.to.plot$col)
          }
        }
        ##legend:
          xbw=diff(c(par()$usr[1]+l.x.edge.dist*diff(par()$usr[c(1, 2)]), par()$usr[1]+l.x.extent*diff(par()$usr[c(1, 2)])))/(l.resol+1)
          ybw=abs(diff(c(par()$usr[3]-l.x.edge.dist*diff(par()$usr[c(1, 2)]), par()$usr[3]-l.x.extent*diff(par()$usr[c(1, 2)])))/(l.resol+1))
          to.add=data.frame(expand.grid(freq=seq(0, max(log(freq.range)), length.out=l.resol), age=seq(min(age.range), max(age.range), length.out=l.resol)))
          xx.freq=(to.add$freq-log(min(freq.range)))/log(max(freq.range))
          xx.freq=(1-freq.spread)+xx.freq*freq.spread
          xx.age=(to.add$age-min(age.range))/diff(range(age.range))
          #if(any(xx.freq>1) | any(xx.age>1)){browser()}
          to.add$col=unlist(lapply(1:nrow(to.add), function(y){
            plasma(n=1, begin=1-xx.age[y], end=1-xx.age[y], alpha=xx.freq[y])
          }))
          to.add=data.frame(to.add, xleft=match(to.add$freq, sort(unique(to.add$freq))), ybottom=match(to.add$age, sort(unique(to.add$age))))
          to.add$xleft=seq(par()$usr[1]+l.x.edge.dist*diff(par()$usr[c(1, 2)]), par()$usr[1]+l.x.extent*diff(par()$usr[c(1, 2)]), length.out=l.resol+1)[to.add$xleft]
          to.add$ybottom=seq(par()$usr[4]-l.y.extent*diff(par()$usr[c(3, 4)]), par()$usr[4]-l.y.edge.dist*diff(par()$usr[c(3, 4)]), length.out=l.resol+1)[to.add$ybottom]
          rect(xleft=to.add$xleft, xright=to.add$xleft+xbw, ybottom=to.add$ybottom, ytop=to.add$ybottom+ybw, border=NA, col=to.add$col)
          legend.text.x=mean(to.add$xleft)+xbw/2
          ##add legend text, age values:
          to.add=pretty(age.range)
          to.add=to.add[to.add>=min(age.range) & to.add<=max(age.range)]
          if(adj.legend){
            if(length(to.add)>=5 & length(to.add)%%2==1){
              to.add=to.add[seq(1, length(to.add), by=2)]
            }
            if(length(to.add)>=6 & length(to.add)%%2==0){
              to.add=to.add[seq(1, length(to.add), by=2)]
            }
          }
          xx=diff(c(par()$usr[4]-l.y.extent*diff(par()$usr[c(3, 4)]), par()$usr[4]-l.y.edge.dist*diff(par()$usr[c(3, 4)])))
          where.y=(to.add-min(age.range))/diff(range(age.range))
          where.y=par()$usr[4]-l.y.extent*diff(par()$usr[c(3, 4)]) + where.y*xx
          where.x=par()$usr[1]+l.x.extent*diff(par()$usr[c(1, 2)])+max(strwidth(s=to.add, cex=1))
          to.add=sprintf("%.1f", to.add)
          text(labels=to.add, x=where.x+diff(par()$usr[c(1, 2)])*0.01, y=where.y, adj=c(1, 0.5))
          text(x=where.x+diff(par()$usr[c(1, 2)])*0.04, y=mean(where.y), labels="age", srt=90)
          ##add legend text, N:
          to.add=10^(0:floor(max(log(freq.range, base=10))))
          where.x=(log(to.add)-min(log(freq.range)))/diff(range(log(freq.range)))
          xx=diff(c(par()$usr[1]+l.x.edge.dist*diff(par()$usr[c(1, 2)]), par()$usr[1]+l.x.extent*diff(par()$usr[c(1, 2)])))
          where.x=par()$usr[1]+l.x.edge.dist*diff(par()$usr[c(1, 2)]) + where.x*xx
          where.y=par()$usr[4]-l.y.extent*diff(par()$usr[c(3, 4)])#-max(strwidth(s=to.add, cex=1))
          text(labels=to.add, x=where.x, y=where.y, pos=1)#, srt=90, adj=c(0.5, 1))#, -diff(par()$usr[c(3, 4)])*0.01
          text(x=legend.text.x, y=where.y-diff(par()$usr[c(3, 4)])*0.15, labels="N estimates")
        ##axes labels, header, etc.:
        plot(1, 1, type="n", axes=F, ann=F)
        text(1, y=0.8, labels="simulated parental investment", cex=1.5)
        plot(1, 1, type="n", axes=F, ann=F)
        text(x=0.8, 1, labels=paste(c(estimation, " parental investment"), collapse=""), cex=1.5, srt=90)
        plot(1, 1, type="n", axes=F, ann=F)
        t2add=paste(c("phi = ", phi, ", ", sampling, " sampling, ", estimation, " investemt"), collapse="")
        text(x=1, labels=bquote(bold(.(t2add))), cex=1.5, adj=0.5)
      }

    ##plotting:
    what="emp"##can be one of "emp" (i.e., conventional) or "mod" (i.e., modelled)
    extract="B"##can be one of "B" (i.e., random sampling), or "C" (i.e., constrained sammpling)
    phi=5##can be one of 5, 10, 20, or 40
    dev.off()
    X11(height=3.5*1.25, width=6*1.25)
    plot.fun.parental.var(what=what, extract=extract, phi=phi, x.bin.w=x.bin.w, y.bin.w=y.bin.w, freq.spread=freq.spread)
```
