## supporting information for "Overcoming Pitfalls when Estimating Variation in Parental Care: A Simulation Study"

### Supporting Information for ”Overcoming Pitfalls when Estimating Variation in Parental behavior: A Simulation Study”

January 2, 2024

In the following, we present the full results of almost all our simulations. The document is divided into three sections. In the first, we present the results we found when simulating no differences in parental behavior (null-scenario). In the second section, we present the results we found when simulating variation in parental behavior (variable parental behavior-scenario). The code for the simulations can be found at .... In the final section, we expand the Discussion section of the main text.

#### 1 No Variation in Parental Behavior (Null-Scenario)

##### 1.1 Evaluation Methods

###### 1.1.1 Estimated Standard Deviations of Parent-Specific Behavior

We first evaluated simulations in which parents did not vary in parental behavior. Here we first focused on the magnitude of the estimated variation between the Behavior of different parents.

In case of the conventional method, we first determined the average behavior per parent and then determined the standard deviation of these averages for each simulated data set. Since we did not simulate any differences between parents, these ’conventional’ standard deviations should be ideally mostly zero or close to zero. We also applied the modelling approach where we estimated the variation in parental behavior using a Generalized Linear Mixed Model (GLMM; Baayen, 2008) with beta error structure and logit link function (McCullagh and Nelder, 1989; Bolker, 2008). The models comprised a fixed effect for the age of the offspring, a random intercepts effect for the ID of the parent-offspring dyad, and a random slope of offspring age. Offspring age entered the model after subtracting one, the mid point between birth and two years of age, the period for which we simulated data. This has the effect that in case the random slope of offspring age in parent ID is not estimated to be zero, the estimated parent specific behaviors (Best Linear Unbiased Predictors, BLUPs; Baayen, 2008) refer to an offspring aged one year. For each simulated data set we fitted such a model and extracted from it the ’modeled’ standard deviation (in link space) for the variation in parental behavior. Again, these standard deviations should ideally mostly be zero or close to zero, given that we did not simulate any variation between parents with regard to their behavior.

One issue with the comparison of the conventionally estimated and the modeled standard deviations was that the modeled standard deviations were determined in link space. Hence, when determining the conventional standard deviations of the average response per parent, the two standard deviations would not be comparable. In order to make them comparable, we first transformed the average response per parent (i.e., the conventional 'estimate' of parent specific behavior) to link space (i.e., $mean'_{B_j} = log(\frac{mean_{B_j}}{1-mean_{B_j}})$ where $mean_{B_j}$ is the average behavior per parent $j$) and then determined the conventionally estimated standard deviation using the values of $mean'_{B_j}$.

We show one plot for each of the four combinations of the two sample sizes with regard to the number of offspring (20 or 80) and the number of observations per offspring (21 or 81). Each of the four plots is then subdivided into three rows, one for each of balanced, random, and constrained sampling (see Fig. 3 of the main text) and four columns, one for each of the four values of the precision parameter *phi* we simulated (see Fig. 4 of the main text). We binned the estimated standard deviations in the resulting plots (Fig. SI 1 to SI 4). Figure SI 1 to SI 4 are summarized in Fig. 6 of the main text.

##### 1.1.2 Estimated Parent-Specific behavior and Average Offspring Age

For simulations with the random and the constrained sampling scheme, we further evaluated the extent to which the estimated parent-specific behavior was related to the average age of the offspring. In the simulated data, the parental behavior decreased as the offspring got older (Fig. 2 in the main text), and in case of random or constrained sampling, the average age (across the observation days) will differ between offspring. Hence, it seemed likely that in such cases the conventional method will overestimate parental behavior for relatively young offspring and underestimate it for relatively old offspring.

To assess the extent to which this was indeed the case and whether the modeling approach suffered in a similar way, we plotted the conventional and the modeled estimates of parent-specific behavior against the average age of the offspring. To this end, we determined, for each simulated data set, the average age of each offspring at the days at which it was observed and also the conventional and the modeled estimates of the parent-specific behaviors. Again, we evaluated the conventional estimates of parental effort in link space (see above). We extracted the conventional and modeled parent-specific estimates as well as the average offspring age separately for each combination of number parent-offspring dyads, number of observations per parent-offspring dyad, sampling scheme (random or constrained sampling; see Fig. 3 of the main text), and precision parameter (*phi*) and pooled them across all simulations with a given setting. We then simply plotted the conventional and the modeled estimates of parent-specific behavior against the average age of the respective offspring.

We created one Figure for each of the combinations of number of parent-offspring dyads, number observations per parent-offspring dyad, and precision parameter *phi*. As the results were not strongly affected by the precision parameter, we only show them for precision parameters of 5 and 40 which were the smallest and largest, respectively. In the resulting plots (Fig. SI 5 to SI 12), we binned offspring age and the estimated parent-specific behavior. This binning solely served in keeping the size of this document manageable. Figure SI 5 to SI 12 are summarized in Fig. 7 of the main text (including also precision parameters being 10 or 20). For summarizing them we determined the $R^2$ of a regression of the Estimated parent-specific behavior against average offspring age, separately per combination parameter settings and estimation method (see also main text).

#### 1.2 Evaluation Results

##### 1.2.1 Estimated Standard Deviations of Parent-Specific behavior

Across all simulated values of *phi* and also for all sample sizes, we found that modeling parental behavior was superior to using the average behavior as an estimate of parental behavior. More specifically, the average variability (per simulation standard deviation of parental behavior) was invariably higher when using the conventional method compared to the modeling approach (Fig. SI 1 to SI 4). Furthermore, the conventionally estimated variation in parental behavior increased considerably from balanced over random to constrained sampling, while the results of the modeling approach were hardly affected by the sampling scheme. Compared to the effects of the sampling scheme and the estimation method (conventional or modeling), the effects of sample size were relatively minor. However, with large numbers of observations per offspring, the conventional approach in combination with balanced sampling and little stochastic variation in the simulated behavior (large value of *phi*) performed almost like the modeling approach (top right plots in Fig. SI 2 and SI 4). Overall, the simulated stochastic variation in the observations (precision parameter *phi*) had a minor effect on the results. Nevertheless, in most scenarios with regard to the number of offspring and observations per offspring, the estimated variation in parental effort decreased with increasing value of *phi* (i.e., smaller stochastic variation in the observations). This decrease happened when using the conventional as well as the modeling approach, but in case of the modeling approach the effect of stochastic variation in the behavior was much smaller than when using the conventional approach (Fig. SI 1 to SI 4).

##### 1.2.2 Estimated Parent-Specific Behavior and Average Offspring Age

Regarding a potential correlation between the average age at which offspring were observed and the conventionally estimated parent-specific behavior, we found that these were clearly negatively correlated, irrespective of sample size, sampling scheme, and precision parameter *phi*. In other words, the conventional estimate of parent-specific behavior overestimated parent-specific behavior in case of relatively young offspring and underestimated it in case of relatively old offspring (Fig. SI 5 to SI 12). The modeled parent-specific behavior, in turn, was not correlated with average offspring age.

A side aspect of this evaluation was that this time, we evaluated the individual (conventional and modeled) estimates of parent-specific behavior rather than their standard deviation. We again found that the magnitude of variation among parents was much larger when using the conventional than when using the modeling approach (compare (b) with (d) and (f) with (h) in Fig. SI 5 to SI 12).

### 2 Variation Parental Behavior (Variable Parental Behavior Scenario)

#### 2.1 Evaluation Methods

In case of simulated data comprising variation in parental behavior, we focused on how well the conventionally estimated and modeled estimates of parent-specific behavior retrieved the simulated ones. To this end, we extracted the average parental behavior (conventional approach) and also the estimated BLUPs (modeling approach) for each parent in each simulation. Again, for comparability we transformed the conventional estimates of parent-specific behavior into link space (see above). We then plotted the estimated behavior against the simulated behavior. We did so only for simulations with the smallest and largest precision parameter ($phi = 5$ or 40). As it seemed plausible that the conventional and perhaps also the modeled parent-specific behavior might in part depend on the average age of the offspring when using random or constrained sampling, we also investigated this question. To this end, we simply colored points on the plots according to the average age of the offspring.

In the resulting plots (Fig. SI 13 to Fig. SI 24), we binned the both simulated and estimated parental effort. The coloration of points depicts the number of estimates falling in the respective combination of binned variables (simulated and estimated parental effort). In Fig. SI 17 to Fig. SI 24 points and legends also depict the average age of the offspring per combination of binned variables. Figure SI 13 to Fig. SI 16 are summarized in Fig. 8 of the main text. To this end we fitted a linear regression of the estimated against the simulated parent-specific behavior to the data depicted in each sub-plot and extracted the $R^2$.

#### 2.2 Evaluation Results

Overall, the conventional as well as the modeled estimates of parent-specific behavior clearly correlated with the simulated ones (Fig. SI 13 to SI 24). In case of balanced sampling, both conventionally estimated and modeled parent-specific behavior retrieved the simulated ones fairly well (Fig. SI 13 to SI 16). Moreover, when we simulated balanced sampling in combination with small stochastic variation in the observed behavior (i.e., larger precision parameter), the agreement between the simulated and the estimated parent-specific behavior improved, and this was the case when using conventional estimates (compare Fig. SI 15 with Fig. SI 13) as well as modeled estimates (compare Fig. SI 16 with Fig. SI 14).

For random and constrained sampling, we also found both conventionally estimated and modeled parent-specific behavior to be correlated with the simulated ones (Fig. SI 17 to SI 24). However, conventionally estimated parent-specific behaviors were also clearly correlated with average offspring age (Fig. SI 17, SI 19, SI 21, and SI 23). In case of random sampling, the impact of average offspring age on the conventionally estimated parent-specific behavior was stronger with 21 observations per parent-offspring dyad as compared to 81 (compare left and right panel in Fig. SI 17 and SI 19). Furthermore, the effect of average offspring age on the conventionally estimated parent-specific behavior was particularly strong in case of constrained sampling (Fig. SI 21 and SI 23).

Comparing conventionally estimated and modeled parent-specific behavior for random and constrained sampling revealed that the modeled behavior much better captured the simulated ones (Fig. SI 18, SI 20, SI 22, and SI 24). Also, modeled parent-specific behaviors did not correlate obviously with average offspring age. The only exception from this we found when using constrained sampling in combination with relatively large stochastic variation in the observed behavior (precision parameter $phi = 5$): here the modeled parent-specific behavior tended to be slightly overestimated for older offspring and slightly underestimated for younger offspring (Fig. SI 22). However, the bias was much weaker than the one we found when using the conventional method.

### 3 Discussion

#### 3.1 Parental Behavior and Random Slopes

In the discussion we raised the question of how to estimate parent-specific behavior in the presence of a random slope of offspring age in parent and/or offspring ID. The problem arises from the fact that in such a case, the parent and/or offspring specific trajectories of the parental behavior as the offspring ages might cross over the period considered (see Fig. 11 of the main text). Here we propose a tentative and *ad-hoc* solution for how to proceed in such a case. The idea is to relate the parent specific trajectory to the average trajectory. More specifically, we propose to determine the integral between the parent-specific and the average trajectory (Fig. SI 25). For a parent with a generally relatively high behavior, this integral will be positive (Fig. SI 25a), for a parent with a generally relatively low behavior it will be negative (Fig. SI 25b), and for a parent with an overall average behavior it will be near zero (Fig. SI 25c). Practically, the probably simplest way to determine the parent-specific integral is to first determine the parent-specific and average behavior for each day between the minimum and maximum age of all offspring and then average the difference between the former and the latter per parent. These parent-specific estimates could then serve as estimates of overall relative parental behavior.

#### 3.2 Non-Linear Mixed Models

First of all, it is important to note that not all seemingly non-linear dependencies of a parental behavior from offspring age need a non-linear model. This is the case because models fitted with a link function other than identity link always fit a predictor-response relationship which is not linear in response space (i.e., when considering the response itself). For instance, in a logit-link model, the *linear predictor* ($LP$) has a linear relation to offspring age (i.e., in the simplest case something like $LP = c_0 + c_1 \cdot age$), but this linear predictor is then transformed to reveal the fitted values ($FV$) in *response space*. In case of the logit link function this transformation is $FV = \frac{e^{LP}}{1+e^{LP}}$. This transformation leads to predictor-response relationship appearing somewhat sigmoidal in response space. Hence, although the predictor-response relationship appears non-linear, it is linear in link space, and a standard linear model might be fully appropriate. For this reason we could conduct our simulations using a behavior-offspring age relationship as depicted in Fig. 2 of the main text and fit models to the simulated data using a standard logit link model. Figure SI 26a and Fig. SI 27a also show situations in which a linear model would be appropriate. In other cases, however, one might need a truly non-linear model.

Modelling truly non-linear relationships between parental behavior and offspring age comes with two major challenges. One of them is the need to define a mathematical equation which allows to capture the hypothesized or likely impact of offspring age on the parental behavior. The other is to actually fit the model.

Finding a mathematical equation which allows to capture the hypothesized or likely impact of offspring age on the response variable begins with consulting the available literature and thinking about how the offspring-ge dependent trajectory of the parental behavior could be shaped. In case of ontogenetic trajectories, we believe that mainly two or three principal processes seem likely. The first one is somewhat sigmoidal and the second exhibits a peak somewhere between minimum and maximum offspring age. Finally, it seems possible that some trajectories have the shape of an exponential function.

Sigmoidal trajectories have a lower and an upper asymptote; that is, the parental behaviour changes relatively little when the offspring is relatively young and also when the offspring is relatively old, and it changes relatively quickly when the offspring is of intermediate age. The proportion of time a primate is in body contact with its mother could, for instance, follow such a trajectory. In case of proportion time in body contact between mother and offspring it might further be that this happens about 100% of the time when the offspring is newborn and goes down to essentially 0% when the offspring matured. In such a case one might not even need a non-linear model as the logit link model typically applied for response variables being a proportion has a lower and upper asymptote of 0 and 1, respectively, anyway (Fig. 26a). However, there are at least two scenarios in which a standard logit link model would not work. One is a scenario in which the trajectory against age is not rotation symmetric[1] around the inflection point. In such a case, one could consider, for instance, a Gompertz function (Fig. SI 26b). The other is one in which the lower asymptote is larger than 0 and/or the upper asymptote is not 1. In such a case one might consider a sigmoidal function (Fig. SI 26c). In both of these cases one will need a genuinely non-linear model.

The second principal scenario is one in which the parental behavior peaks at intermediate offspring age (Fig. SI 27). For instance, the number of times a parent grabs its offspring in a given observation period might peak when the offspring is of intermediate age and begins to explore its environment. The most simple scenario is one in which the trajectory is mirror symmetric[2] and for which the observed behavior approaches

[1] a geometrical shape is rotation symmetric if it can be rotated by an angle other than 360° without changing its appearance. For instance, a rectangle can be turned by 180° without changing its appearance

[2] a geometrical shape is mirror symmetric if it can be reflected at an internal axis without changing its appearance. For

negative infinity (in link space) or zero (in response space or after transforming the response), the more offspring age deviates from the 'optimal' age at which the behavior peaks[3]. In such a case, one could simply add offspring age squared in addition to offspring age into the model (Fig. SI 27a). However, there are many scenarios under which such a simple model would not be appropriate. For instance, the parental behavior might be at about zero for very young offspring, peak at intermediate offspring ages, and finally level off at a value above 0 (Fig. SI 27c). The number of friendly approaches by the parent towards an offspring of the philopatric sex might, for instance, behave in such a way. There is a multitude of functions allowing to model such trajectories and the choice of the function depends on the specific nature of the trajectory of the parental behavior against age (e.g., whether the behavior levels off at about zero when the offspring ages or at a higher value or whether the parental behavior under consideration increases relatively slowly after the birth of the offspring (Fig. SI 27a, c, e) or kicks of rapidly (Fig. SI 27b, d, and f)). When searching for an appropriate mathematical function describing such a relationship, Bolker (2008) is a good start, but one needs to be aware that more potential functional relationships do exist (e.g., Marumo et al., 2022).

Finally, it might be worth mentioning exponential functions. For instance, a parental behavior might be common when the offspring is very young, monotonously decrease from offspring birth on, and finally level of at a value of zero or above (Fig. SI 28). If the parental behaviour is counted and approaches zero as the offspring gets older, a standard linear model with log-link function might be fully appropriate, but if the the behavior levels of at a value above zero as the offspring matures such a model will not do an appropriate job. In case the parental behavior is $\geq 0$, continuously varying, and approaches zero as the offspring matures (e.g., a latency), log-transforming the behavior might linearize the relation and a standard linear model (with identity link function) might appropriately model the relationship

The second major challenge is fitting the model. In case of a general (i.e., Gaussian) model with identity link, one might try the function nlmer of the package lme4 (Bates et al., 2015). However, from my (RM) experience one frequently encounters serious convergence issues that are hard to resolve. An alternative, and, to my knowledge, the only one which provides all of the potentially required flexibility with regard to the available error functions, potentially nested or partly crossed grouping factors (a.k.a. 'random effects factors'), etc. is the stan environment (Stan Development Team, 2023). Its use is easier accessible when interfacing it from R (R Core Team, 2023) via the package brms (Bürkner, 2017; Bürkner, 2018). From my (limited) experience, an attempt to fit a general or Generalized non-linear mixed model is more likely to be successful when using brms to generate the stan syntax, and then compile and run it with the aid of the package cmdstanr (Gabry et al., 2023). In either case, one has to specify carefully chosen priors, and the attempt to fit the model might also benefit from setting reasonable starting values. A further issue is how to constrain the model parameters. In fact, in most non-linear functions some or even all of its parameters are constrained to certain values. For instance, a parameter might need to be positive or limited to be between zero and one. It might also be that two parameters might be both limited to a space between zero and

instance, a square can be reflected at its diagonals and also at lines intersecting the midpoints of two opposing sides without changing its appearance

[3]in principle its also possible that the paternal behavior is the least common when the offspring is of intermediate age, but we cannot think of a practical scenario in which this seems likely

one and furthermore such that their sum is at most one. Such constraints can usually be ensured in two possible ways. One is to use appropriately constrained priors; the other one is to let the parameters vary in an unconstrained space but within the model fitting function transform them such that they are constrained appropriately (Bolker, 2008).

After the model is fitted, one has to carefully check whether the fitted offspring-age dependent trajectory of the observed parental behavior appropriately captures the actually observed parental behavior. This check is required because of at least two reasons. First, fitting non-linear models can terribly fail, and from my experience, whatever approach one uses to fit such a model, one can end with a fit model which has more or less nothing to do with the actual trajectory. Such a failure is probably best ruled out by plotting the actual data together with the fitted relationship between parental behavior and offspring age. Second, the implemented mathematical function basically represents a hypothesis (e.g., 'the ontogenetic trajectory of behavior X follows a sigmoidal shape'), which might or might not hold. Plotting the actual data together with the fitted relationship between parental behavior and offspring age will give an idea about how well the mathematical function used captures the relationship. However, plotting residuals of the model against offspring age likely is a more sensible option. This is because such a plot potentially reveals that at certain ages residuals are mostly positive or mostly negative, which would be indicative of the modelled functional relationship not matching the ontogenetic trajectory. Furthermore it is be a good idea to conduct some posterior predictive checks which allow to identify potential mismatches between the assumed and actual distribution of the response.

A question the reader might have is why we do not consider Generalized Additive Mixed Models (GAMMs) or Lowess functions here. The reasons for this are twofold. First, non-linear models as described so far have interpretable and informative model parameters (see FIG. SI 26, SI 27, and SI 28). As such, the coefficients of a fitted model translate into quantitative statements about the offspring-age dependent trajectory of the parental behavior (e.g. 'the parental behavior peaked at an offspring age of about ... months (CI: ... to ...) and from then on decreased and leveled off at a value of ... (CI: ... to ...) which was reached at an age of about ... months'). To our knowledge, a GAMM or lowess function might reveal a nicely fitting model, but it does not allow for such statements. The second, and maybe more important is that GAMMs or lowess function tend to over-fit and reveal impossible trajectories. Such over-fitted models can be perhaps best seen in the literature about growth (of bones) in which one quite frequently encounters fitted models, which indicate that individuals did begin shrinking soon after they matured – a result which is obviously biologically meaningless and which a carefully chosen proper non-linear model cannot reveal.

Another question the reader might have is why we do not consider polynomial functions other than a second order polynomial (i.e., offspring age squared; see Fig. SI 27a). In fact, quite regularly researchers wonder whether higher order polynomials (of, e.g., third or forth order) would allow for modelling more complex dependencies between, for instance, a parental behavior and offspring age. The reason why we did not consider higher order polynomials here is that higher order polynomials are rarely, if ever at all, a sensible approach. This because polynomial functions never approach an asympote but rather invariably move towards negative or positive infinity (in link space) with increasing and also with decreasing age. At the same time, though, we cannot think of a parental behavior that could follow a trajectory that moves towards (positive or negative) infinity when moving towards relatively small and also when moving towards relatively large values of offspring age. Furthermore, polynomial functions usually represent trajectories which seem biologically implausible. For instance, a third order polynomial could first increase, then decrease, and then increase again as an offspring gets older. For these two reasons we did not consider the option of using polynomials of an order higher than two here.

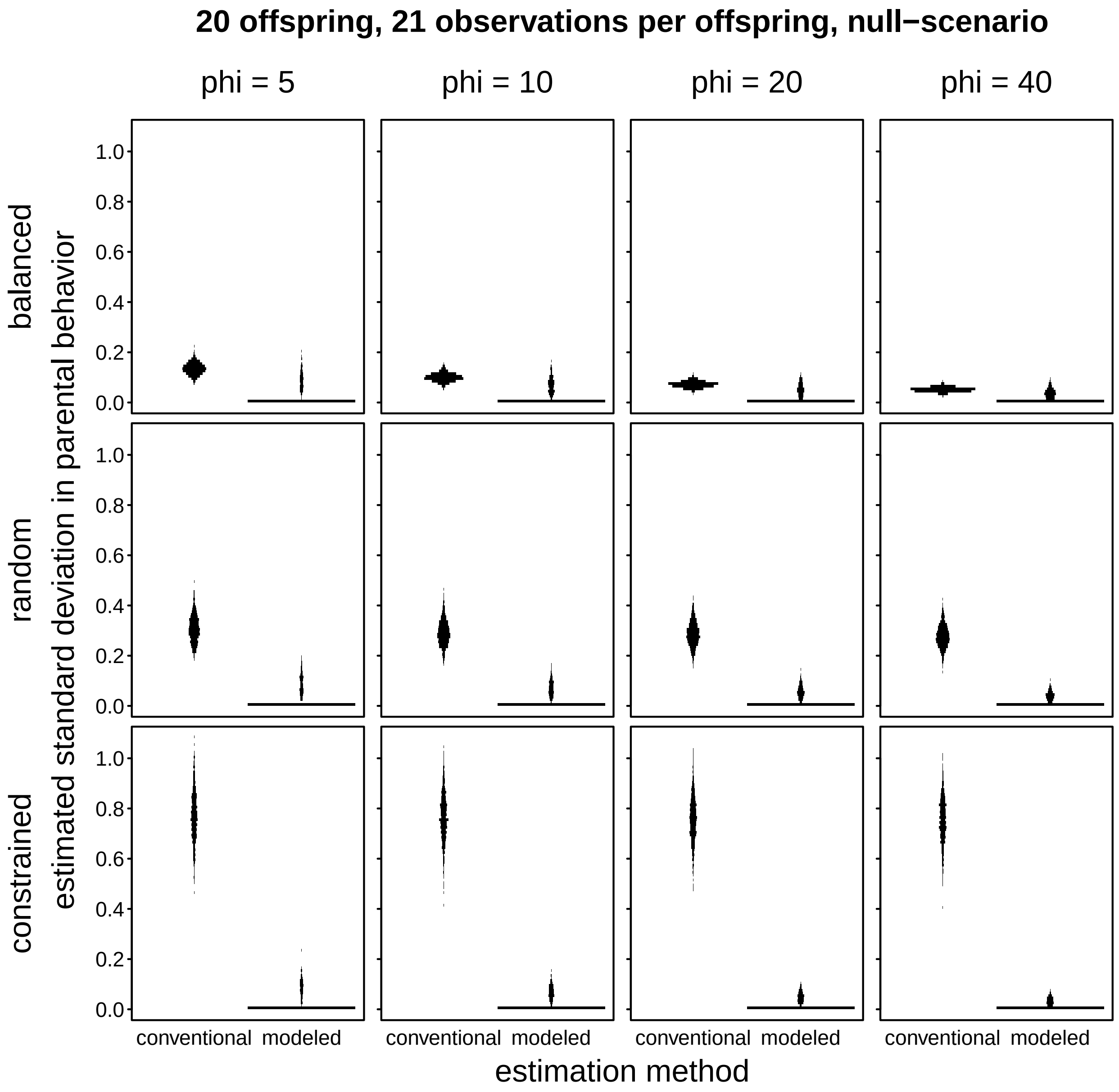


Figure SI 1. Standard deviation of conventionally estimated and modeled parent-specific behaviors, separately for each of the four values of the precision parameter *phi* (columns) and sampling schemes (rows) we simulated. The length of the horizontal line segments depicts the number of simulations (total of 1000 per plot) revealing a standard deviation in the respective bin. The simulated scenario is one in which parents did not differ at all in their parental behavior (null-scenario); sample size was 21 observations per each of 20 parents. The standard deviations depicted for *conventional* are based on the average behaviors per parent transformed to link space, and those for *modeled* represent the estimated standard deviation (in link space) for variation among parents (results for the random intercepts effect of parent-offspring ID in a GLMM). Note that the conventional approach consistently overestimated the magnitude of variation in parental behavior (which was simulated to be zero). Also the modeled standard deviation occasionally overestimated the magnitude of variation in parental behavior, but to a much lesser extent. Note that the magnitude of variation in the conventional estimates of parent-specific behavior slightly decreased with decreasing magnitude of stochastic variation in the observed proportions (increasing value of the precision parameter *phi*; left to right). Furthermore, it was lowest in case of balanced sampling and highest in case of constrained sampling (top to bottom).

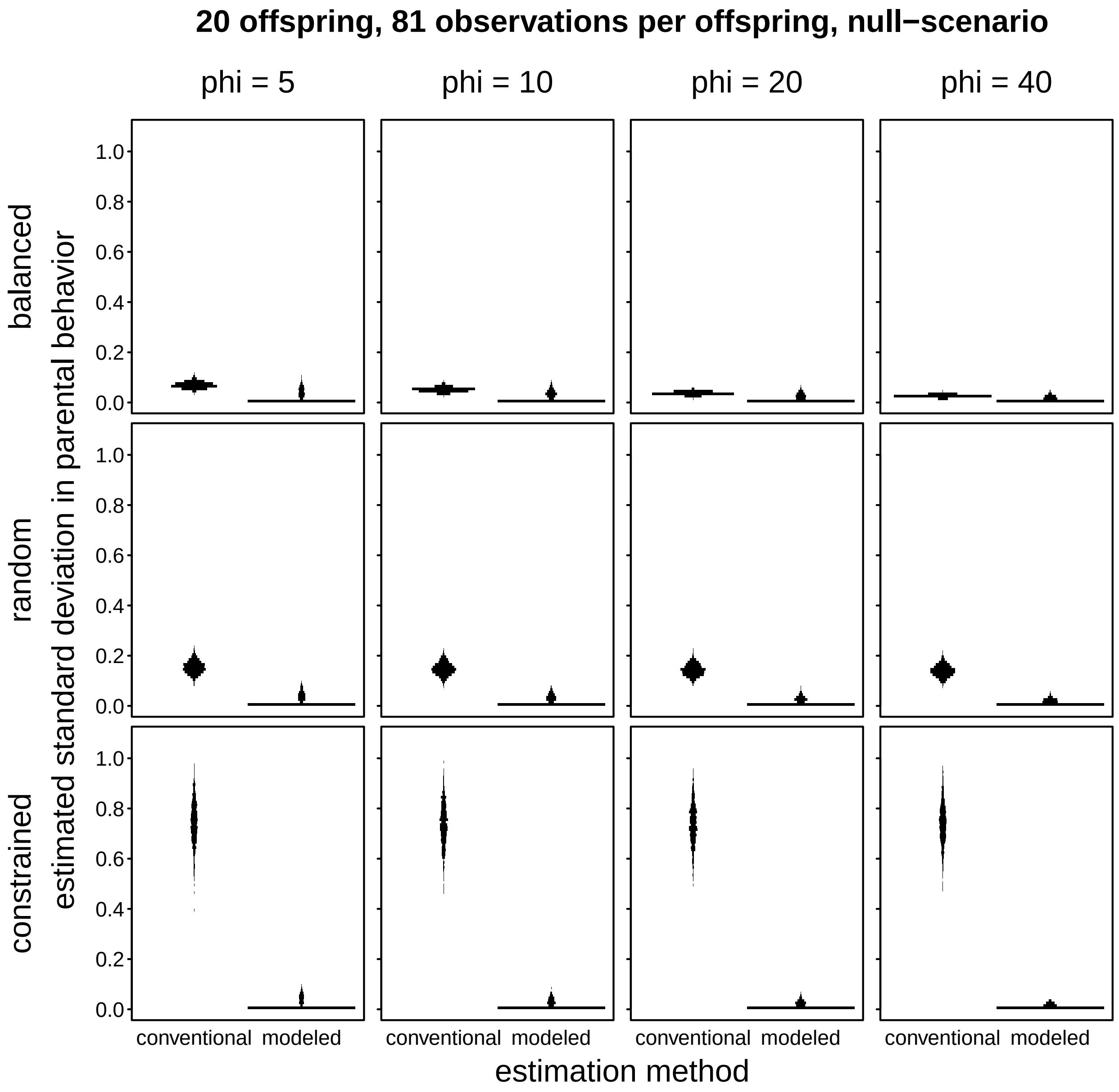


Figure SI 2. Standard deviation of conventionally estimated and modeled parent-specific behaviors, separately for each of the four values of the precision parameter *phi* (columns) and sampling schemes (rows) we simulated. The length of the horizontal line segments depicts the number of simulations (N=1000) revealing a standard deviation in the respective bin. The simulated scenario is one in which parents did not differ at all in their parental behavior (null-scenario); sample size was 81 observations per each of 20 parents. The standard deviations depicted for *conventional* are based on the average behaviors per parent transformed to link space, and those for *modeled* represent the estimated standard deviation (in link space) for variation among parents (results for the random intercepts effect of parent-offspring ID in a GLMM). Note that the conventional approach consistently overestimated the magnitude of variation in parental behavior (which was simulated to be zero). Also the modeled standard deviation occasionally overestimated the magnitude of variation in parental behavior, but to a much lesser extent. Note that the magnitude of variation in the conventional estimates of parent-specific behavior slightly decreased with decreasing magnitude of stochastic variation in the observed proportions (increasing value of the precision parameter *phi*; left to right). Furthermore, it was lowest in case of balanced sampling and highest in case of constrained sampling (top to bottom).

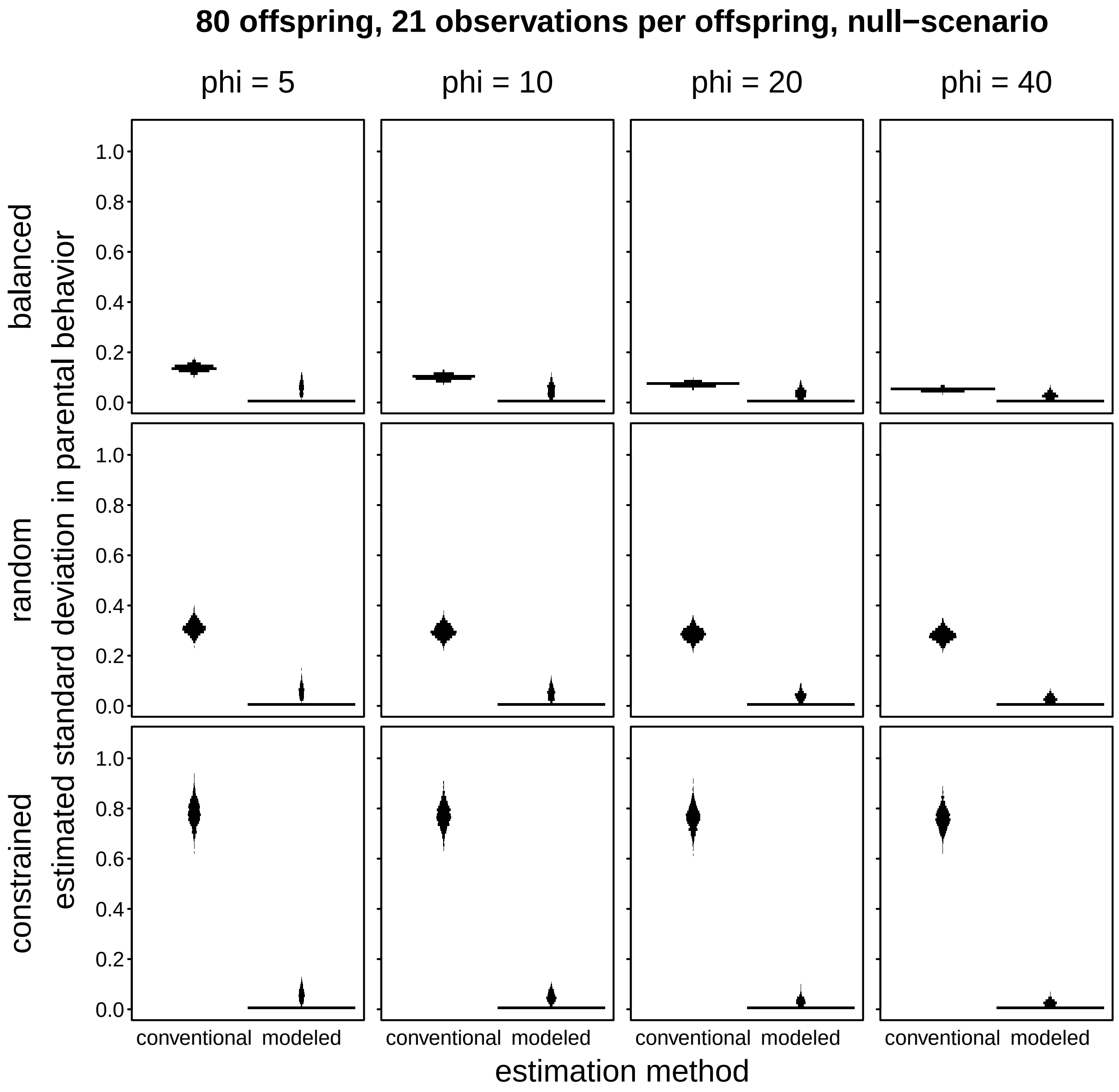


Figure SI 3. Standard deviation of conventionally estimated and modeled parent-specific behaviors, separately for each of the four values of the precision parameter *phi* (columns) and sampling schemes (rows) we simulated. The length of the horizontal line segments depicts the number of simulations (N=1000) revealing a standard deviation in the respective bin. The simulated scenario is one in which parents did not differ at all in their parental behavior (null-scenario); sample size was 21 observations per each of 80 parents. The standard deviations depicted for *conventional* are based on the average behaviors per parent transformed to link space, and those for *modeled* represent the estimated standard deviation (in link space) for variation among parents (results for the random intercepts effect of parent-offspring ID in a GLMM). Note that the conventional approach consistently overestimated the magnitude of variation in parental behavior (which was simulated to be zero). Also the modeled standard deviation occasionally overestimated the magnitude of variation in parental behavior, but to a much lesser extent. Note that the magnitude of variation in the conventional estimates of parent-specific behavior slightly decreased with decreasing magnitude of stochastic variation in the observed proportions (increasing value of the precision parameter *phi*; left to right). Furthermore, it was lowest in case of balanced sampling and highest in case of constrained sampling (top to bottom).

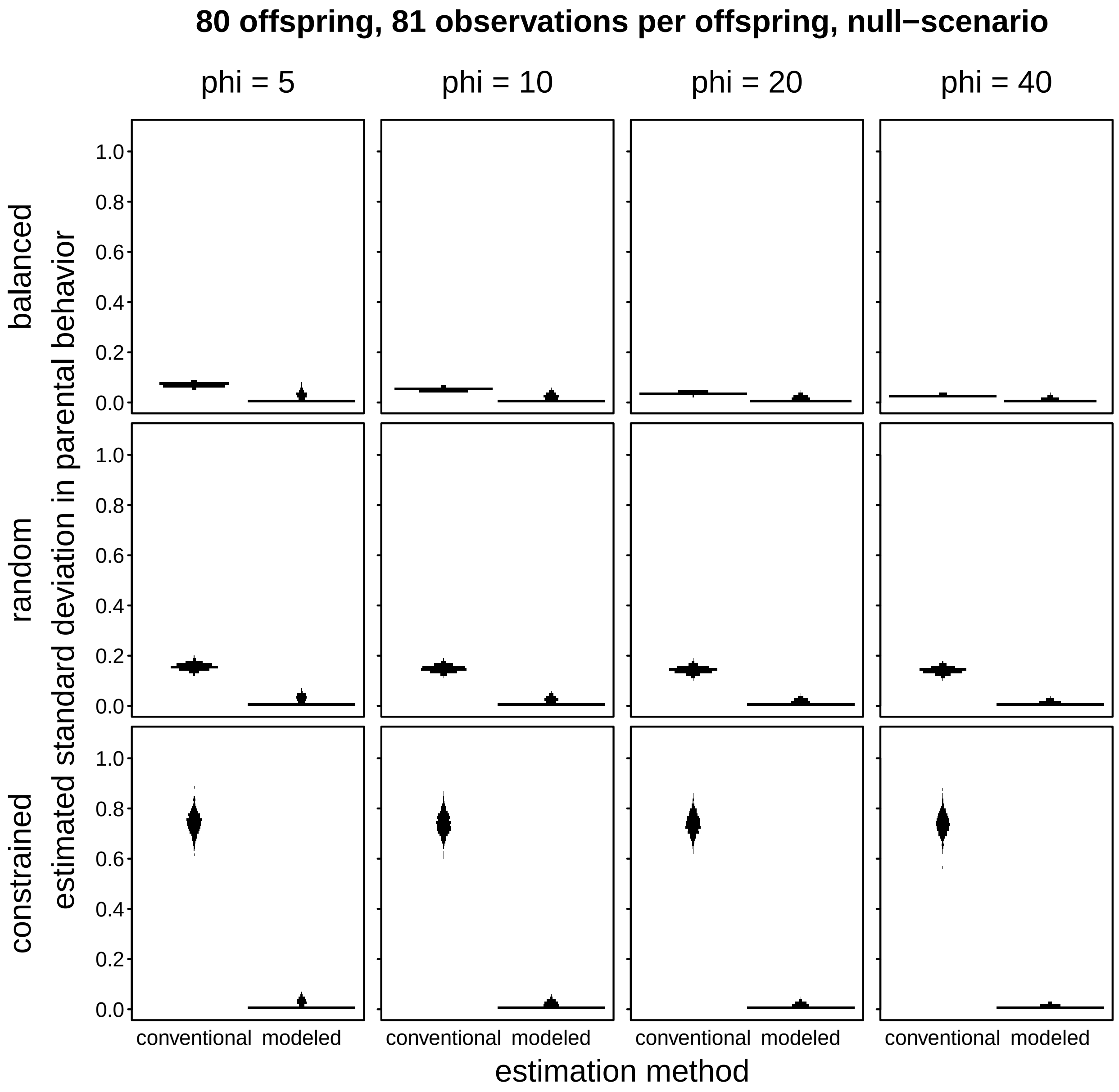


Figure SI 4. Standard deviation of conventionally estimated and modeled parent-specific behaviors, separately for each of the four values of the precision parameter *phi* (columns) and sampling schemes (rows) we simulated. The length of the horizontal line segments depicts the number of simulations (N=1000) revealing a standard deviation in the respective bin. The simulated scenario is one in which parents did not differ at all in their parental behavior (null-scenario); sample size was 81 observations per each of 80 parents. The standard deviations depicted for *conventional* are based on the average behaviors per parent transformed to link space, and those for *modeled* represent the estimated standard deviation (in link space) for variation among parents (results for the random intercepts effect of parent-offspring ID in a GLMM). Note that the conventional approach consistently overestimated the magnitude of variation in parental behavior (which was simulated to be zero). Also the modeled standard deviation occasionally overestimated the magnitude of variation in parental behavior, but to a much lesser extent. Note that the magnitude of variation in the conventional estimates of parent-specific behavior slightly decreased with decreasing magnitude of stochastic variation in the observed proportions (increasing value of the precision parameter *phi*; left to right). Furthermore, it was lowest in case of balanced sampling and highest in case of constrained sampling (top to bottom).

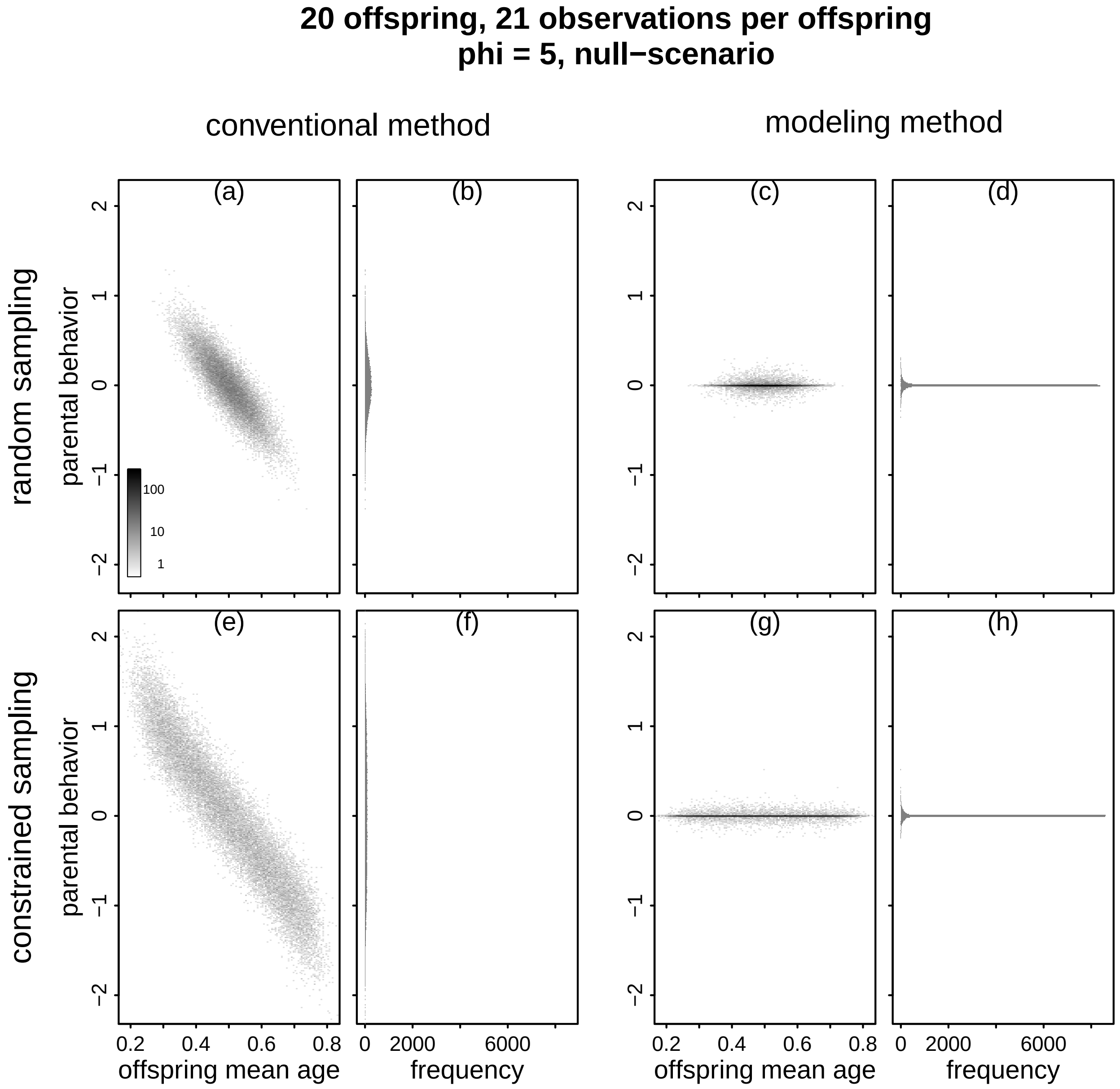


Figure SI 5. Estimated parent-specific behavior (y-axes) against the average age of an offspring across the days at which it was observed (a, c, e, g) and the frequency distribution of the estimates besides them (b, d, f, h). The left two columns depict conventional estimates of parent-specific behavior and the right two columns depict modeled estimates of parent-specific behavior. The top row shows the results in case of random sampling and the bottom row shows the results in case of constrained sampling (see Fig. 3 of the main text). The simulated parents did not differ in their parental effort. Sample size was 21 observations per each of 20 offspring, and we used a precision parameter *phi* of 5. For (a), (c), (e), and (g) we binned average offspring age (bin width: 0.01) and estimated parental effort (bin width: 0.01) and scaled the darkness of points proportionate to the logarithm of the number of estimates falling in the respective combination of binned age and estimated parent-specific effort. The histograms in (b), (d), (f), and (h) depict the frequency distribution of the estimates of parent-specific behavior shown to the left of them. Note that all y-axes are at the same scale, and x-axes are at the same scale in (a), (c), (e), and (g) as they are in (b), (d), (f), and (h). Also note that the conventional estimates of parent-specific behavior clearly decreased with increasing average age of the offspring at the observation days, whereas the modeled estimates of parent-specific behavior did not. Also note that the effect of mean offspring age on the conventional estimates of parental effort was stronger in case of constrained as compared to random sampling. Conventional estimates of parent-specific behavior are depicted in link space.

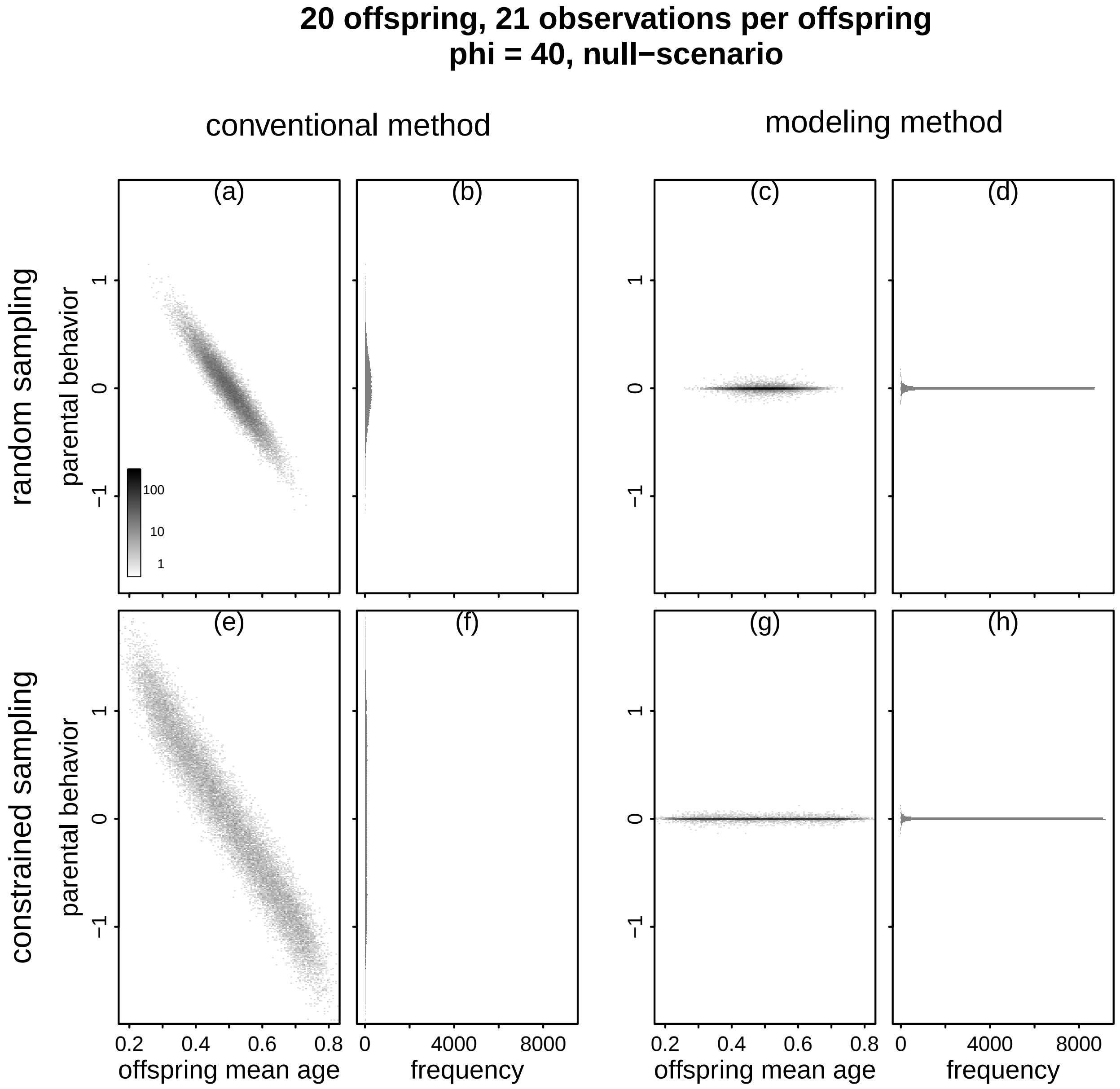


Figure SI 6. Estimated parent-specific behavior (y-axes) against the average age of an offspring across the days at which it was observed (a, c, e, g) and the frequency distribution of the estimates besides them (b, d, f, h). The left two columns depict conventional estimates of parent-specific behavior and the right two columns depict modeled estimates of parent-specific behavior. The top row shows the results in case of random sampling and the bottom row shows the results in case of constrained sampling (see Fig. 3 of the main text). The simulated parents did not differ in their parental effort. Sample size was 21 observations per each of 20 offspring, and we used a precision parameter *phi* of 40. For (a), (c), (e), and (g) we binned average offspring age (bin width: 0.01) and estimated parental effort (bin width: 0.01) and scaled the darkness of points proportionate to the logarithm of the number of estimates falling in the respective combination of binned age and estimated parent-specific effort. The histograms in (b), (d), (f), and (h) depict the frequency distribution of the estimates of parent-specific behavior shown to the left of them. Note that all y-axes are at the same scale, and x-axes are at the same scale in (a), (c), (e), and (g) as they are in (b), (d), (f), and (h). Also note that the conventional estimates of parent-specific behavior clearly decreased with increasing average age of the offspring at the observation days, whereas the modeled estimates of parent-specific behavior did not. Also note that the effect of mean offspring age on the conventional estimates of parental effort was stronger in case of constrained as compared to random sampling. Conventional estimates of parent-specific behavior are depicted in link space.

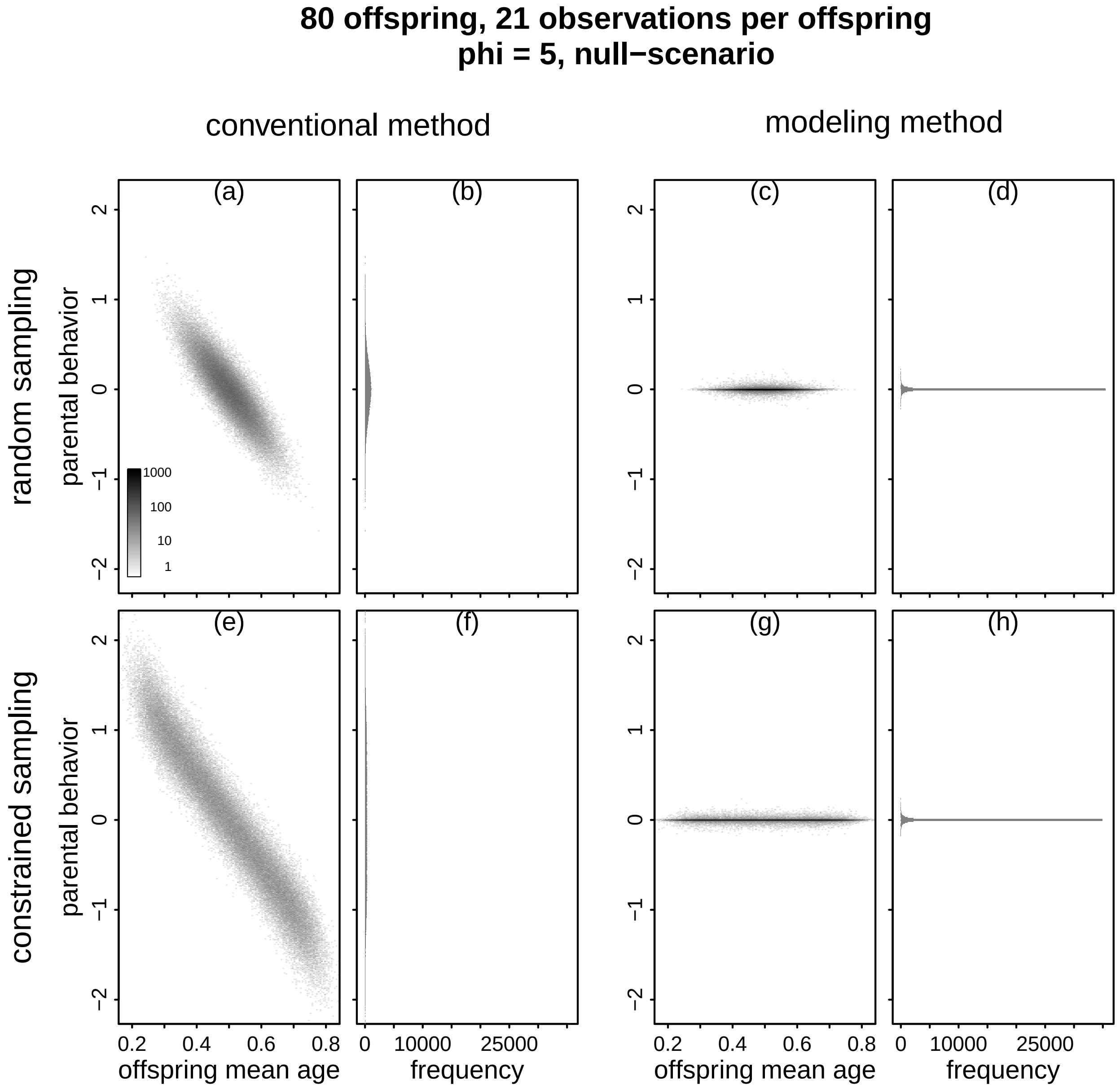


Figure SI 7. Estimated parent-specific behavior (y-axes) against the average age of an offspring across the days at which it was observed (a, c, e, g) and the frequency distribution of the estimates besides them (b, d, f, h). The left two columns depict conventional estimates of parent-specific behavior and the right two columns depict modeled estimates of parent-specific behavior. The top row shows the results in case of random sampling and the bottom row shows the results in case of constrained sampling (see Fig. 3 of the main text). The simulated parents did not differ in their parental effort. Sample size was 21 observations per each of 80 offspring, and we used a precision parameter *phi* of 5. For (a), (c), (e), and (g) we binned average offspring age (bin width: 0.01) and estimated parental effort (bin width: 0.01) and scaled the darkness of points proportionate to the logarithm of the number of estimates falling in the respective combination of binned age and estimated parent-specific effort. The histograms in (b), (d), (f), and (h) depict the frequency distribution of the estimates of parent-specific behavior shown to the left of them. Note that all y-axes are at the same scale, and x-axes are at the same scale in (a), (c), (e), and (g) as they are in (b), (d), (f), and (h). Also note that the conventional estimates of parent-specific behavior clearly decreased with increasing average age of the offspring at the observation days, whereas the modeled estimates of parent-specific behavior did not. Also note that the effect of mean offspring age on the conventional estimates of parental effort was stronger in case of constrained as compared to random sampling. Conventional estimates of parent-specific behavior are depicted in link space.

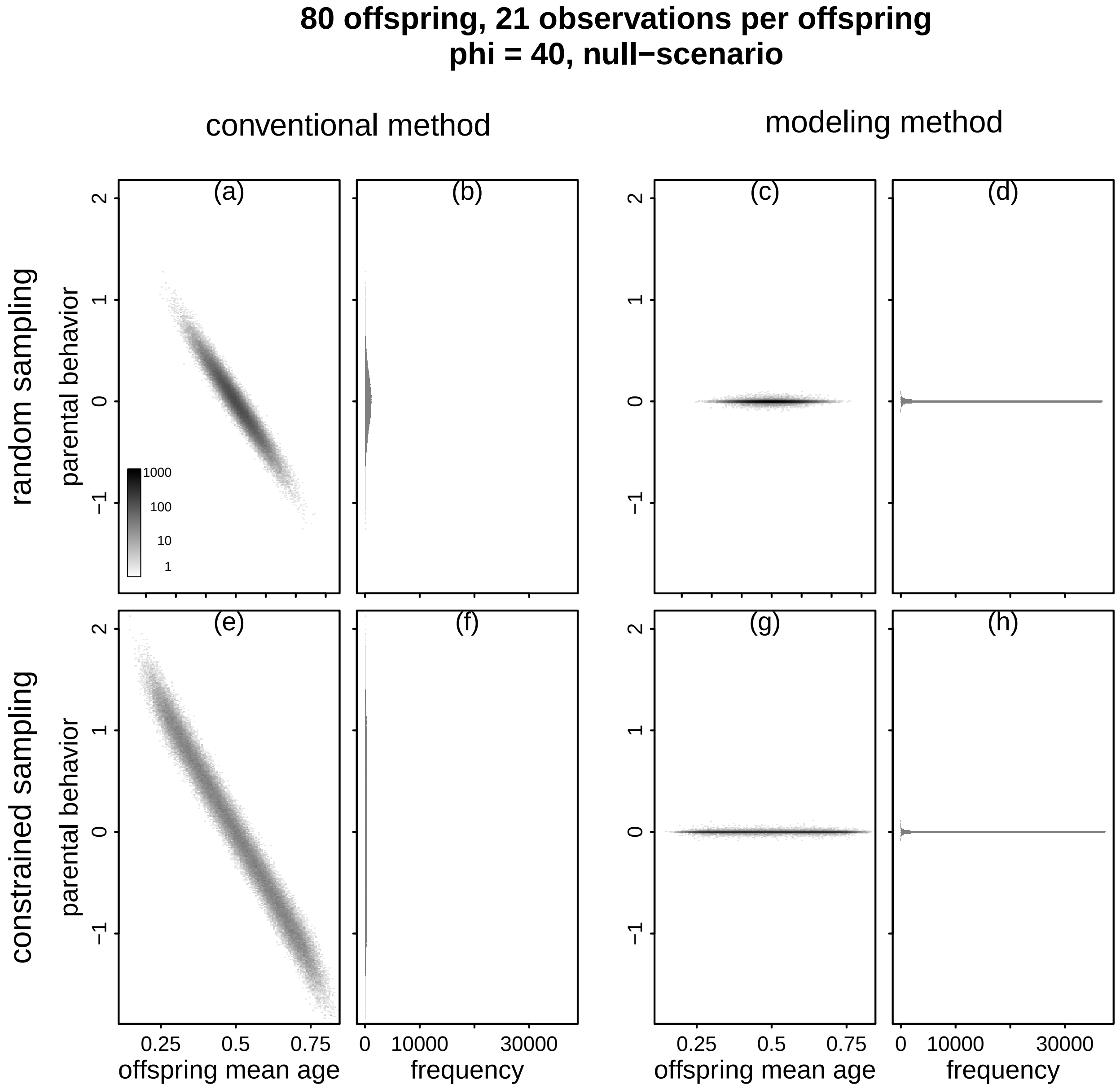


Figure SI 8. Estimated parent-specific behavior (y-axes) against the average age of an offspring across the days at which it was observed (a, c, e, g) and the frequency distribution of the estimates besides them (b, d, f, h). The left two columns depict conventional estimates of parent-specific behavior and the right two columns depict modeled estimates of parent-specific behavior. The top row shows the results in case of random sampling and the bottom row shows the results in case of constrained sampling (see Fig. 3 of the main text). The simulated parents did not differ in their parental effort. Sample size was 21 observations per each of 80 offspring, and we used a precision parameter *phi* of 40. For (a), (c), (e), and (g) we binned average offspring age (bin width: 0.01) and estimated parental effort (bin width: 0.01) and scaled the darkness of points proportionate to the logarithm of the number of estimates falling in the respective combination of binned age and estimated parent-specific effort. The histograms in (b), (d), (f), and (h) depict the frequency distribution of the estimates of parent-specific behavior shown to the left of them. Note that all y-axes are at the same scale, and x-axes are at the same scale in (a), (c), (e), and (g) as they are in (b), (d), (f), and (h). Also note that the conventional estimates of parent-specific behavior clearly decreased with increasing average age of the offspring at the observation days, whereas the modeled estimates of parent-specific behavior did not. Also note that the effect of mean offspring age on the conventional estimates of parental effort was stronger in case of constrained as compared to random sampling. Conventional estimates of parent-specific behavior are depicted in link space.

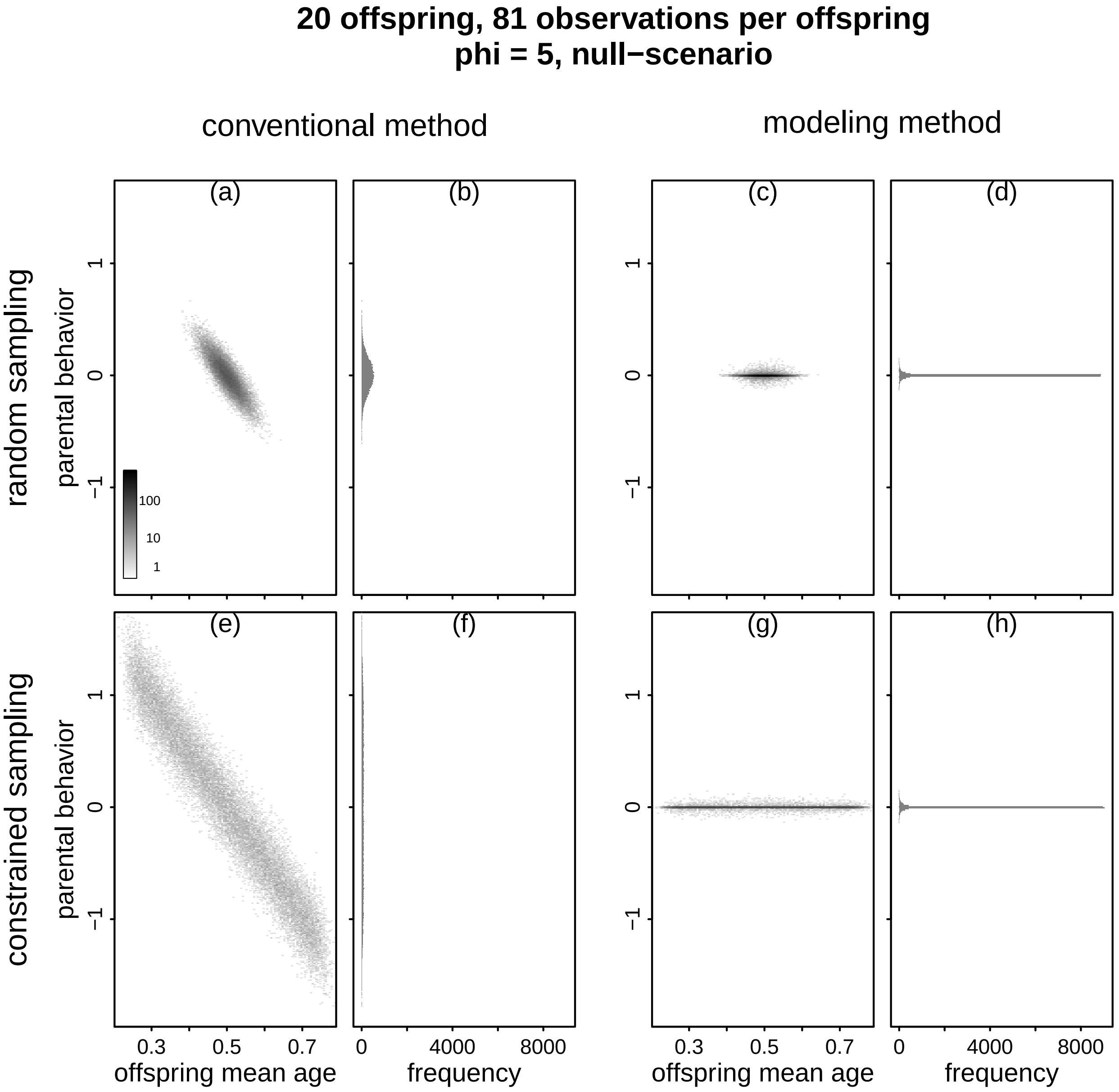


Figure SI 9. Estimated parent-specific behavior (y-axes) against the average age of an offspring across the days at which it was observed (a, c, e, g) and the frequency distribution of the estimates besides them (b, d, f, h). The left two columns depict conventional estimates of parent-specific behavior and the right two columns depict modeled estimates of parent-specific behavior. The top row shows the results in case of random sampling and the bottom row shows the results in case of constrained sampling (see Fig. 3 of the main text). The simulated parents did not differ in their parental effort. Sample size was 81 observations per each of 20 offspring, and we used a precision parameter *phi* of 5. For (a), (c), (e), and (g) we binned average offspring age (bin width: 0.01) and estimated parental effort (bin width: 0.01) and scaled the darkness of points proportionate to the logarithm of the number of estimates falling in the respective combination of binned age and estimated parent-specific effort. The histograms in (b), (d), (f), and (h) depict the frequency distribution of the estimates of parent-specific behavior shown to the left of them. Note that all y-axes are at the same scale, and x-axes are at the same scale in (a), (c), (e), and (g) as they are in (b), (d), (f), and (h). Also note that the conventional estimates of parent-specific behavior clearly decreased with increasing average age of the offspring at the observation days, whereas the modeled estimates of parent-specific behavior did not. Also note that the effect of mean offspring age on the conventional estimates of parental effort was stronger in case of constrained as compared to random sampling. Conventional estimates of parent-specific behavior are depicted in link space.

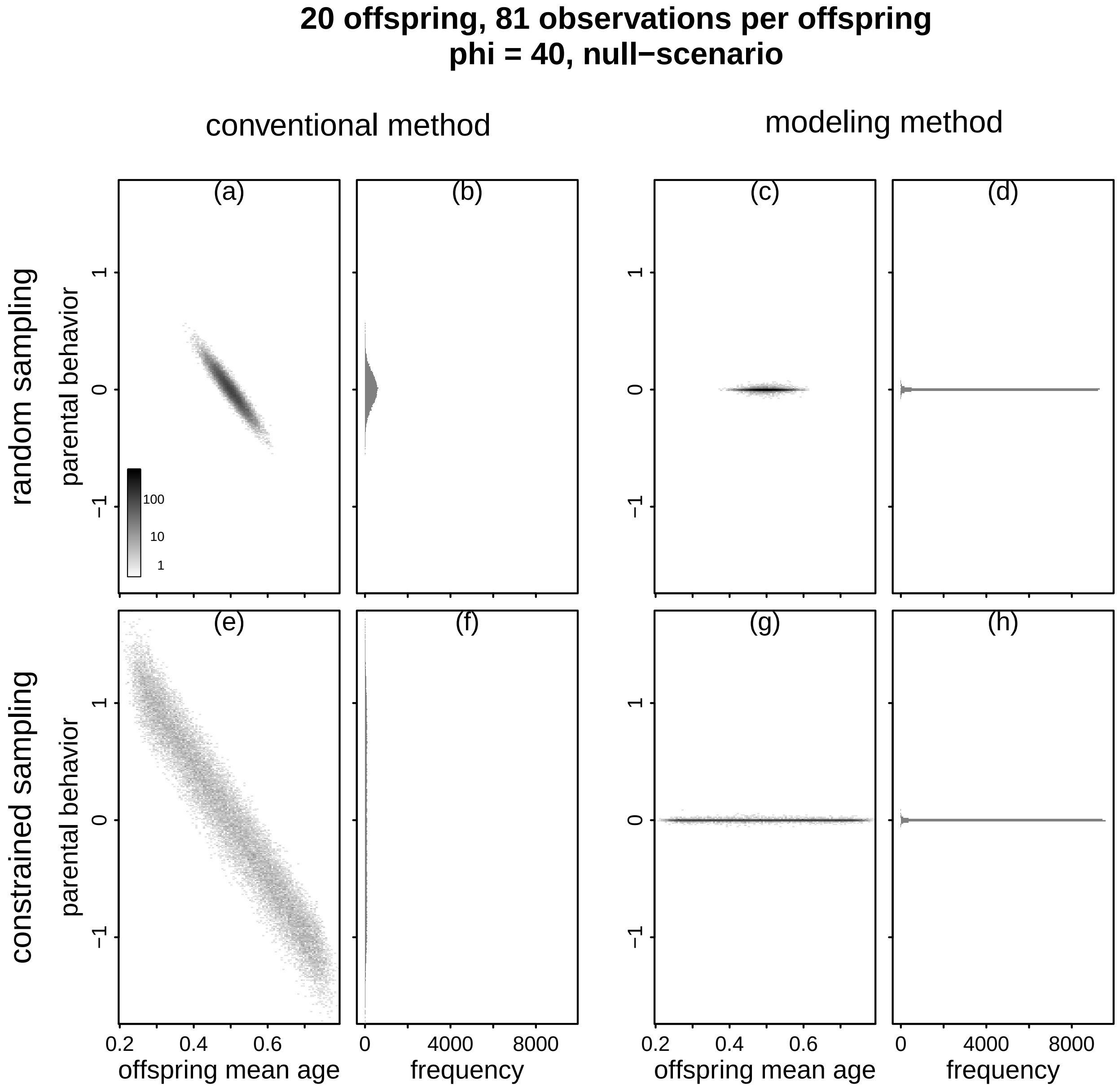


Figure SI 10. Estimated parent-specific behavior (y-axes) against the average age of an offspring across the days at which it was observed (a, c, e, g) and the frequency distribution of the estimates besides them (b, d, f, h). The left two columns depict conventional estimates of parent-specific behavior and the right two columns depict modeled estimates of parent-specific behavior. The top row shows the results in case of random sampling and the bottom row shows the results in case of constrained sampling (see Fig. 3 of the main text). The simulated parents did not differ in their parental effort. Sample size was 81 observations per each of 20 offspring, and we used a precision parameter *phi* of 40. For (a), (c), (e), and (g) we binned average offspring age (bin width: 0.01) and estimated parental effort (bin width: 0.01) and scaled the darkness of points proportionate to the logarithm of the number of estimates falling in the respective combination of binned age and estimated parent-specific effort. The histograms in (b), (d), (f), and (h) depict the frequency distribution of the estimates of parent-specific behavior shown to the left of them. Note that all y-axes are at the same scale, and x-axes are at the same scale in (a), (c), (e), and (g) as they are in (b), (d), (f), and (h). Also note that the conventional estimates of parent-specific behavior clearly decreased with increasing average age of the offspring at the observation days, whereas the modeled estimates of parent-specific behavior did not. Also note that the effect of mean offspring age on the conventional estimates of parental effort was stronger in case of constrained as compared to random sampling. Conventional estimates of parent-specific behavior are depicted in link space.

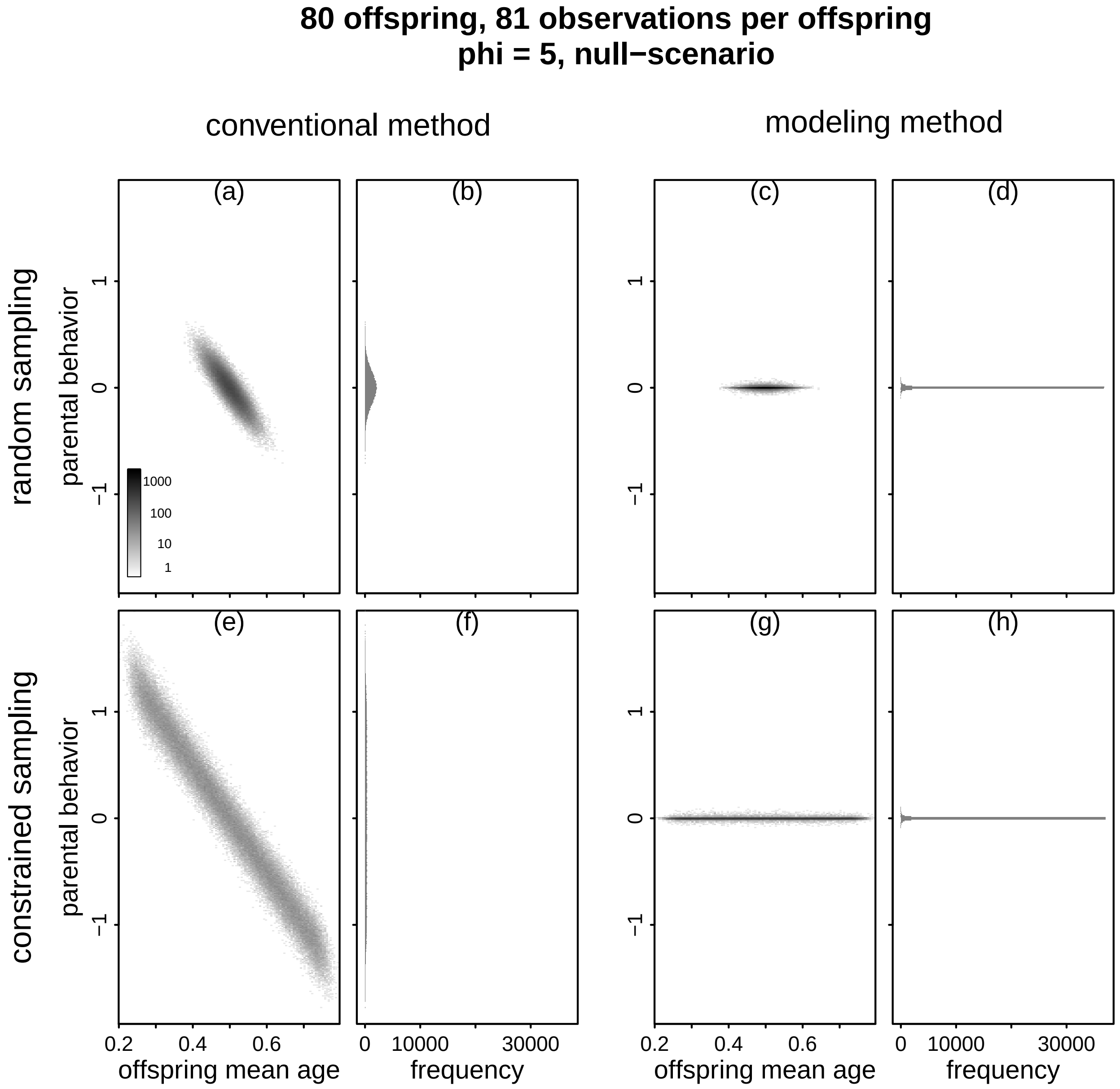


Figure SI 11. Estimated parent-specific behavior (y-axes) against the average age of an offspring across the days at which it was observed (a, c, e, g) and the frequency distribution of the estimates besides them (b, d, f, h). The left two columns depict conventional estimates of parent-specific behavior and the right two columns depict modeled estimates of parent-specific behavior. The top row shows the results in case of random sampling and the bottom row shows the results in case of constrained sampling (see Fig. 3 of the main text). The simulated parents did not differ in their parental effort. Sample size was 81 observations per each of 80 offspring, and we used a precision parameter *phi* of 5. For (a), (c), (e), and (g) we binned average offspring age (bin width: 0.01) and estimated parental effort (bin width: 0.01) and scaled the darkness of points proportionate to the logarithm of the number of estimates falling in the respective combination of binned age and estimated parent-specific effort. The histograms in (b), (d), (f), and (h) depict the frequency distribution of the estimates of parent-specific behavior shown to the left of them. Note that all y-axes are at the same scale, and x-axes are at the same scale in (a), (c), (e), and (g) as they are in (b), (d), (f), and (h). Also note that the conventional estimates of parent-specific behavior clearly decreased with increasing average age of the offspring at the observation days, whereas the modeled estimates of parent-specific behavior did not. Also note that the effect of mean offspring age on the conventional estimates of parental effort was stronger in case of constrained as compared to random sampling. Conventional estimates of parent-specific behavior are depicted in link space.

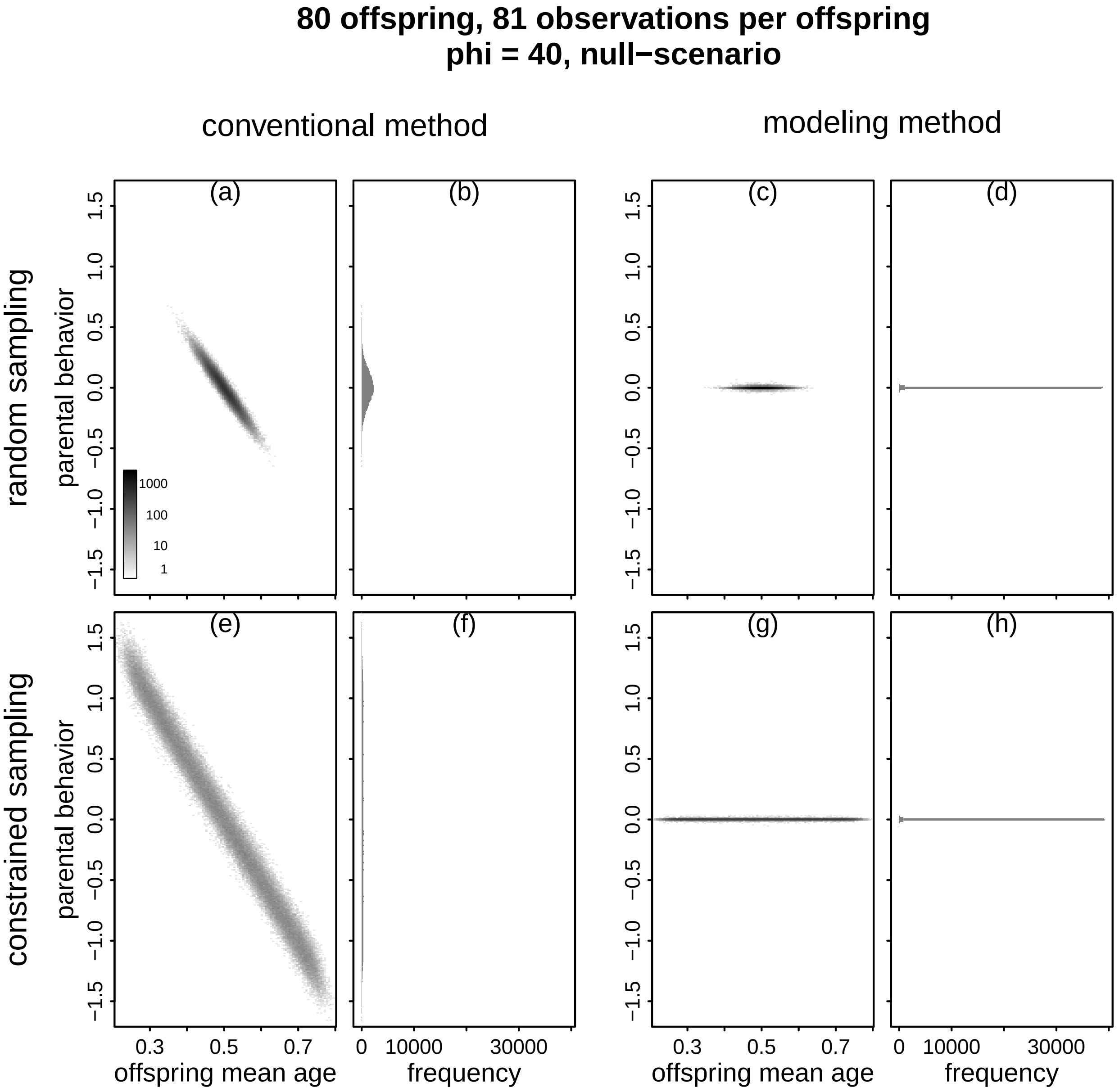


Figure SI 12. Estimated parent-specific behavior (y-axes) against the average age of an offspring across the days at which it was observed (a, c, e, g) and the frequency distribution of the estimates besides them (b, d, f, h). The left two columns depict conventional estimates of parent-specific behavior and the right two columns depict modeled estimates of parent-specific behavior. The top row shows the results in case of random sampling and the bottom row shows the results in case of constrained sampling (see Fig. 3 of the main text). The simulated parents did not differ in their parental effort. Sample size was 81 observations per each of 80 offspring, and we used a precision parameter *phi* of 40. For (a), (c), (e), and (g) we binned average offspring age (bin width: 0.01) and estimated parental effort (bin width: 0.01) and scaled the darkness of points proportionate to the logarithm of the number of estimates falling in the respective combination of binned age and estimated parent-specific effort. The histograms in (b), (d), (f), and (h) depict the frequency distribution of the estimates of parent-specific behavior shown to the left of them. Note that all y-axes are at the same scale, and x-axes are at the same scale in (a), (c), (e), and (g) as they are in (b), (d), (f), and (h). Also note that the conventional estimates of parent-specific behavior clearly decreased with increasing average age of the offspring at the observation days, whereas the modeled estimates of parent-specific behavior did not. Also note that the effect of mean offspring age on the conventional estimates of parental effort was stronger in case of constrained as compared to random sampling. Conventional estimates of parent-specific behavior are depicted in link space.

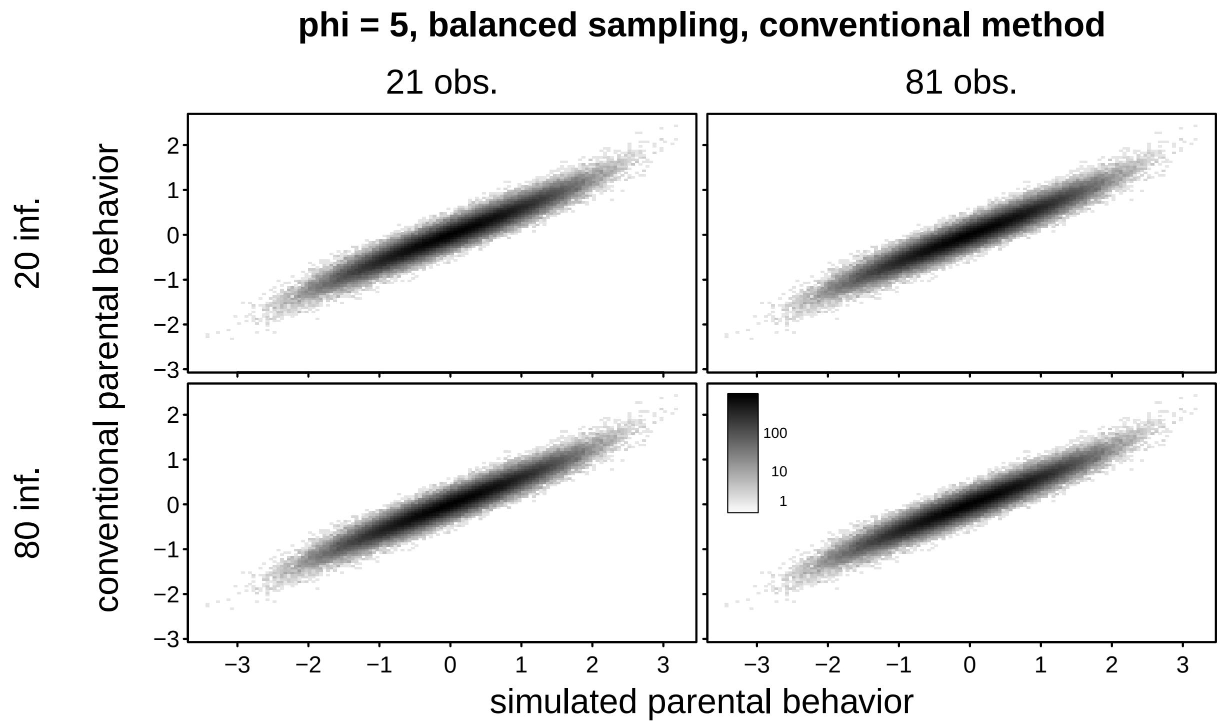


Figure SI 13. Conventional estimates of parent-specific behavior as a function of simulated parent-specific behavior, separately for the different combinations of number parent offspring dyads (rows) and number observations per dyad (columns). Sampling was balanced and the precision parameter *phi* was 5. Note that the simulated and estimated parent-specific behavior were in fairly good agreement.

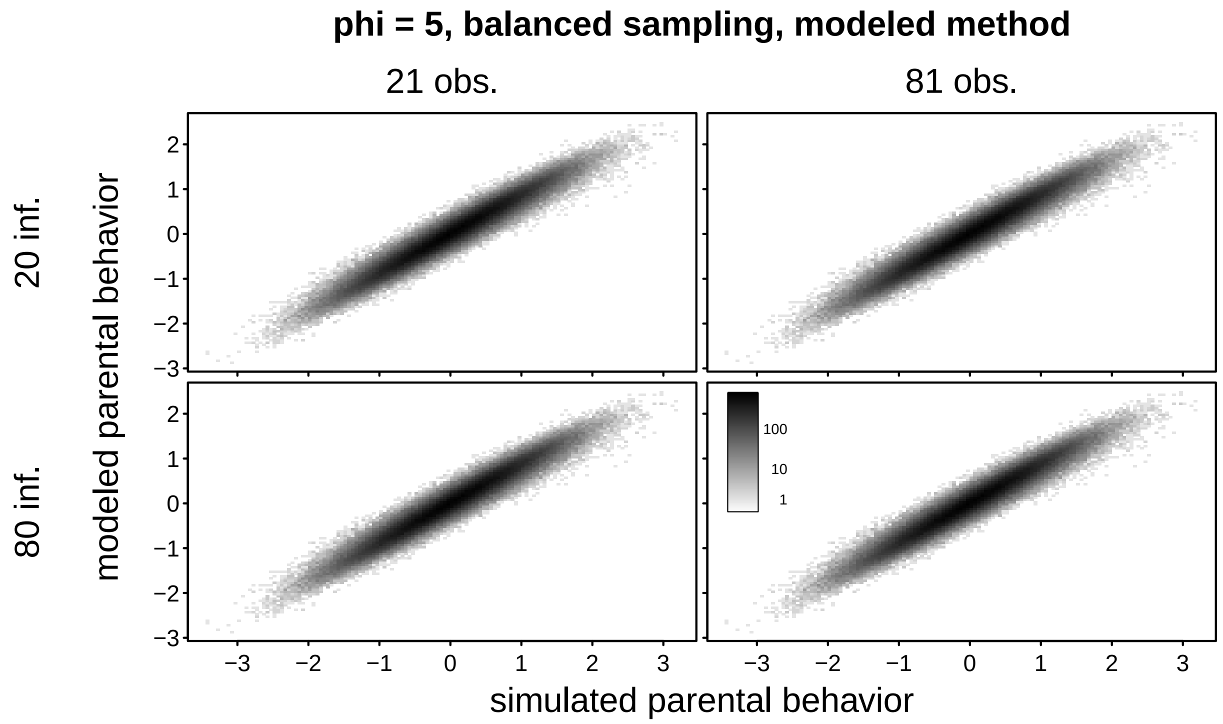


Figure SI 14. Modeled estimates of parent-specific behavior as a function of simulated parent-specific behavior, separately for the different combinations of number parent offspring dyads (rows) and number observations per dyad (columns). Sampling was balanced and the precision parameter *phi* was 5. Note that the simulated and estimated parent-specific behavior were in fairly good agreement.

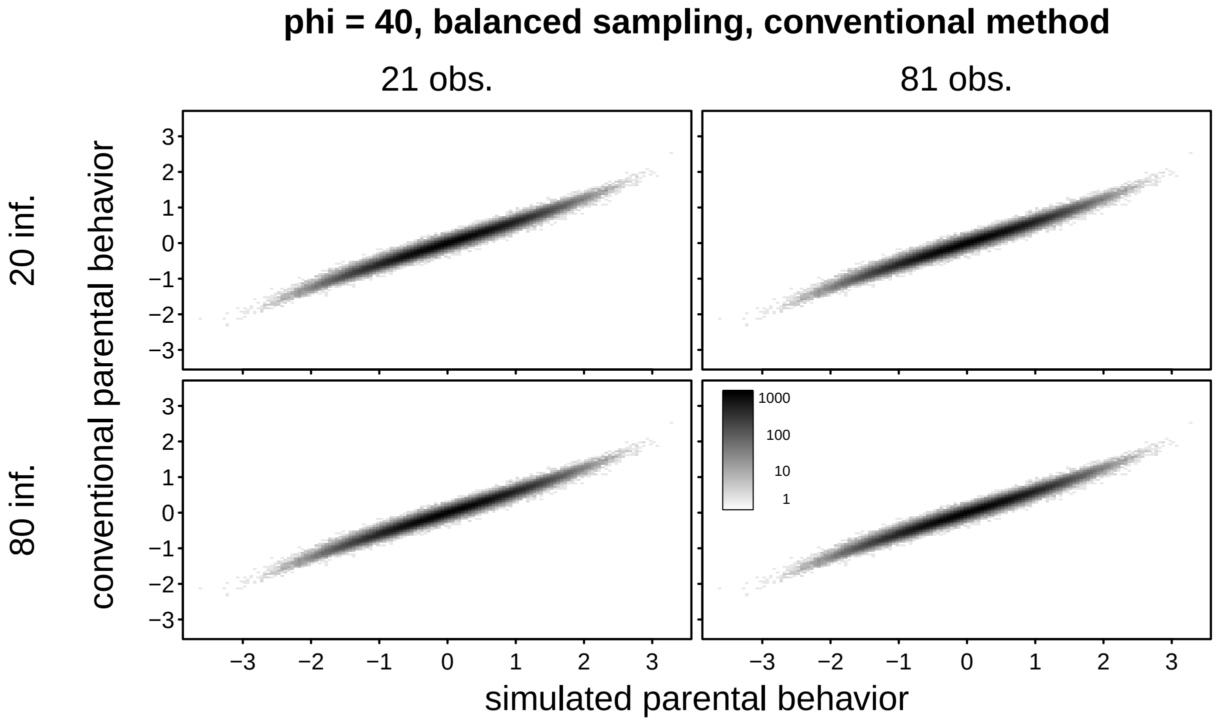


Figure SI 15. Conventional estimates of parent-specific behavior as a function of simulated parent-specific behavior, separately for the different combinations of number parent offspring dyads (rows) and number observations per dyad (columns). Sampling was balanced and the precision parameter *phi* was 40. Note that the simulated and estimated parent-specific behavior were in fairly good agreement.

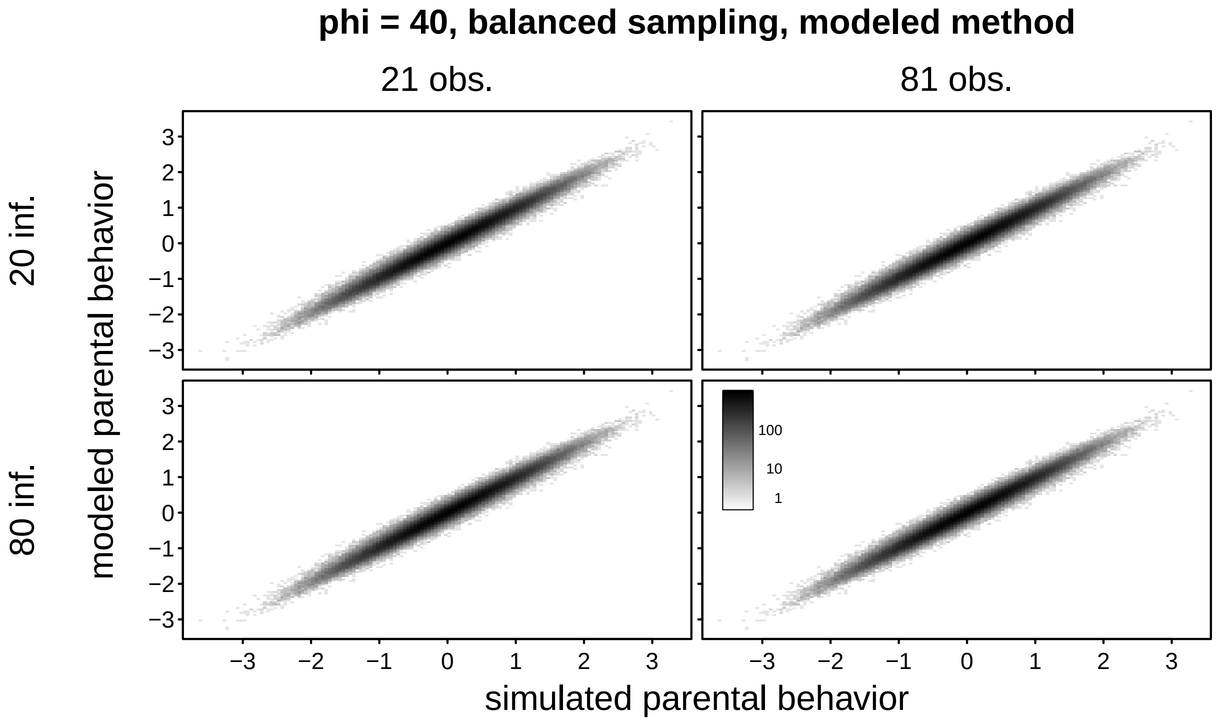


Figure SI 16. Modeled estimates of parent-specific behavior as a function of simulated parent-specific behavior, separately for the different combinations of number parent offspring dyads (rows) and number observations per dyad (columns). Sampling was balanced and the precision parameter *phi* was 40. Note that the simulated and estimated parent-specific behavior were in fairly good agreement.

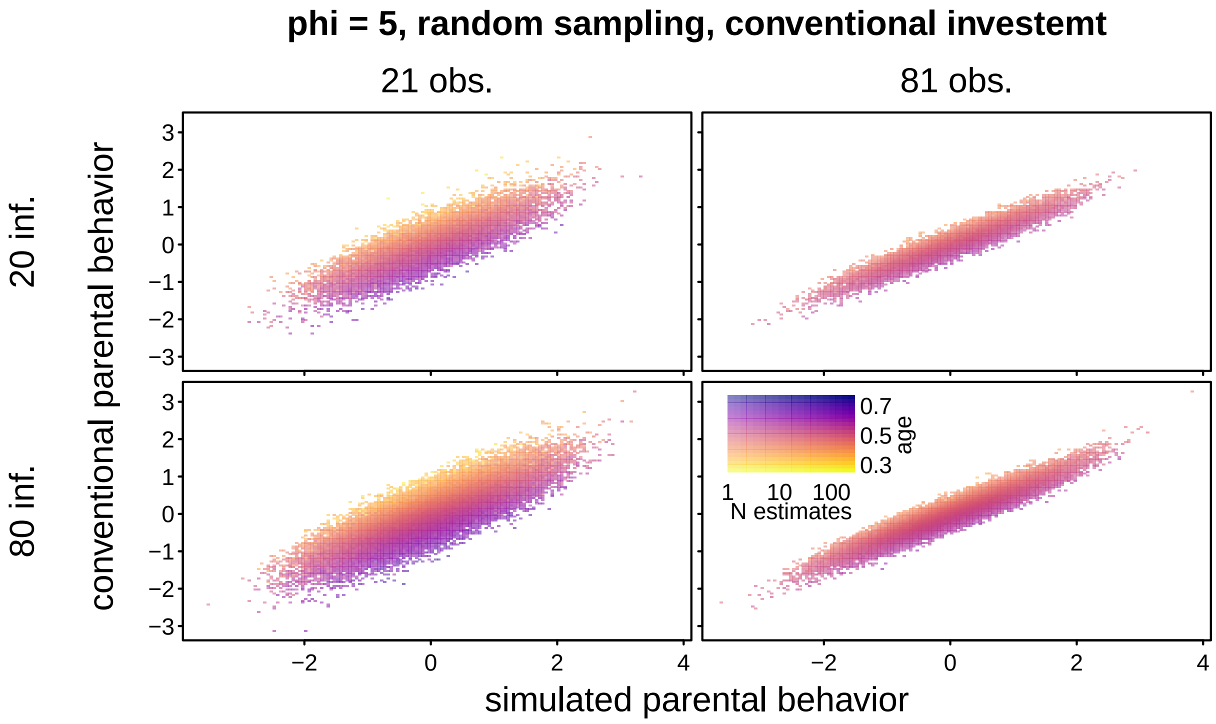


Figure SI 17. Conventional estimates of parent-specific behavior as a function of simulated parent-specific behavior, separately for the different combinations of number parent offspring dyads (rows) and number observations per dyad (columns). Sampling was random and the precision parameter *phi* was 5. The color (yellow to blue) codes average offspring age, and saturation codes number estimates per bin.

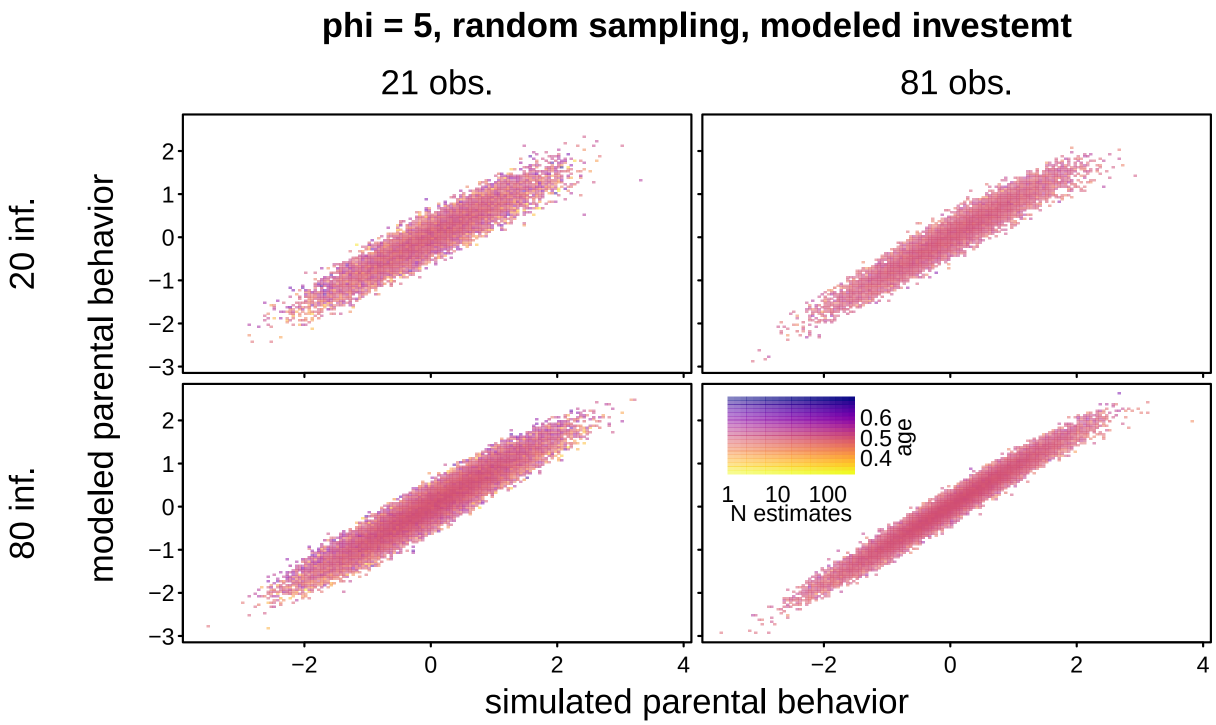


Figure SI 18. Modeled estimates of parent-specific behavior as a function of simulated parent-specific behavior, separately for the different combinations of number parent offspring dyads (rows) and number observations per dyad (columns). Sampling was random and the precision parameter *phi* was 5. The color (yellow to blue) codes average offspring age, and saturation codes number estimates per bin.

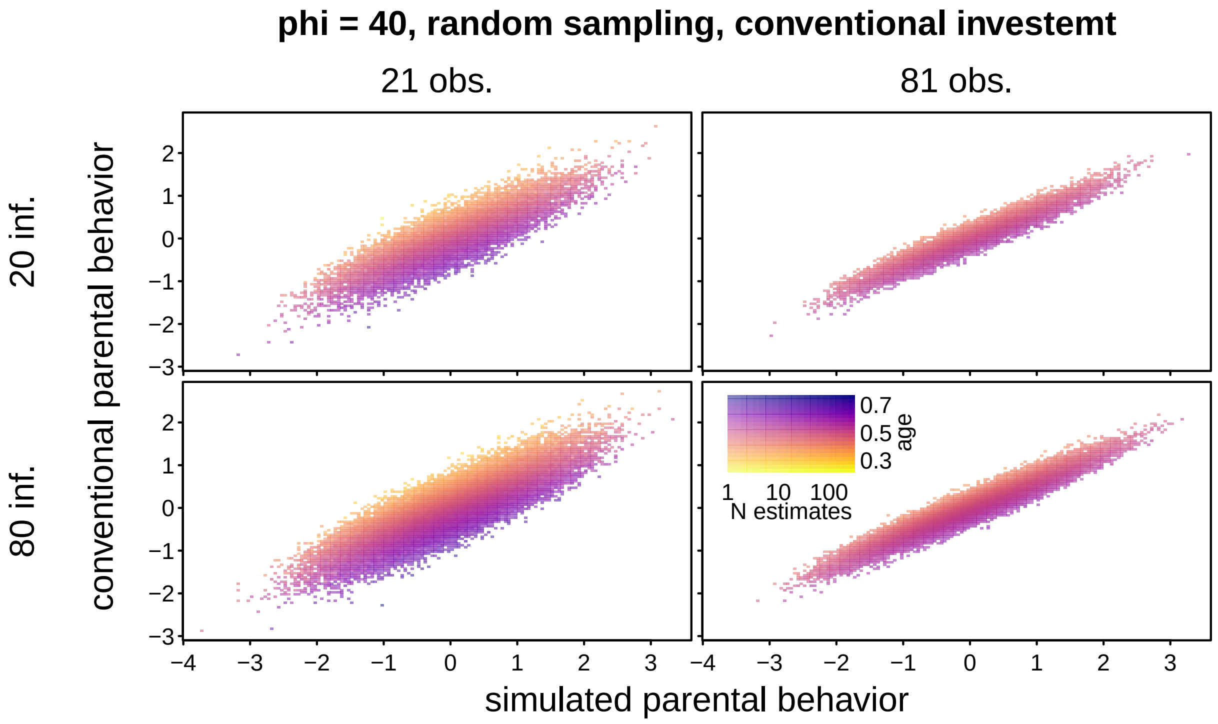


Figure SI 19. Conventional estimates of parent-specific behavior as a function of simulated parent-specific behavior, separately for the different combinations of number parent offspring dyads (rows) and number observations per dyad (columns). Sampling was random and the precision parameter *phi* was 40. The color (yellow to blue) codes average offspring age, and saturation codes number estimates per bin.

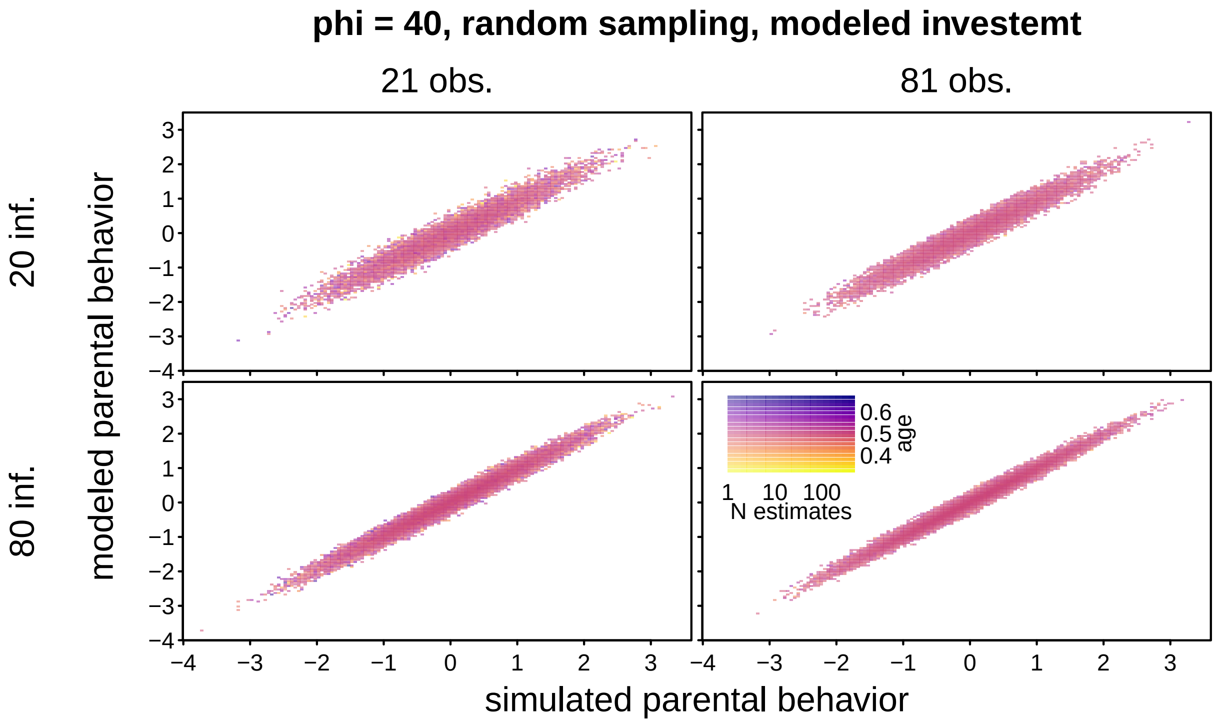


Figure SI 20 Modeled estimates of parent-specific behavior as a function of simulated parent-specific behavior, separately for the different combinations of number parent offspring dyads (rows) and number observations per dyad (columns). Sampling was random and the precision parameter *phi* was 40. The color (yellow to blue) codes average offspring age, and saturation codes number estimates per bin.

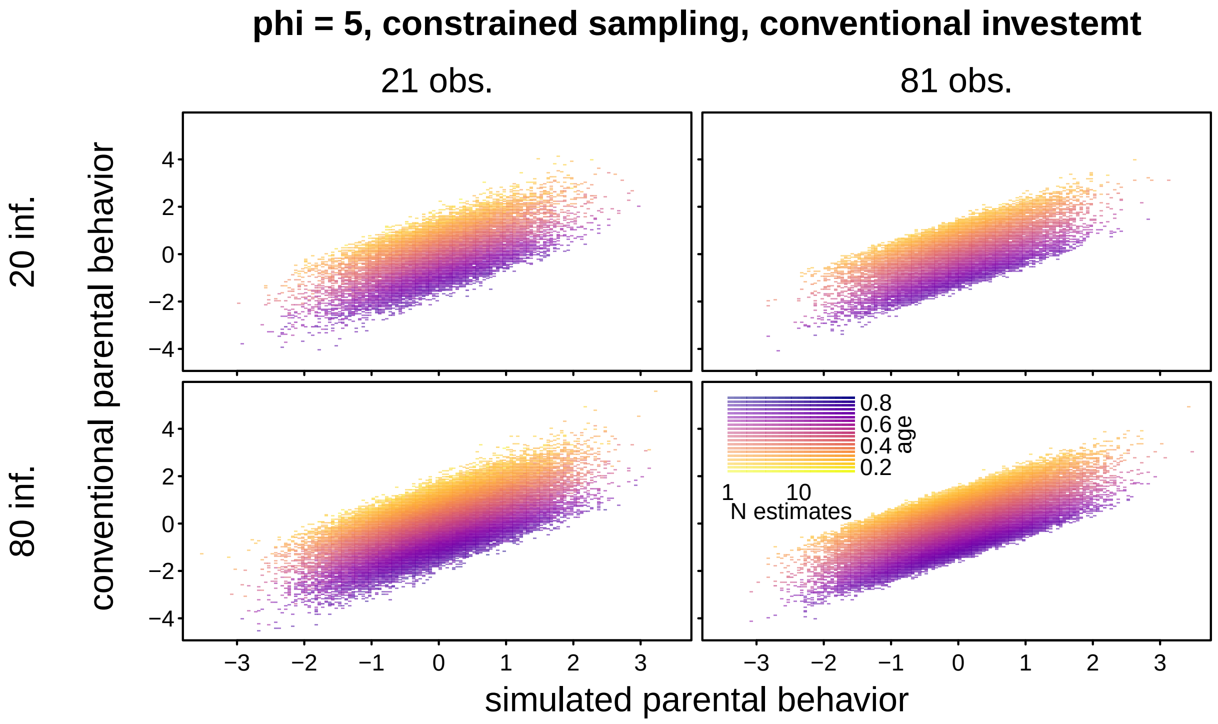


Figure SI 21. Conventional estimates of parent-specific behavior as a function of simulated parent-specific behavior, separately for the different combinations of number parent offspring dyads (rows) and number observations per dyad (columns). Sampling was constrained and the precision parameter *phi* was 5. The color (yellow to blue) codes average offspring age, and saturation codes number estimates per bin.

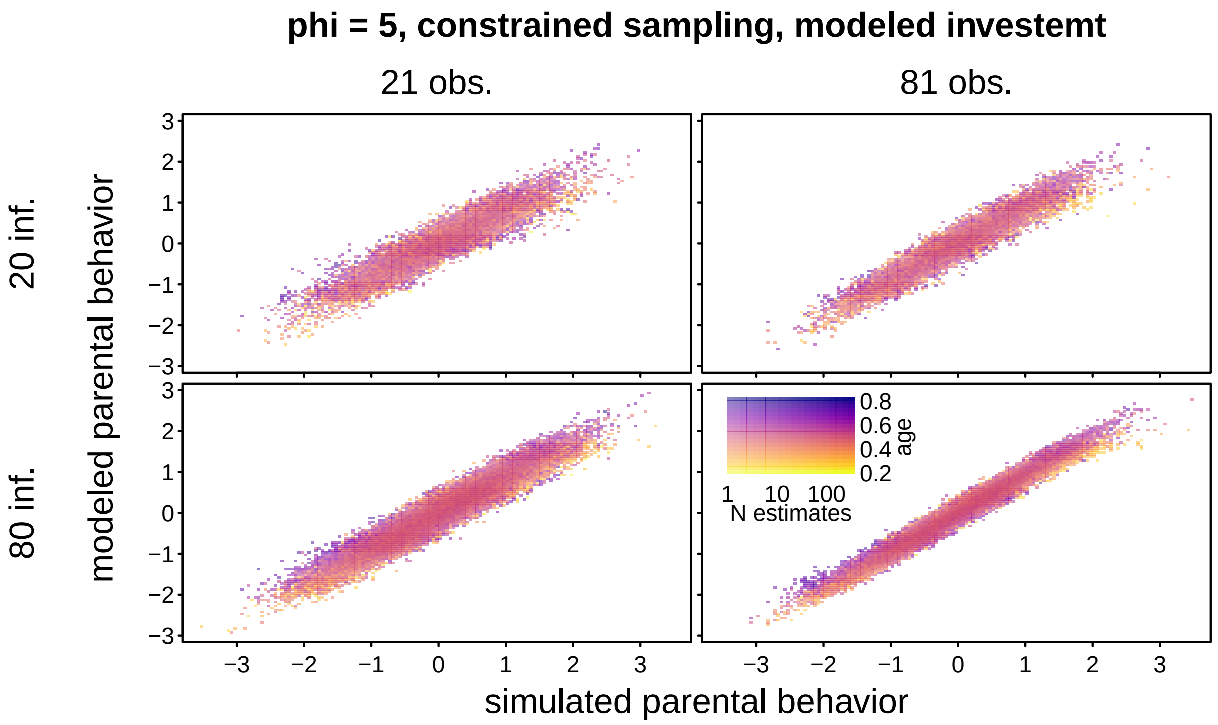


Figure SI 22. Modeled estimates of parent-specific behavior as a function of simulated parent-specific behavior, separately for the different combinations of number parent offspring dyads (rows) and number observations per dyad (columns). Sampling was constrained and the precision parameter *phi* was 5. The color (yellow to blue) codes average offspring age, and saturation codes number estimates per bin.

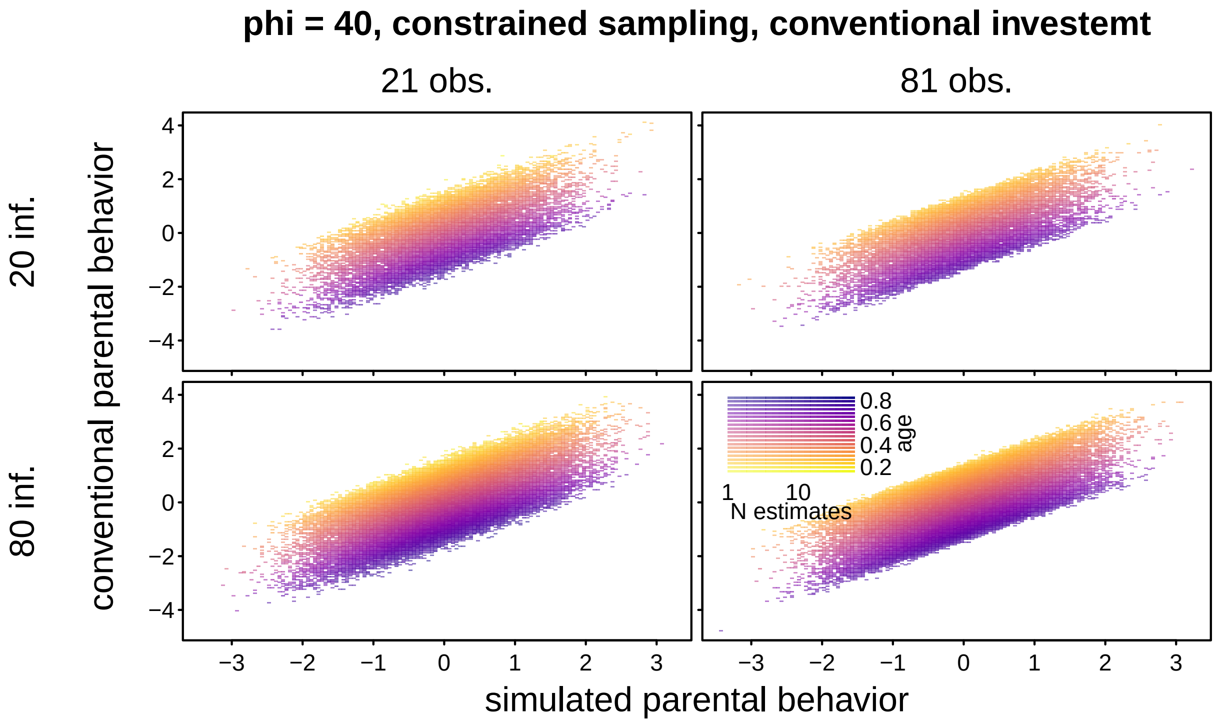


Figure SI 23. Conventional estimates of parent-specific behavior as a function of simulated parent-specific behavior, separately for the different combinations of number parent offspring dyads (rows) and number observations per dyad (columns). Sampling was constrained and the precision parameter *phi* was 40. The color (yellow to blue) codes average offspring age, and saturation codes number estimates per bin.

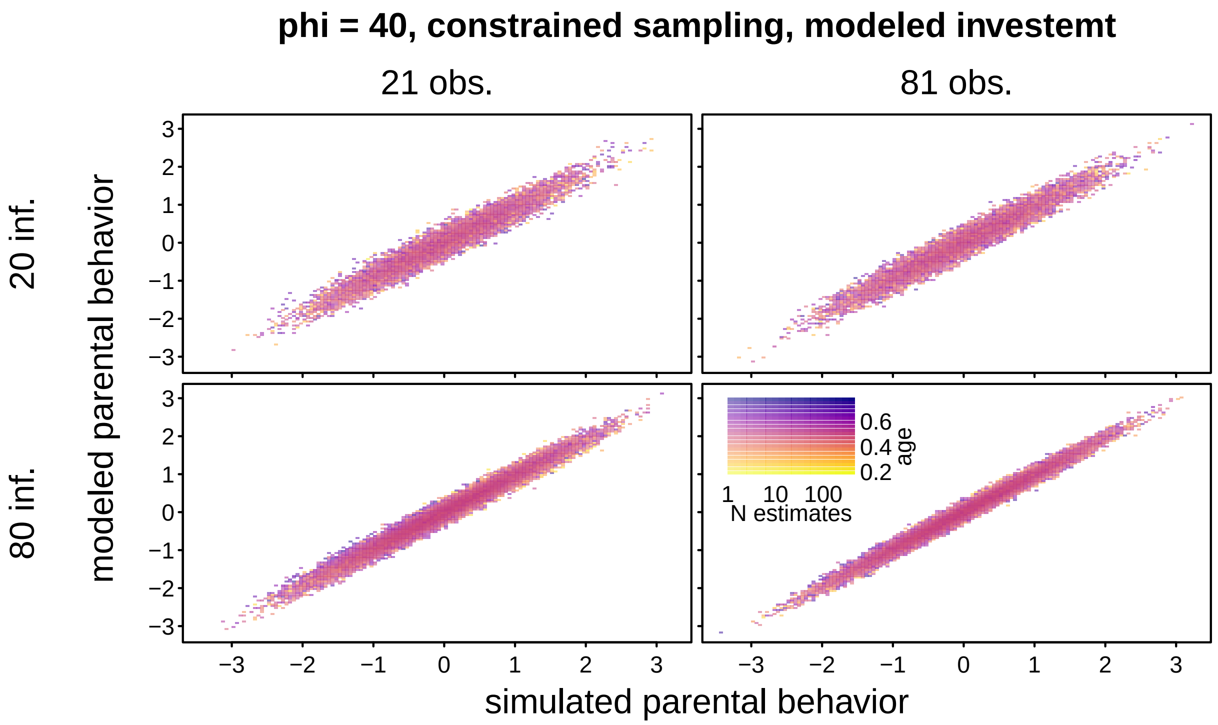


Figure SI 24. Modeled estimates of parent-specific behavior as a function of simulated parent-specific behavior, separately for the different combinations of number parent offspring dyads (rows) and number observations per dyad (columns). Sampling was constrained and the precision parameter *phi* was 40. The color (yellow to blue) codes average offspring age, and saturation codes number estimates per bin.

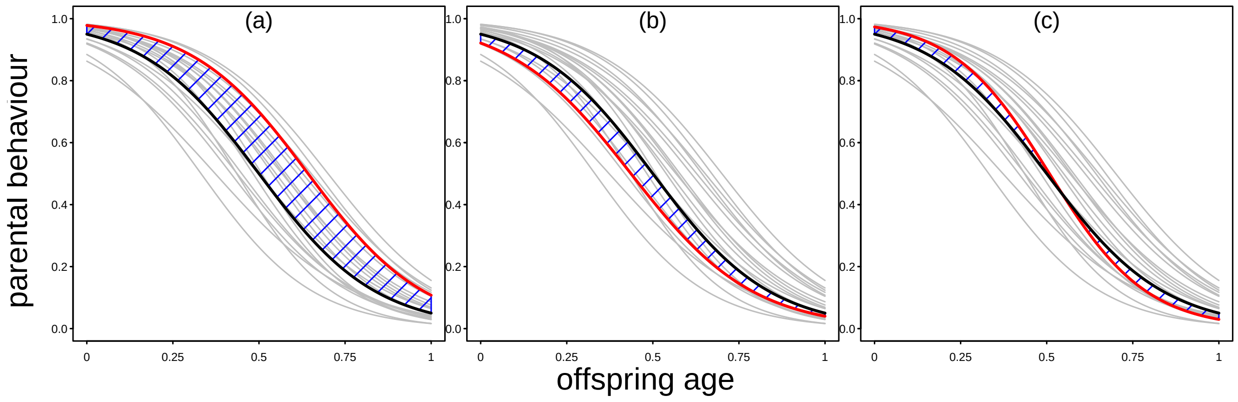


Figure SI 25. Proposed *ad-hoc* solution to estimate parent-specific behavior for a model comprising a random slope of offspring age within parent-offspring ID. Each grey and the red line represent one parent-offspring specific trajectory, and the black line represents the average trajectory (across all parent-offspring dyads). The area shaded in blue depicts the integral between a given parent-offspring specific and the average trajectory. In (a), the highlighted parent shows relatively large behavior throughout and has a relatively large (i.e., positive) integral. In (b), the highlighted parent shows relatively little behavior throughout and has a relatively small (i.e., negative) integral. In (c), the parent shows an behavior above the average parent when its offspring is young and an behavior below average when its offspring is old. Its integral thus comprises a positive and negative section and hence will be about zero.

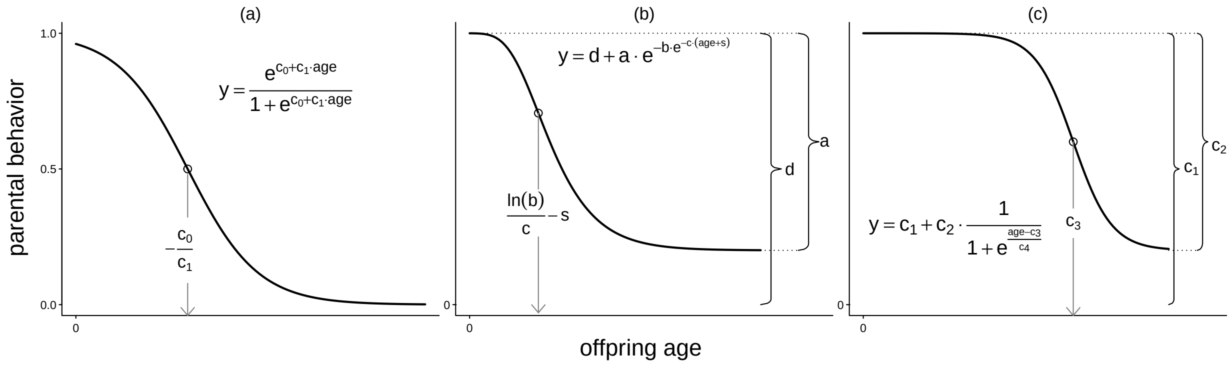


Figure SI 26 . Illustration of three possible age-dependent trajectories of a parental behavior. Depicted are a standard logit-link linear model (a), a modified Gompertz function (b), and a sigmoidal function (c), together with the respective model equation (y=...). Where possible, the interpretation of the parameters of the function is indicated (the parameter $c_4$ in (c) determines the steepness of the trajectory). The open dot shows the location of the inflection point, and the expression laying over the downwards arrow indicates the age at which the parental behavior changes most quickly. Throughout, $e$ represents Euler's number and ln refers to the natural logarithm. It is important to note that these are just three out of a larger number of functions allowing similar trajectories, and that not all offspring age dependent trajectories of a similar principal form can be modeled with any of the three functions depicted here.

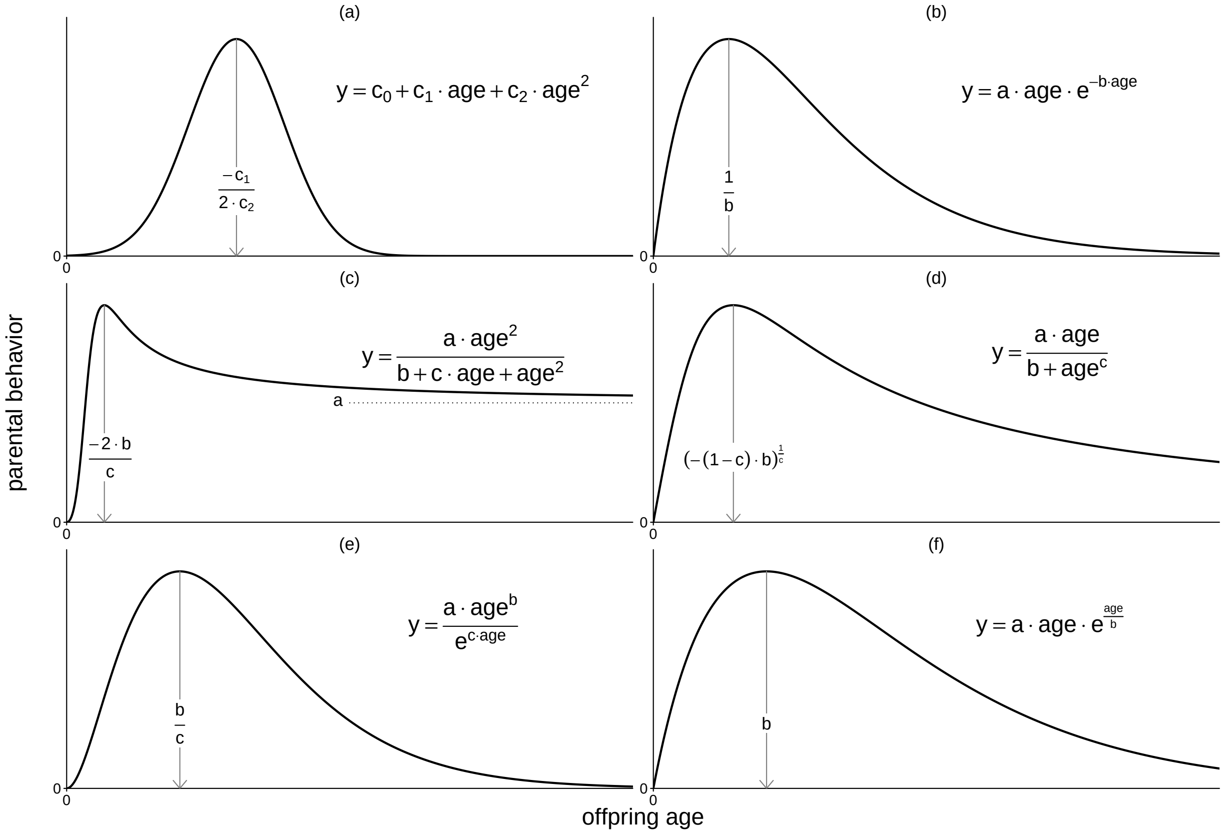


Figure SI 27. Illustration of possible age-dependent trajectories of a parental behavior in which the parental behavior peaks when the offspring is at intermediate age. Depicted are a standard log-link linear model with offspring age linear and squared entered as predictors (a), a Ricker function (b), a Holling type IV function (c), a Sheperd function (d), an incomplete Wood Gamma function (e), and a negative exponential function (f), together with the respective model equation (y=...). The expression laying over the downwards arrow indicates the age at which the parental behavior peaks. Throughout, $e$ represents Euler's number. The Ricker (b), Holling type IV (c), the Sheperd (d), and the negative exponential function (f) are taken from Bolker (2008; modified) and the incomplete Wood Gamma (e) is taken from Marumo et al. (2022; modified). Both the Sheperd (d) and the negative exponential function (f) approach zero as offspring age increases. It is important to note that these are a selection out of a larger number of functions allowing similar trajectories, and that not all offspring age dependent trajectories of a similar principal form can be modeled with any of the functions depicted here.

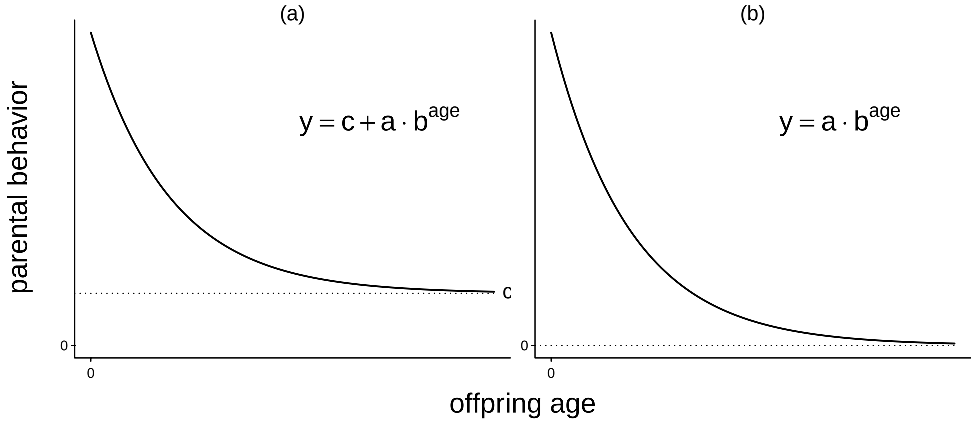


Figure SI 28. Illustration of exponential trajectories of a parental behavior against offspring age, together with the respective model equation (y=...). Note the relationship between offspring age and parental behavior depicted in (b) would be linear when the parental behavior is log-transformed. Practically, however, a log-transformation of the parental behavior might make sense in case of a behavior which is $\geq 0$ and continuously varying such as latencies. But for counts or proportions log-transforming the behavior is not a viable option. For a count response, however, a standard linear model with log-link function is potentially fully appropriate.
